# Pooled CRISPR Inverse PCR sequencing (PCIP-seq): simultaneous sequencing of retroviral insertion points and the integrated provirus with long reads

**DOI:** 10.1101/558130

**Authors:** Maria Artesi, Vincent Hahaut, Basiel Cole, Laurens Lambrechts, Fereshteh Ashrafi, Ambroise Marçais, Olivier Hermine, Philip Griebel, Natasa Arsic, Frank van der Meer, Arsène Burny, Dominique Bron, Elettra Bianchi, Philippe Delvenne, Vincent Bours, Carole Charlier, Michel Georges, Linos Vandekerkhove, Anne Van den Broeke, Keith Durkin

**Author notes:** These authors contributed equally: Maria Artesi, Vincent Hahaut. These authors jointly supervised the work: Anne Van den Broeke, Keith Durkin.

## Abstract

Retroviral infections create a large population of cells, each defined by a unique proviral insertion site. Methods based on short-read high throughput sequencing can identify thousands of insertion sites, but the proviruses within remain unobserved. We have developed Pooled CRISPR Inverse PCR sequencing (PCIP-seq), a method that leverages long reads on the Oxford Nanopore MinION platform to sequence the insertion site and its associated provirus. We have applied the technique to natural infections produced by three exogenous retroviruses, HTLV-1, BLV and HIV-1 as well as endogenous retroviruses in both cattle and sheep. The high efficiency of the method facilitated the identification of tens of thousands of insertion sites in a single sample. We observed thousands of SNPs and dozens of structural variants within proviruses. While initially developed for retroviruses the method has also been successfully extended to DNA extracted from HPV positive PAP smears, where it could assist in identifying viral integrations associated with clonal expansion.

## Introduction

The integration of viral DNA into the host genome is a defining feature of the retroviral life cycle, irreversibly linking provirus and cell. This intimate association facilitates viral persistence and replication in somatic cells, and with integration into germ cells bequeaths the provirus to subsequent generations. Considerable effort has been expended to understand patterns of proviral integration, both from a basic virology stand point, and due to the use of retroviral vectors in gene therapy^1^. The application of next generation sequencing (NGS) over the last ∼10 years has had a dramatic impact on our ability to explore the landscape of retroviral integration for both exogenous and endogenous retroviruses. Methods based on ligation mediated PCR and Illumina sequencing have facilitated the identification of hundreds of thousands of insertion sites in exogenous viruses such as Human T-cell leukemia virus-1 (HTLV-1)^2^ and Human immunodeficiency virus (HIV-1)^3-6^. These techniques have shown that in HTLV-1^2^, Bovine Leukemia Virus (BLV)^7^ and Avian Leukosis Virus (ALV)^8^ integration sites are not random, pointing to clonal selection. In HIV-1 it has also become apparent that provirus integration can drive clonal expansion^3,4,6,9^, magnifying the HIV-1 reservoir and placing a major road block in the way of a complete cure.

Current methods based on short-read sequencing identify the insertion point, but the provirus itself is largely unexplored. Whether variation in the provirus influences the fate of the clone remains difficult to investigate. Using long range PCR it has been shown that proviruses in HTLV-1 induced Adult T-cell leukemia (ATL) are frequently (∼45%) defective^10^, although the abundance of defective proviruses within asymptomatic HTLV-1 carriers has not been systematically investigated. Recently, there has been a concerted effort to better understand the structure of HIV-1 proviruses in the latent reservoir. Methods such as Full-Length Individual Proviral Sequencing (FLIPS) have been developed to identify functional proviruses^11^ but without identifying the provirus integration site. More recently matched integration site and proviral sequencing (MIP-Seq) has allowed the sequence of individual proviruses to be linked to integration site in the genome^6^. However, this method relies on whole genome amplification of isolated HIV-1 genomes, with separate reactions to identify the integration site and sequence the associated provirus^6^. As a result, this method is quite labor intensive limiting the number of proviruses one can reasonably interrogate.

Retroviruses are primarily associated with the diseases they provoke through the infection of somatic cells. Over the course of evolutionary time they have also played a major role in shaping the genome. Retroviral invasion of the germ line has occurred multiple times, resulting in the remarkable fact that endogenous retrovirus (ERV)-like elements comprise a larger proportion of the human genome (8%) than protein coding sequences (∼1.5%)^12^. With the availability of multiple vertebrate genome assemblies, much of the focus has been on comparison of ERVs between species. However, single genomes represent a fraction of the variation within a species, prompting some to take a population approach to investigate ERV– host genome variation^13^. While capable of identifying polymorphic ERVs in the population, approaches relying on conventional paired-end libraries and short reads cannot capture the sequence of the provirus beyond the first few hundred bases of the proviral long terminal repeat (LTR), leaving the variation within uncharted.

In contrast to retroviruses, papillomaviruses do not integrate into the host genome as part of their lifecycle. Human papillomavirus (HPV) is usually present in the cell as a multi copy circular episome (∼8kb in size), however in a small fraction of infections, it can integrate into the host genome leading to the dysregulation of the viral oncogenes E6 and E7^14^. Genome wide profiling of HPV integration sites via capture probes and Illumina sequencing has also identified hotspots of integration indicating that disruption of host genes may also play a role in driving clonal expansion^15^. As a consequence, HPV integration is a risk factor for the development of cervical carcinoma^16^, however its study is hampered by the unpredictability of the breakpoint sites in the integrated HPV genome. This limits the applicability of approaches based on ligation mediated PCR and short read sequencing.

The application of NGS as well as Sanger sequencing before, has had a large impact on our understanding of both exogenous and endogenous proviruses. The development of long-read sequencing, linked-read technologies and associated computational tools^17^ have the potential to explore questions inaccessible to short reads. Groups investigating Long interspersed nuclear elements-1 (LINE-1) insertions^18^ and the koala retrovirus, KoRV^19^ have highlighted this potential and described techniques utilizing the Oxford Nanopore and PacBio platforms, to investigate insertion sites and retroelement structure.

To more fully exploit the potential of long reads we developed Pooled CRISPR Inverse PCR sequencing (PCIP-seq), a method that leverages selective cleavage of circularized DNA fragments carrying proviral DNA with a pool of CRISPR guide RNAs, followed by inverse long-range PCR and multiplexed sequencing on the Oxford Nanopore MinION platform. Using this approach, we can now simultaneously identify the integration site and track clone abundance while also sequencing the provirus inserted at that position. We have successfully applied the technique to the retroviruses HTLV-1, HIV-1 and BLV, endogenous retroviruses in cattle and sheep as well as HPV18.

## Results

### Overview of PCIP-seq (Pooled CRISPR Inverse PCR-sequencing)

The genome size of the viruses targeted ranged from 6.8 to 9.7kb, therefore we chose to shear the DNA to ∼8kb in length. In most cases this creates two fragments for each provirus, one containing the 5’ end with host DNA upstream of the insertion site and the second with the 3’ end and downstream host DNA. Depending on the shear site the amount of host and proviral DNA in each fragment will vary (Fig. 1a). To facilitate identification of the provirus insertion site via inverse PCR we carry out intramolecular ligation, followed by digestion of the remaining linear DNA. To selectively linearize the circular DNA containing proviral sequences (this helps increase PCR efficiency), regions adjacent to the 5’ and 3’ LTRs in the provirus are targeted for CRISPR mediated cleavage. We sought a balance between ensuring that the majority of the reads contained part of the flanking DNA (for clone identification) while also generating sufficient reads extending into the midpoint of the provirus. We found that using a pool of CRISPR guides for each region increased the efficiency and by multiplexing the guide pools and PCR primers for the 5’ and 3’ ends we could generate coverage for the majority of a clonally expanded provirus in a single reaction (Fig. 1b). The multiplexed pool of guides and primers leaves coverage gaps in the regions flanked by the primers. To address these coverage gaps we designed a second set of guides and primers. Following separate CRISPR cleavage and PCR amplification the products of these two sets of guides and primers were combined for sequencing (Fig. 1c). This approach ensured that the complete provirus was sequenced (Fig. 1d).

**Figure 1.**
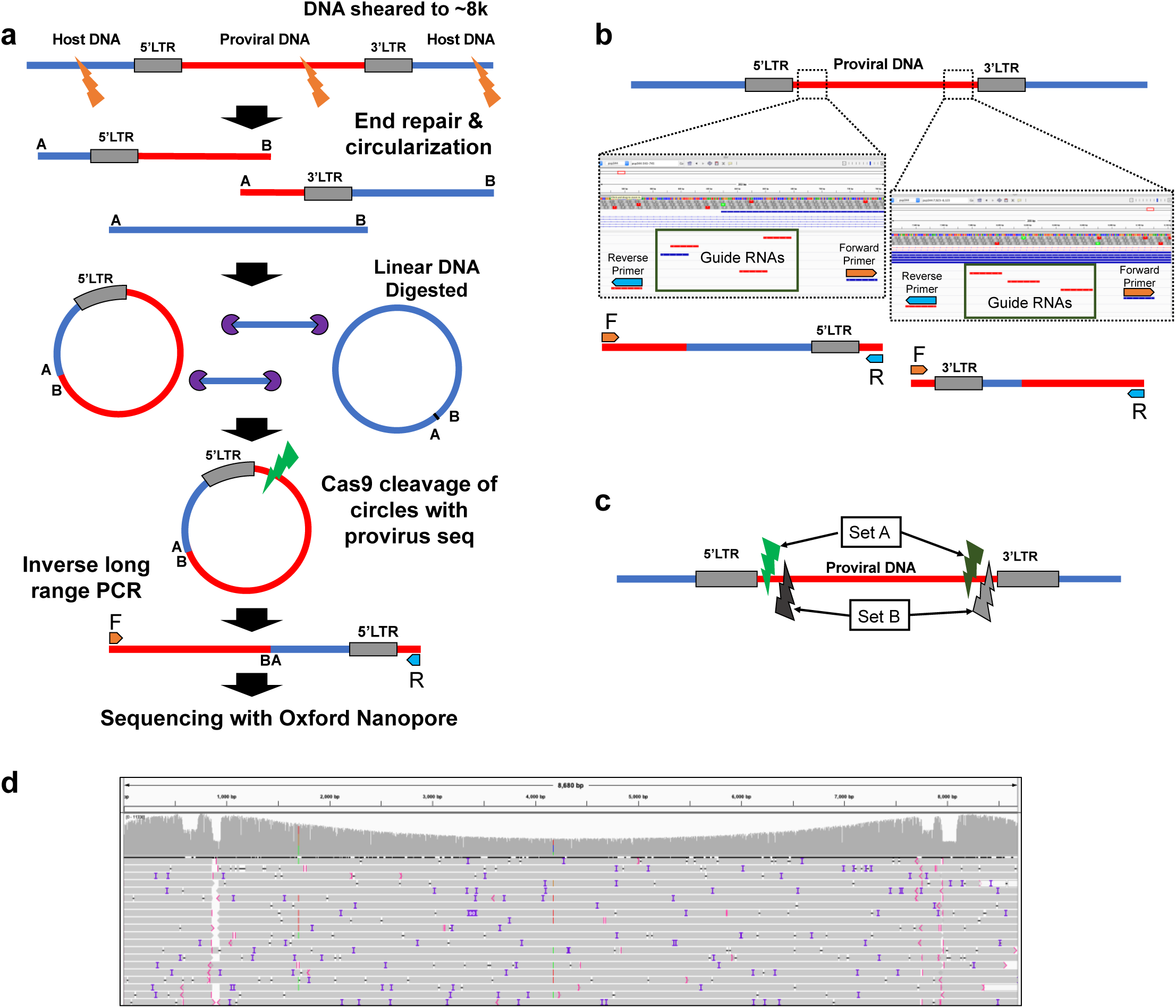
Overview of the PCIP-seq method **(a)** Simplified outline of method **(b)** A pool of CRISPR guide-RNAs targets each region, the region is flanked by PCR primers. Guides and primers adjacent to 5’ & 3’ LTRs are multiplexed. **(c)** As the region between the PCR primers is not sequenced we created two sets of guides and primers. Following circularization, the sample is split, with CRISPR mediated cleavage and PCR occurring separately for each set. After PCR the products of the two sets of guides and primers are combined for sequencing. **(d)** Screen shot from the Integrative Genomics Viewer (IGV) showing a small fraction of the resultant reads (grey bars) mapped to the provirus, coverage is shown on top, coverage drops close to the 5’ and 3’ ends are regions flanked by primers.

### Identifying genomic insertions and internal variants in HTLV-1

Adult T-cell leukemia (ATL) is an aggressive cancer induced by HTLV-1. It is generally characterized by the presence of a single dominant malignant clone, identifiable by a unique proviral integration site. We and others have developed methods based on ligation mediated PCR and Illumina sequencing to simultaneously identify integration sites and determine the abundance of the corresponding clones ^2,7^. We initially applied PCIP-seq to two HTLV-1 induced cases of ATL, both previously analyzed with our Illumina based method (ATL2^7^ & ATL100^20^). In ATL100 both methods identify a single dominant clone, with >95% of the reads mapping to a single insertion site on chr18 (Fig. 2a, 2b & Table1). Using the integration site information, we extracted the PCIP-seq hybrid reads spanning the provirus/host insertion site, uncovering a ∼3,600bp deletion within the provirus (Fig. 2c).

**Table 1.** Number of insertion sites (IS) identified via PCIP-seq. Chimeric reads = reads containing host and viral DNA. Largest clone % = insertion site with highest number of reads in that sample. PVL = Proviral Load. (Percentage cells carrying a single copy of integrated provirus or number proviral copies per 100 cells).

| Sample name | Virus | Host | PVL | Template μg | raw reads | Chimeric reads (%) | Pure Host / Pure Viral reads | Insertion sites | Largest clone (%) |
| --- | --- | --- | --- | --- | --- | --- | --- | --- | --- |
| ATL2 | HTLV-1 | HSA | nd | 4 | 81,219 | 68.21 | 0.0037 / 31.8 | 160 | 49.5 |
| ATL100 | HTLV-1 | HSA | 106 | 4 | 4,838 | 64.14 | 9.16 / 26.7 | 13 | 89.624 |
| 233 | BLV | OAR | 78.3 | 7 | 524,698 | 53.4 | 0.04 / 46.53 | 5311 | 5.22 |
| 221 (022016) | BLV | OAR | 63 | 4 | 180,276 | 67.14 | 3.59 / 29.27 | 8023 | 0.625 |
| 221 (032014) | BLV | OAR | 16 | 4 | 32,266 | 68.69 | 0.11 / 31.20 | 5374 | 0.279 |
| 220 | BLV | OAR | 3.8 | 2 | 44,876 | 67.38 | 0 / 32.62 | 1352 | 3.55 |
| 1439 | BLV | BosT | 45 | 3 | 181,055 | 70.52 | 0.19 / 29.29 | 5773 | 1.17 |
| 560 | BLV | BosT | 0.644 | 1 | 6,802 | 69.83 | 1.12 / 29.06 | 172 | 4.59 |
| 1053 | BLV | BosT | 23.5 | 6 | 367,454 | 72.13 | 0.04 / 27.83 | 17903 | 0.353 |
| HIV_U1 | HIV-1 | HSA | 200 | 2 | 94,086 | 54.66 | 2.75 / 42.59 | 728 | 47.2 |
| Jurkat U1-0.1 | HIV-1 | HSA | 0.2 | 5 | 252,913 | 43.33 | 0.04 / 56.62 | 4 | 71.7 |
| Jurkat U1-0.01 | HIV-1 | HSA | 0.02 | 5 | 234,421 | 43.33 | 0.04 / 56.52 | 2 | 90.2 |
| Jurkat neg | HIV-1 | HSA | 0 | 5 | 12,137 | 0 | 100 / 0 | 0 | 0 |
| 02006 | HIV-1 | HSA | 0.46 | 12 | 240,641 | 51.63 | 1.10 / 47.27 | 159 | 7.82 |
| 06042 | HIV-1 | HSA | 0.56 | 8 | 226,685 | 21.18 | 0.41 / 78.41 | 74 | 4.77 |
| HPV18_PX | HPV18 | HAS | nd | 4 | 180,550 | 21.36 | 0.29 / 78.35 | 55 | nd |
| HPV18_PY | HPV18 | HAS | nd | 4 | 82,807 | 0.09 | 0.05 / 99.86 | 19 | nd |

**Figure 2.**
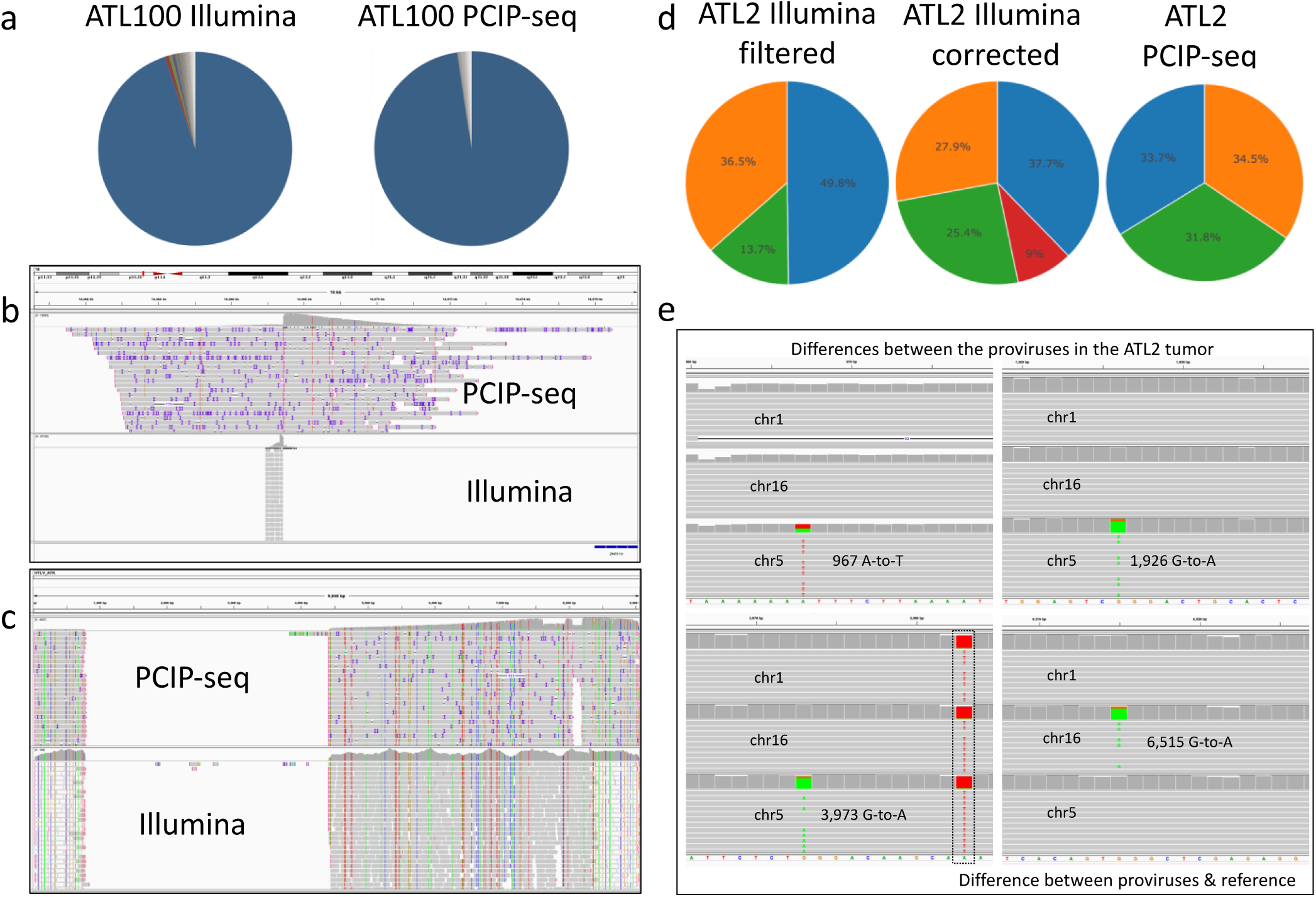
PCIP-seq applied to ATL **(a)** In ATL100 both Illumina and Nanopore based methods show a single predominant insertion site **(b)** Screen shot from IGV shows a ∼16kb window with the provirus insertion site in the tumor clone identified via PCIP-seq and ligation mediated PCR with Illumina sequencing **(c)** PCIP-seq reads in IGV show a ∼3,600bp deletion in the provirus, confirmed via long range PCR and Illumina sequencing. **(d)** The ATL2 tumor clone contains three proviruses (named according to chromosome inserted into), the provirus on chr1 inserted into a repetitive element (LTR) and short reads generated from host DNA flanking the insertion site map to multiple positions in the genome. Filtering out multi-mapping reads causes an underestimation of the abundance of this insertion site (13.6 %), this can be partially corrected by retaining multi-mapping reads at this position (25.4 %). However, that approach can cause the potentially spurious inflation of other integration sites (red slice 9%). The long PCIP-seq reads can span repetitive elements and produce even coverage for each provirus without correction. **(e)** Screen shot from IGV shows representative reads coming from the three proviruses at positions where four *de novo* mutations were observed.

In the case of ATL2, PCIP-seq showed three major proviruses located on chr5, chr16 and chr1, each responsible for ∼33% of the HTLV-1/host hybrid reads. We had previously established that these three proviruses are in a single clone via examination of the T-cell receptor gene rearrangement^7^. However, it is interesting to note that this was not initially obvious using our Illumina based method as the proviral insertion site on chr1 falls within a repetitive element (LTR) causing many of the reads to map to multiple regions in the genome. If multi mapping reads are filtered out, the chr1 insertion site accounted for 13.7% of the remaining reads, while retaining multi mapping produces values closer to reality (25.4%). In contrast the long reads from PCIP-seq allow unambiguous mapping and closely matched the expected 33% for each insertion site (Fig. 2d), highlighting the advantage long reads have in repetitive regions. Looking at the three proviruses, proviral reads revealed all to be full length. Three de novo mutations were observed in one provirus and a single de novo mutation was identified in the second (Fig. 2e).

### Insertion sites identified in samples with multiple clones of low abundance

The samples utilized above represent a best-case scenario, with ~100% of cells infected and a small number of major clones. We next applied PCIP-seq to four samples from BLV infected sheep (experimental infection^21^) and three cattle (natural infection) to explore its performance on polyclonal and low proviral load (PVL) samples and compared PCIP-seq to our previously published Illumina method^7^. PCIP-seq revealed all samples to be highly polyclonal (Supplementary Fig. 1 and Table 1) with the number of unique insertion sites identified varying from 172 in the bovine sample 560 (1μg template, PVL 0.644%) to 17,903 in bovine sample 1053 (6μg template, PVL 23.5%). In general, PCIP-seq identified more insertion sites, using less input DNA than our Illumina based method (Supplementary Table 1). Comparison of the results showed a significant overlap between the two methods. When we consider insertion sites supported by more than three reads in both methods (larger clones, more likely to be present in both samples), in the majority of cases >50% of the insertion sites identified in the Illumina data were also observed via PCIP-seq (Supplementary Table 1). These results show the utility of PCIP-seq for insertion site identification, especially considering the advantages long reads have in repetitive regions of the genome.

### Identifying SNPs in BLV proviruses

Portions of the proviruses with more than ten supporting reads (PCR duplicates removed) were examined for SNPs with LoFreq^22^. For the four sheep samples, the variants were called relative to the pBLV344 provirus (used to infect the animals). For the bovine samples 1439 and 1053 custom consensus BLV sequences were generated for each and the variants were called in relation to the appropriate reference (SNPs were not called in 560). Across all the samples 3,209 proviruses were examined, 934 SNPs were called and 680 (21%) of the proviruses carried one or more SNPs (Supplementary Table 2). We validated 10 BLV SNPs in the ovine samples and 15 in the bovine via clone specific long-range PCR and Illumina sequencing (Supplementary Fig. 2). For Ovine 221, which was sequenced twice over a two-year interval, we identified and validated three instances where the same SNP and provirus were observed at both time points (Supplementary Fig. 2). We noted a small number of positions in the BLV provirus prone to erroneous SNP calls. By comparing allele frequencies from bulk Illumina and Nanopore data these problematic positions could be identified (Supplementary Fig. 3a).

Approximately half of the SNPs (47.1% sheep, 51.6% cattle) were found in multiple proviruses. Generally, SNPs found at the same position in multiple proviruses were concentrated in a single individual, indicating their presence in a founder provirus or via a mutation in the very early rounds of viral replication (Supplementary Fig. 3b). Alternatively, a variant may also rise in frequency due to increased fitness of clones carrying a mutation in that position. In this instance, we would expect to see the same position mutated in multiple individuals. One potential example is found in the first base of codon 303 (position 8155) of the viral protein Tax, a potent viral transactivator, stimulator of cellular proliferation and highly immunogenic^23^. A variant was observed at this position in five proviruses for sheep 233 and three for sheep 221 as well as one provirus from bovine 1439 (Fig. 3 a). Using less stringent criteria for the inclusion of a proviral region (>10 reads, not filtered for PCR duplicates) we found 34 proviruses in the ovine and 3 in the bovine carrying a variant in this position. The majority of the variants observed were G-to-A transitions (results in E-to-K amino acid change), however we also observed G-to-T (E-to-STOP) and G-to-C (E-to-Q) transversions. It has been previously shown that the G-to-A mutation abolishes the Tax proteins transactivator activity^23,24^. The repeated selection of variants at this specific position suggests that they reduce viral protein recognition by the immune system, while preserving the Tax proteins other proliferative properties.

**Figure 3.**
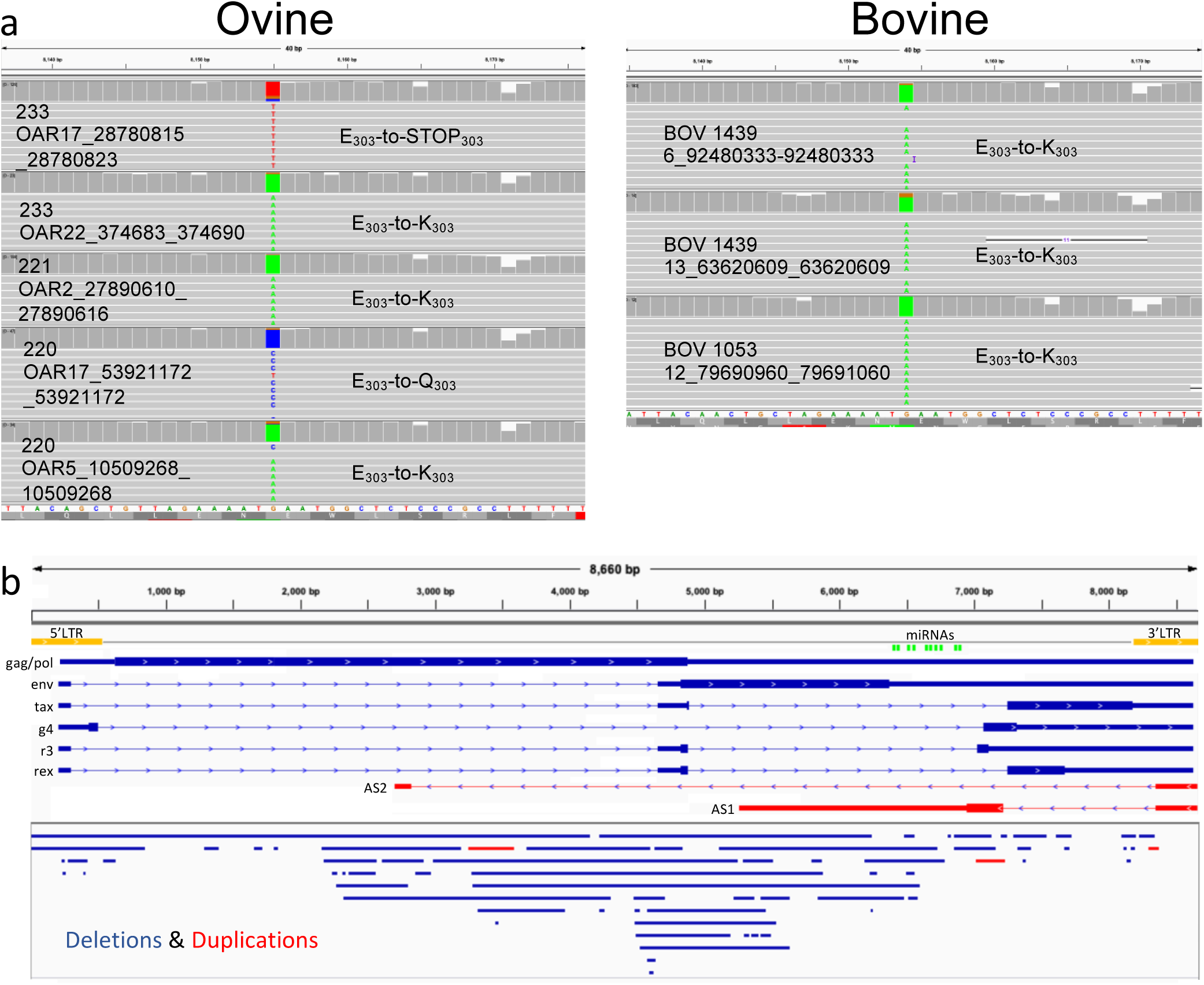
**(a)** Screen shot from IGV shows representative reads from a subset of the clones from each BLV-infected animal with a mutation in the first base of codon 303 in the viral protein Tax. **(b)** Structural variants observed in the BLV provirus. BLV sense and antisense transcripts are shown on top. Deletions (blue bars) and duplications (red bars) observed in the BLV provirus from both ovine and bovine samples are shown below.

Patterns of provirus-wide APOBEC3G^25^ induced hypermutation (G-to-A) were not observed in BLV. However, three proviruses (two from sheep 233 and one in bovine 1053) showed seven or more A-to-G transitions, confined to a ∼70bp window in the first half of the U3 portion of the 3’LTR (Supplementary Fig. 4). The pattern of mutation, as well as their location in the provirus suggests the action of RNA adenosine deaminases 1 (ADAR1)^26,27^.

### PCIP-seq identifies BLV structural variants in multiple clones

Proviruses were also examined for structural variants (SVs) using a custom script and via visualization in IGV (see methods). Between the sheep and bovine samples, we identified 66 deletions and 3 tandem duplications, with sizes ranging from 15bp to 4,152bp, with a median of 113bp (Supplementary Table 3). We validated 14 of these via clone specific PCR (Supplementary Fig. 5). As seen in Fig. 3b SVs were found throughout the majority of the provirus, encompassing the highly expressed microRNAs^28^ as well as the second exon of the constitutively expressed antisense transcript *AS1*^29^. Only two small regions at the 3’ end lacked any SVs. More proviruses will need to be examined to see if this pattern holds, but these results again suggest the importance of the 3’LTR and its previously reported interactions with adjacent host genes^7^.

### Identifying HIV-1 integration sites and the associated provirus

Despite the effectiveness of combination antiretroviral therapy (ART) in suppressing HIV-1 replication, cART is not capable of eliminating latently infected cells, ensuring a viral rebound if cART is suspended^30^. This HIV-1 reservoir represents a major obstacle to a HIV cure^31^ making its exploration a priority. However, this task is complicated by its elusiveness, with only ∼0.1% of CD4^+^ T cells carrying integrated HIV-1 DNA^32^. To see if PCIP-seq could be applied to these extremely low proviral loads we initially carried out dilution experiments using U1^33^, a HIV-1 cell line containing replication competent proviruses^34^. PCIP-seq on undiluted U1 DNA found the major insertion sites on chr2 and chrX (accounting for 47% & 41% of the hybrid reads respectively) and identified the previously reported variants that disrupt Tat function^35^ in both proviruses (Supplementary Fig. 6a). In addition to the two major proviruses we identified an additional ∼700 low abundance insertion sites (Table 1) including one on chr19 (0.8%) reported by Symons et al 2017^34^ that is actually a product of recombination between the major chrX and chr2 proviruses (Supplementary Fig. 6b). We then serially diluted U1 DNA in Jurkat cell line DNA. PCIP-seq was carried out with 5 μg of template DNA where U1 represents 0.1% and 0.01% of the total DNA. We also processed 5 μg of Jurkat DNA in parallel as a negative control. We were able to detect the major proviruses on chr2 and chrX in both dilutions (Supplementary Fig. 7a & Table 1). No reads mapping to HIV-1 were observed in the negative control (Supplementary Fig. 7b & Table 1).

We next carried out PCIP-seq on DNA extracted from the CD4^+^ T cells of two HIV-1 infected patients (06042 & 02006) on long term cART (Supplementary Table 4). Using 8 μg of template DNA we identified 74 unique integration sites in 06042. In 02006 using 12 μg template DNA we identified 159 (Fig. 4 & Supplementary Table. 5). We validated the integration site of 5 proviruses using clone specific PCR (Supplementary Table 5). In the majority of the integration sites only a subset of the associated provirus is sequenced, however it was still possible to identify 12 proviruses from 06042 and 52 in 02006 with large deletions (Supplementary Fig. 8a & Supplementary Table 5). In 02006 we found four clonally expanded full-length proviruses with reads covering the entire provirus (Supplementary Table 5). Of these three had sufficient coverage to generate a consensus sequence. One contained a ∼115 bp deletion just upstream of *gag*, disrupting the packaging signal (Ψ) (Supplementary Fig. 8b). The remaining two appeared to be intact. One maps to a segmentally duplicated region just below the centromere on chr10, while the other has flanking sequence that matches the satellite repeats of the centromeres of chr13, chr14, chr21 and chr22. Both patients had four integration sites in intron 1 of *STAT5B*, all were in the same transcriptional orientation as *STAT5B* (Fig. 4). An enrichment of HIV-1 integrations in this region has previously been reported^3,4,6^, with recent work showing them to cause insertional activation of *STAT5B*, which favours T regulatory cell persistence^36^.

**Figure 4.**
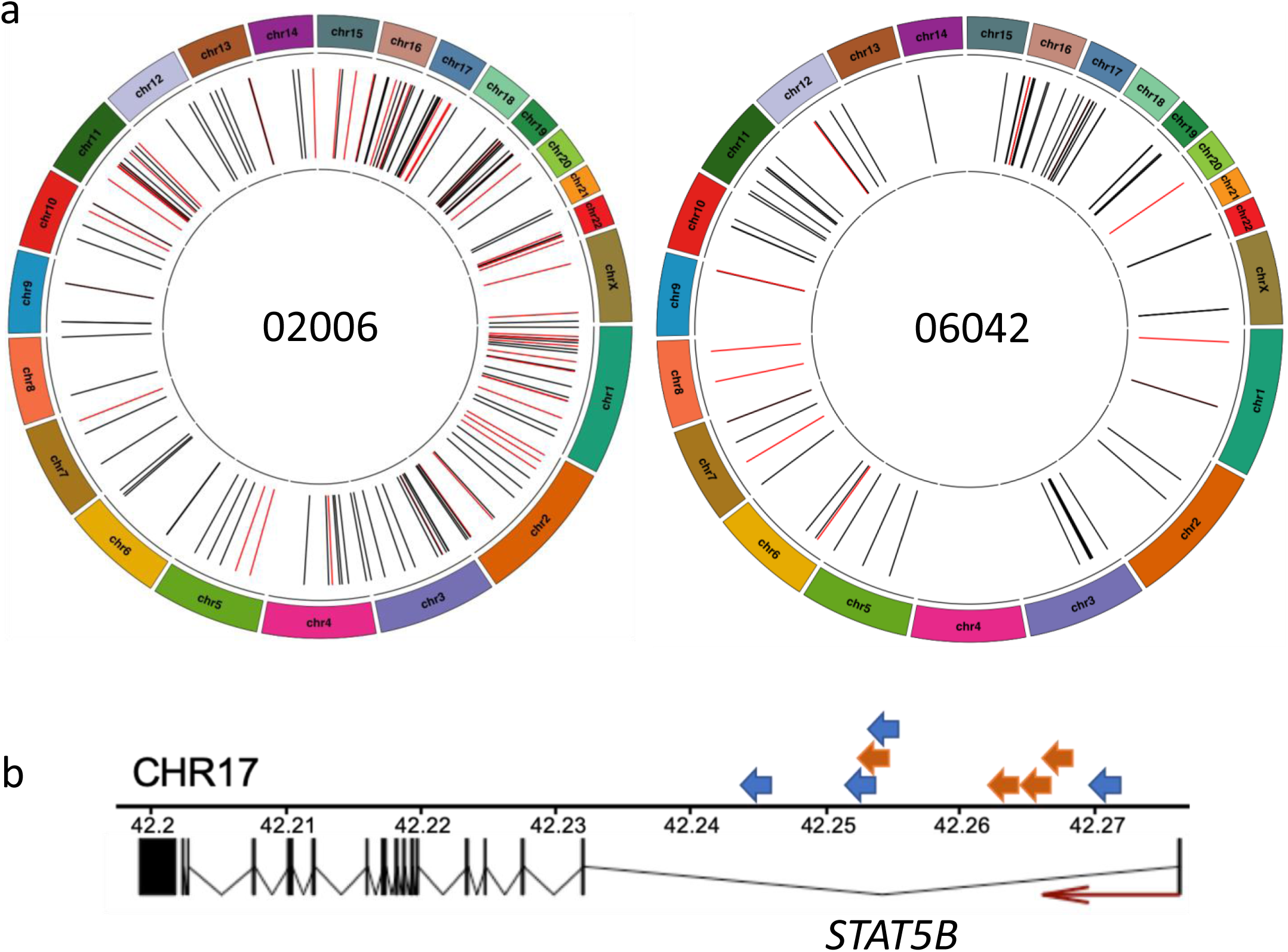
**(a)** HIV-1 proviral integration sites identified by PCIP-seq in two HIV-1 patients (02006 & 06042). Black lines represent integration sites where the portion of the provirus sequence shows no evidence of a large deletion, red lines indicate sites where a large deletion was observed in the provirus. **(b)** A hotspot of proviral integration in intron 1 of *STAT5B.* Arrows represent individual proviruses (02006=blue, 06042=orange) and direction indicates the orientation of the provirus. All proviruses have the same transcriptional orientation as *STAT5B.*

### Identifying full-length and polymorphic endogenous retroviruses in cattle and sheep

ERVs in the genome can be present as full length, complete provirus, or more commonly as solo-LTRs, the products of non-allelic recombination^37^. At the current time conventional short read sequencing, using targeted or whole genome approaches, cannot distinguish between the two classes. Examining full length ERVs would provide a more complete picture of ERV variation, while also revealing which elements can produce *de novo* ERV insertions. As PCIP-seq targets inside the provirus we can preferentially amplify full length ERVs, opening this type of ERV to study in larger numbers of individuals. As a proof of concept we targeted the class II bovine endogenous retrovirus BERVK2, known to be transcribed in the bovine placenta^38^. We applied the technique to three cattle, of which one (10201e6) was a Holstein suffering from cholesterol deficiency, an autosomal recessive genetic defect recently ascribed to the insertion of a 1.3kb LTR in the *APOB* gene^39^. PCIP-seq clearly identified the *APOB* ERV insertion in 10201e6 and in contrast to previous reports^39^ shows it to be a full-length element (Supplementary Fig. 9). We identified a total of 67 ERVs (Fig. 5), with 8 present in all three samples (Supplementary Table 6). We validated three ERVs via long range PCR and Illumina sequencing (Supplementary Fig. 10). We did not find any with an identical sequence to the *APOB* ERV, although the ERV BTA3_115.3 has an identical LTR sequence, highlighting that the sequence of the LTR cannot be used to infer the complete sequence of the ERV (Supplementary Fig. 11).

**Figure 5.**
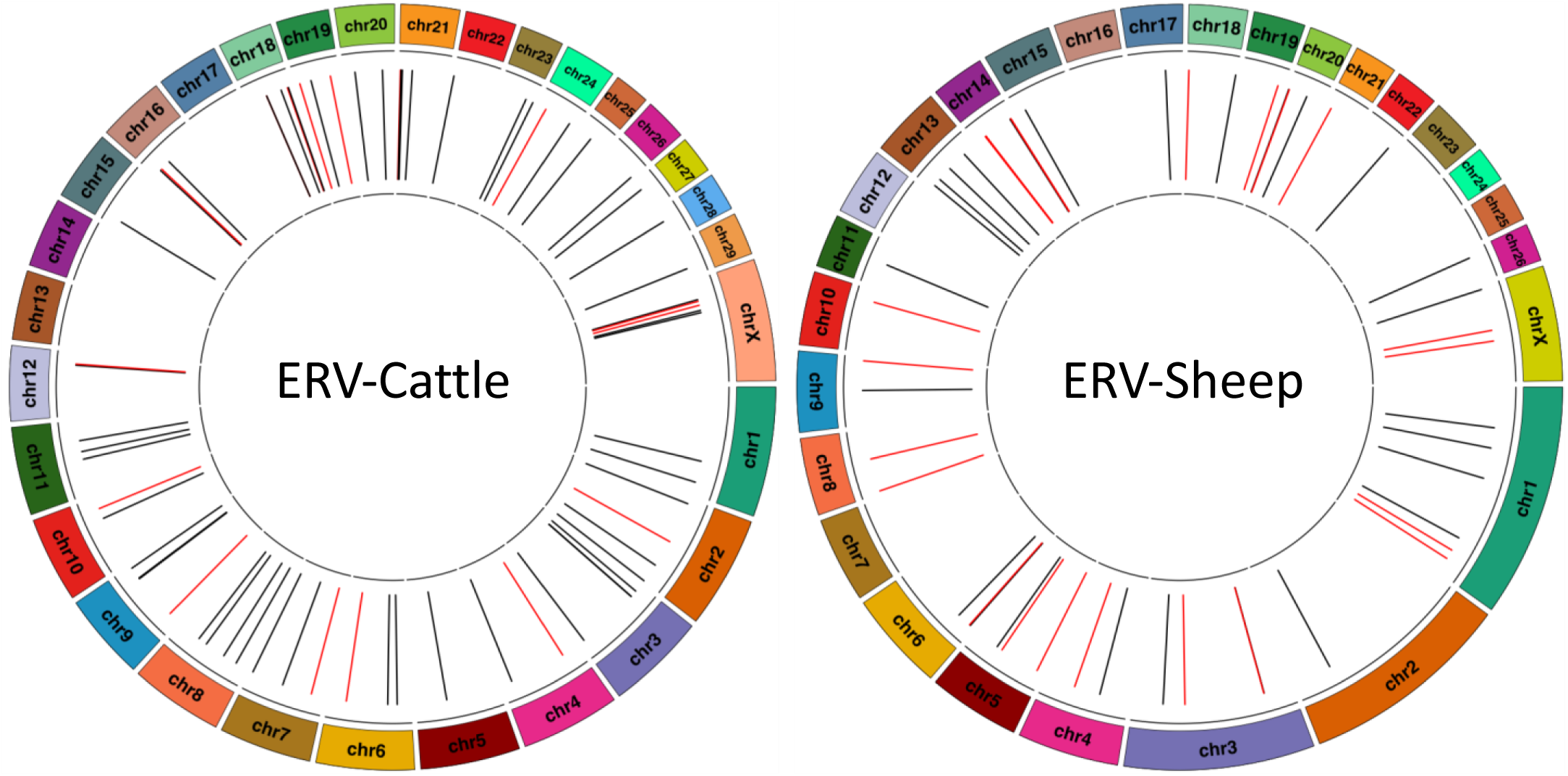
Location of endogenous retroviruses identified by PCIP-seq in cattle and sheep genomes. Based on three cattle and two sheep. Black lines represent full length proviruses, red contain large deletions.

We also adapted PCIP-seq to amplify the Ovine endogenous retrovirus Jaagsiekte sheep retrovirus (enJSRV), a model for retrovirus-host co-evolution^40^. Using two sheep (220 & 221) as template we identified a total of 48 enJSRV proviruses (Fig. 5), (33 in 220 and 38 in 221, with 22 common to both) and of these ∼54% were full length (Supplementary Table 7). We validated seven proviruses via long-range PCR and Illumina sequencing (Supplementary Fig. 12).

### Extending PCIP-seq to human papillomaviruses (HPV)

The majority of HPV infections clear or are suppressed within 1–2 years^41^, however a minority evolve into cancer, and these are generally associated with integration of the virus into the host genome. This integration into the host genome is not part of the viral lifecycle and the breakpoint in the viral genome can occur at any point across is 8kb circular genome^16^. As a consequence the part of the viral genome found at the virus host breakpoint varies considerably, making the identifying of integration sites difficult using existing approaches^16^. The long reads employed by PCIP-seq mean that even when the breakpoint is a number of kb away from the position targeted by primers we should still capture the integration site. As a proof of concept, we applied PCIP-seq to two HPV18 positive cases, (HPV18_PX and HPV18_PY) using 4 μg of DNA extracted from Pap smear material. We identified 55 integration sites in HPV18_PX and 19 integration sites in HPV18_PY (Supplementary Table 8). In HPV18_PY the vast majority of the reads only contained HPV sequences, the integration sites identified were defined by single reads, suggesting little or no clonal expansion (Table 1). In HPV18_PX most integration sites were again defined by a single read, however there were some exceptions (Supplementary Table 8). The most striking of these was a cluster of what appeared to be three integration sites located within the region chr3:52477576-52564190 (Fig. 6a). The unusual pattern of read coverage combined with the close proximity of the virus-host breakpoints indicated that these three integration sites were connected. Long range PCR with primers spanning positions α-β and α-γ, showed that a genomic rearrangement had occurred in this clonally expanded cell (Fig. 6a). Regions α and β are adjacent to one another with HPV integrated between, however PCR also showed regions α and γ to be adjacent to one another, again with the HPV genome integrated between (Fig. 6b). The sequence of the virus found between α-β looks to be derived from the α-γ virus as it shares a breakpoint and is slightly shorter (Fig. 6b). This complex arrangement suggests that this rearrangement was generated via the recently described ‘looping’ integration mechanism^16,42^. The α and β breakpoints fall within exons of the *NISCH* gene while the γ breakpoint falls within exon 27 of *PBRM1* (Fig. 6c), a gene previously shown to be a cancer driver in renal carcinoma^43^ and intrahepatic cholangiocarcinomas^44^.

**Figure 6.**
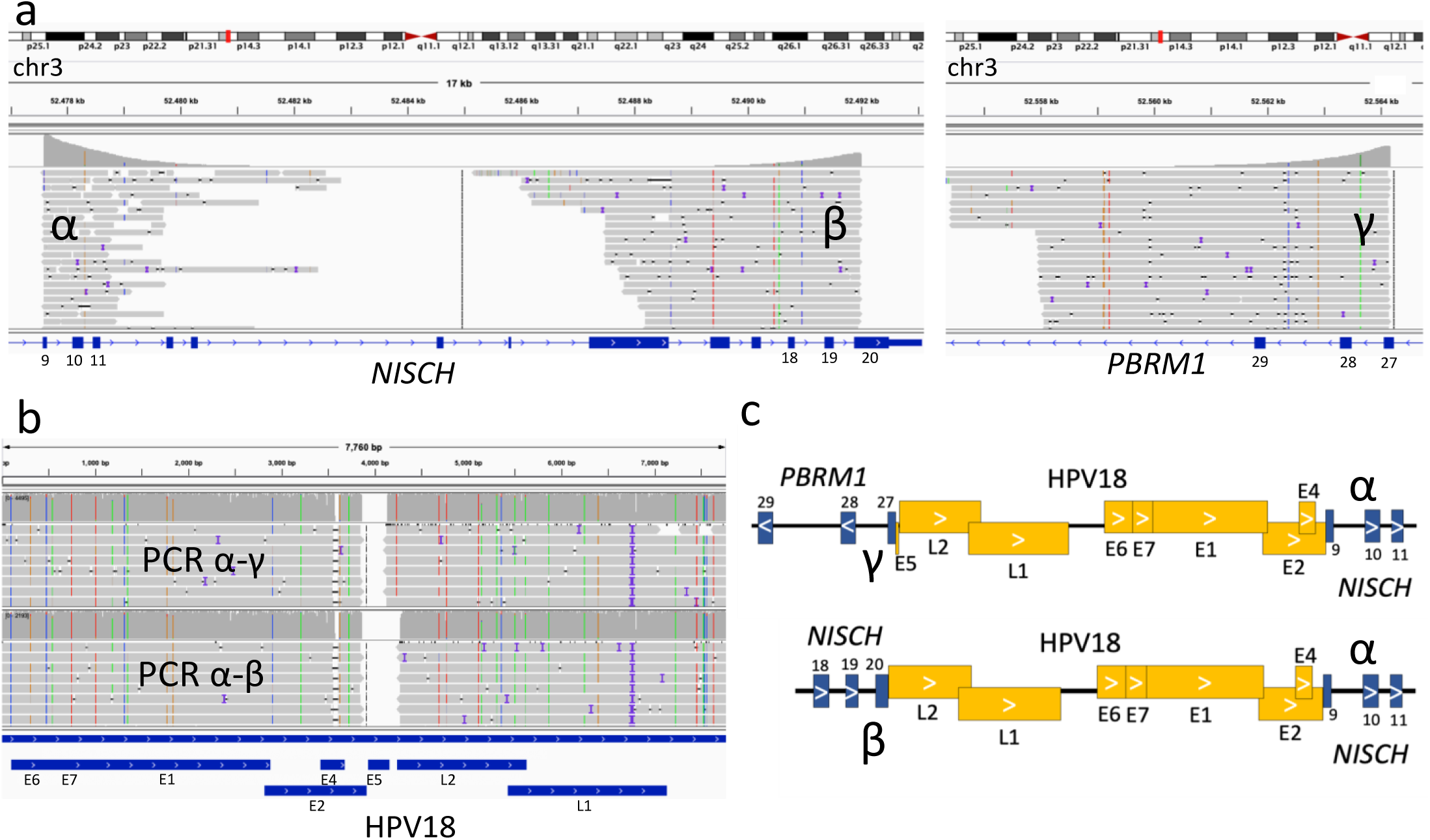
HPV ‘looping’ integration in an expanded clone **(a)** PCIP-seq reads mapping to a ∼87 kb region on chr3 revealed three HPV-host breakpoints. The large number of reads suggests expansion of the clone carrying these integrations. **(b)** PCR was carried out with primer pairs matching regions α and β, as well as α and γ. Both primer pairs produced a ∼9kb PCR product. Nanopore sequencing of the PCR products show the HPV genome connects these breakpoints. **(c)** Schematic of the breakpoints with the integrated HPV genome. This conformation indicates that this dramatic structural rearrangement in the host genome was generated via ‘looping’ integration of the HPV genome.

## Discussion

In the present report we describe how PCIP-seq can be utilized to identify insertion sites while also sequencing parts of, and in some cases the entire associated provirus, and confirm this methodology is effective with a number of different retroviruses as well as HPV. For insertion site identification, the method was capable of identifying more than ten thousand BLV insertion sites in a single sample, using ∼4μg of template DNA. Even in samples with a PVL of 0.66%, it was possible to identify hundreds of insertion sites with only 1μg of DNA as template. The improved performance of PCIP-seq in repetitive regions further highlights its utility, strictly from the standpoint of insertion site identification. In addition to its application in research, high throughput sequencing of retrovirus insertion sites has shown promise as a clinical tool to monitor ATL progression^20^. Illumina based techniques require access to a number of capital-intensive instruments. In contrast PCIP-seq libraries can be generated, sequenced and analyzed with the basics found in most molecular biology labs, moreover, preliminary results are available just minutes after sequencing begins^45^. As a consequence, the method may have use in a clinical context to track clonal evolutions in HTLV-1 infected individuals, especially as the majority of HTLV-1 infected individuals live in regions of the world with poor biomedical infrastructure^46^.

One of the common issues raised regarding Oxford Nanopore data is read accuracy. Early versions of the MinION had read identities of less than 60%^47^, however the development of new pores and base calling algorithms make read identities of ∼90% achievable^48^. Accuracy can be further improved by generating a consensus from multiple reads, making accuracies of ∼99.4%^48^ possible. Recently Greig et al^49^ compared the performance of Illumina and Oxford Nanopore technologies for SNP identification in two isolates of *Escherichia coli.* They found that after accounting for variants observed at 5-methylcytosine motif sequences only ∼7 discrepancies remained between the platforms. It should be noted that as PCIP-seq sequences PCR amplified DNA, errors generated by base modifications will be avoided. Despite these improvements in accuracy, Nanopore specific errors can be an issue at some positions (Supplementary Fig. 3a). Comparison with Illumina data is helpful in the identification of problematic regions and custom base calling models may be a way to improve accuracy in such regions^48^. Additionally, PCIP-seq libraries could equally be sequenced using long reads on the Pacific Biosciences platform or via 10X Genomics linked reads on Illumina if high single molecule accuracy is required^17^. In the current study we focused on SNPs observed in clonally expanded BLV proviruses. For viruses such as HIV-1, which have much lower proviral loads, more caution will be requited as the majority of proviral sequences will be generated from single provirus, making errors introduced by PCR more of an issue.

When analyzing SNPs from BLV the most striking result was the presence of the recurrent mutations at the first base of codon 303 in the viral protein Tax, a central player in the biology of both HTLV-1^46^ and BLV^50^. It has previously been reported that this mutation causes an E-to-K amino acid substitution which ablates the transactivator activity of the Tax protein^23^. Collectively, these observations suggest this mutation confers an advantage to clones carrying it, possibly contributing to immune evasion, while retaining Tax protein functions that contribute to clonal expansion. However, there is a cost to the virus as this mutation prevents infection of new cells due to the loss of Tax mediated transactivation of the proviral 5’LTR making it an evolutionary dead end. It will be interesting to see if PCIP-seq can provide a tool to identify other examples of variants that increase the fitness of the provirus in the context of an infected individual but hinder viral spread to new hosts. Additionally, the technique could be used to explore the demographic features of the proviral population within and between hosts, how these populations evolve over time and how they vary.

A second notable observation is the cluster of A-to-G transitions observed within a ∼70bp window in the 3’LTR. Similar patterns have been ascribed to ADAR1 hypermutation in a number of viruses^26^, including the close BLV relatives HTLV-2 and simian T-cell leukemia virus type 3 (STLV-3)^51^. Given the small number of hypermutated proviruses observed, it appears to be a minor source of variation in BLV, although it will be interesting to see it this holds for different retroviruses and at different time points during infection.

Only a small fraction of proviruses (∼2.4%) in the HIV-1 reservoir are intact, yet these are more than sufficient for the disease to rebound if antiretroviral therapy is interupted^5^. As strategies are developed to target these intact proviruses it will be essential to distinguish between intact and defective proviruses^5^. As can be seen from the two long term cART patients examined, PCIP-seq is capable of identifying proviral integration sites, sequencing part, and in some cases all of the associated provirus. While the number of patients examined in this work is limited, it is interesting to note that we were able to extract full length, intact sequences for two proviruses in patient 02006 (on cART for 15 years), indicating that both are present in clonally expanded cells. These proviruses are integrated within highly repetitive/heterochromatic regions and as a result they are likely to be resistant to reactivation. This result is somewhat surprising and additional samples will need to be sequenced to determine how common HIV-1 integration in such regions is. Others have recently speculated that proviruses integrated into parts of the genome that provide an unfavorable environment for viral expression are protected against recognition by the host immune system, favoring their survival in patients on long term cART^6^.

In the current study we focused our analysis on retroviruses and ERVs. However, as this methodology is potentially applicable to a number of different targets we extended its use to HPV as a proof of concept. It is estimated that HPV is responsible for >95% of cervical carcinoma and ∼70% of oropharyngeal carcinoma^52^. While infection with a high-risk HPV strain (HPV16 & HPV18) is generally necessary for the development of cervical cancer, it is not sufficient and the majority of infections resolve without adverse consequences^41^. The use of next-generation sequencing has highlighted the central role HPV integration plays in driving the development of cervical cancer^16^. Our results show that PCIP-seq can be applied to identify HPV integration sites in early precancerous samples. This opens up the possibility of generating a more detailed map of HPV integrations as well as potentially providing a biomarker to identify HPV integrations on the road to cervical cancer.

Looking beyond viruses tested in the current study, hepatitis B virus (HBV) is an obvious candidate for PCIP-seq. Like HPV it has a circular DNA genome that integrates into the host genome with variable breakpoints in the viral genome. HBV integrations contribute to genomic instability and play a key role in driving hepatocarcinogenesis^53^. Other potential applications include determining the insertion sites and integrity of retroviral vectors^54^ and detecting transgenes in genetically modified organisms. We envision that in addition to the potential applications outlined above many other novel targets/questions could be addressed using this method.

## Methods

### Samples

Both the BLV infected sheep^7^ and HTLV-1 samples^7,20^ have been previously described. Briefly, the sheep were infected with the molecular clone pBLV344^21^, following the experimental procedures approved by the University of Saskatchewan Animal Care Committee based on the Canadian Council on Animal Care Guidelines (Protocol #19940212). The HTLV-1 samples^7,20^ were obtained with informed consent following the institutional review board-approved protocol at the Necker Hospital, University of Paris, France, in accordance with the Declaration of Helsinki. The BLV bovine samples were natural infections, obtained from commercially kept adult dairy cows in Alberta, Canada. Sampling was approved by VSACC (Veterinary Sciences Animal care Committee) of the University of Calgary: protocol number: AC15-0159. The bovine 571 used for ERV identification was collected as part of this cohort. The two sheep samples used for Jaagsiekte sheep retrovirus (enJSRV) identification were the BLV infected ovine samples (220 & 221 (032014)), with a PVL of 3.8 and 16% respectively. PBMCs were isolated using standard Ficoll-Hypaque separation. The DNA for the bovine Mannequin was extracted from sperm, while the DNA for bovine 10201e6 was extracted from whole blood using standard procedures. The HIV-1 U1 cell line DNA sequenced without dilution was provided by Dr. Carine Van Lint, IBMM, Gosselies, Belgium. The HIV-1 U1 cell line dilutions in Jurkat were generated at Ghent University Hospital. HIV-1 positive primary PBMCs were collected at the Ghent University Hospital from two HIV-1-positive individuals (patients 02006 and 06042, Supplementary Table 4) on cART for 15 and 8 years respectively. The study was approved by the Ethics Committee of Ghent University Hospital (Reference number: 2016/0457). HPV material was prepared from PAP smears obtained from HPV-infected patients at the CHU Liège University hospital. Both patients were PCR positive for HPV18, HPV18_PY was classified as having Atypical Squamous Cell of Undetermined Significance (ASC-US), while HPV18_PX was classified as having Atypical Glandular Cells (AGC). Patients provided written informed consent and the study was approved by the Comité d’Ethique Hospitalo-Facultaire Universitaire de Liège (Reference number: 2019/139). No statistical test was used to determine adequate sample size and the study did not use blinding.

### CD4 enrichment of HIV-1 patient PBMCs

CD4^+^ T cells were enriched from PBMCs by negative MACS selection using the EasySep™ Human CD4^+^ T Cell Isolation Kit (STEMCELL Technologies SARL, Grenoble, France), according to manufacturer’s instructions.

### PCIP-seq

Total genomic DNA isolation was carried out using the Qiagen AllPrep DNA/RNA/miRNA kit (BLV, HTLV-1 and HPV infected individuals) or the Qiagen DNeasy Blood & Tissue Kit (HIV-1 patients) according to manufacturer’s protocol. High molecular weight DNA was sheared to ∼8kb using Covaris g-tubesTM (Woburn, MA) or a Megaruptor (Diagenode), followed by end-repair using the NEBNext EndRepair Module (New England Biolabs). Intramolecular circularization was achieved by overnight incubation at 16°C with T4 DNA Ligase. Remaining linear DNA was removed with Plasmid-Safe-ATP-Dependent DNAse (Epicentre, Madison WI). Guide RNAs were designed using chopchop (http://chopchop.cbu.uib.no/index.php). The EnGen™ sgRNA Template Oligo Designer (http://nebiocalculator.neb.com/#!/sgrna) provided the final oligo sequence. Oligos were synthesized by Integrated DNA Technologies (IDT). Oligos were pooled and guide RNAs synthesized with the EnGen sgRNA Synthesis kit, S. pyogenes (New England Biolabs). Selective linearization reactions were performed with the Cas-9 nuclease, S. pyogenes (New England Biolabs). (See Supplementary Note 1 for the rationale behind using of CRISPR-cas9 to cleave the circular DNA). PCR primers flanking the cut sites were designed using primer3 (http://bioinfo.ut.ee/primer3/). Primers were tailed to facilitate the addition of Oxford Nanopore indexes in a subsequent PCR reaction. The linearized fragments were PCR amplified with LongAmp Taq DNA Polymerase (New England Biolabs) and purified using 1x AmpureXP beads, (Beckman Coulter). A second PCR added the appropriate Oxford Nanopore index. PCR products were visualized on a 1% agarose gel, purified using 1x AmpureXP beads and quantified on a Nanodrop spectrophotometer. Indexed PCR products were multiplexed and Oxford Nanopore libraries prepared with either the Ligation Sequencing Kit 1D (SQK-LSK108) or 1D^2 Sequencing Kit (SQK-LSK308) (only the 1D were used) The resulting libraries were sequenced on Oxford Nanopore MinION R9.4 or R9.5 flow cells respectively. The endogenous retrovirus libraries were base called using albacore 2.3.1, all other PCIP-seq libraries were base called with Guppy 3.1.5 (https://nanoporetech.com) using the “high accuracy” base calling model. For the endogenous retrovirus libraries, demultiplexing was carried out via porechop (https://github.com/rrwick/Porechop) using the default setting. The HIV, HTLV-1, BLV and HPV PCIP-seq libraries were subjected to a more stringent demultiplexing with the guppy_barcoder (https://nanoporetech.com) tool using the --require_barcodes_both_ends option. The output was also passed through porechop, again barcodes were required on both ends, adapter sequence was trimmed and reads with middle adapters were discarded.

### Identification of proviral integrations sites in PCIP-seq

Reads were mapped with Minimap2^55^ to the host genome with the proviral genome as a separate chromosome. In-house R-scripts were used to identify integration sites (IS). Briefly, chimeric reads that partially mapped to at least one extremity of the proviral genome were used to extract virus-host junctions and shear sites. Junctions within a 200bp window were clustered together to form an “IS cluster”, compensating for sequencing/mapping errors. The IS retained corresponded to the position supported by the highest number of virus-host junctions in each IS cluster. Clone abundance was estimated based on the number of reads supporting each IS cluster. Reads sharing the same integration site and same shear site were considered PCR duplicates. Custom software, code description and detailed outline of the workflow are available on Github: https://github.com/GIGA-AnimalGenomics-BLV/PCIP.

### Measure of proviral load (PVL) and identification of proviral integration sites (Illumina)

PVLs and integration sites of HTLV-1- and BLV-positive individuals were determined as previously described in Rosewick et al 2017^7^ and Artesi et al 2017^20^. PVL represents the percentage of infected cells, considering a single proviral integration per cell. Total HIV-1 DNA content of CD4 T-cell DNA isolates was measured by digital droplet PCR (ddPCR, QX200 platform, Bio-Rad, Temse, Belgium), as described by Rutsaert et al.^56^ The DNA was subjected to a restriction digest with EcoRI (Promega, Leiden, The Netherlands) for one hour, and diluted 1:2 in nuclease free water. HIV-1 DNA was measured in triplicate using 4 µL of the diluted DNA as input into a 20µL reaction, while the RPP30 reference gene was measured in duplicate using 1 µL as input. Thermocycling conditions were as follows: 95°C for 10 min, followed by 40 cycles of 95°C for 30 s and 56°C for 60 s, followed by 98°C for 10 min. Data was analyzed with the ddpcRquant analysis software^57^.

### Variant Calling

After PCR duplicate removal, proviruses with an IS supported by more than 10 reads were retained for further processing. SNPs were identified using LoFreq^22^ with default parameters, only SNPs with an allele frequency of >0.6 in the provirus associated with the insertion site were considered. We also called variants on proviruses supported by more than 10 reads without PCR duplicate removal (this greatly increased the number of proviruses examined). This data was used to explore the number of proviruses carrying the Tax 303 variant. Deletions were called on proviruses supported by more than 10 reads without PCR duplicate removal using an in house R-scripts. Briefly, samtools pileup^58^ was used to calculate/compute coverage and deletions at base resolution. We used the changepoint detection algorithm PELT^59^ to identify genomic windows showing an abrupt change in coverage. Windows that showed at least a 4-fold increase in the frequency of deletions (absence of a nucleotide for that position within a read) were flagged as deletions and visually confirmed in IGV^60^.

### HIV-1 proviral sequences

Sequences of the two major proviruses integrated in chr2 and chrX of the U1 cell line were generated by initially mapping the reads from both platforms to the HIV-1 provirus, isolate NY5 (GenBank: M38431.1), where the 5’LTR sequence is appended to the end of the sequence to produce a full-length HIV-1 proviral genome reference. The sequence was then manually curated to produce the sequence for each provirus. To check for recombination, reads of selected clones were mapped to the sequence from the chrX provirus and the patterns of SNPs examined to determine if the variants matched the chrX or chr2 proviruses.

The consensus HIV-1 sequences for patient 02006 as well as the individual provirus sequences from this patient was generated using medaka (https://github.com/nanoporetech/medaka), followed by manual correction guided by Illumina reads generated from the same PCIP-seq library. The Illumina libraries were prepared as described in Durkin et al 2016^29^, briefly, this involved fragmenting the PCIP-seq library, adding Nextera indexes and sequencing on a Illumina miSeq. Hypermutation of the provirus was initially identified by manually inspecting the reads in IGV, the consensus sequence of the provirus was checked for hypermutation with Hypermut (https://www.hiv.lanl.gov/content/sequence/HYPERMUT/hypermut.html). We determined if the proviral sequences were intact using the Gene Cutter tool (https://www.hiv.lanl.gov/content/sequence/GENE_CUTTER/cutter.html). Proviruses that did not contain a frameshift or stop codons not observed in the consensus sequence generated for patient 02006 were classified as intact. Deletions in the HIV-1 proviruses were identified by manual inspection of the integration site and proviral reads in IGV.

### Endogenous retroviruses

The sequence of bovine *APOB* ERV was generated by PCR amplifying the full length ERV with LongAmp Taq DNA Polymerase (New England Biolabs) from a Holstein suffering from cholesterol deficiency. The resultant PCR product was sequenced on the Illumina platform as described below. It was also sequenced with an Oxford Nanopore MinION R7 flow cell as previously described^29^. Full length sequence of the element was generated via manual curation. Guide RNAs and primer pairs were designed using this ERV reference. For the Ovine ERV we used the previously published enJSRV-7 sequence^40^ as a reference to design PCIP-seq guide RNAs and PCR primers.

As the ovine and bovine genome contains sequences matching the ERV, mapping ERV PCIP-seq reads back to the reference genome creates a large pileup of reads in these regions. To avoid this, prior to mapping to the reference we first used BLAST^61^ to identify the regions in the reference genome containing sequences matching the ERV, we then used BEDtools^62^ to mask those regions. The appropriate ERV reference was then added as an additional chromosome in the reference.

### PCR validation and Illumina sequencing

Clone specific PCR products were generated by placing primers in the flanking DNA as well as inside the provirus. LongAmp Taq DNA Polymerase (New England Biolabs) was used for amplification following the manufacturer’s guidelines. Resultant PCR products were sheared to ∼400bp using the Bioruptor Pico (Diagenode) and Nextera XT indexes added as previously described^29^. Illumina PCIP-seq libraries were generated in the same manner. Sequencing was carried out on either an Illumina MiSeq or NextSeq 500. Clone specific PCR products sequenced on Nanopore were indexed by PCR, multiplexed and libraries prepared using the Ligation Sequencing Kit 1D (SQK-LSK108) and sequenced on a MinION R9.4 flow cell.

### BLV references

The sequence of the pBLV344 provirus was generated via a combination of Sanger and Illumina based sequencing with manual curation of the sequence to produce a full length proviral sequence. The consensus BLV sequences for the bovine samples 1439 & 1053 were generated by first mapping the PCIP-seq Nanopore reads to the pBLV344 provirus. We then used Nanopolish^63^ to create an improved consensus. PCIP-seq libraries sequenced on the Illumina and Nanopore platform were mapped to this improved consensus visualized in IGV and manually corrected.

### Genome references used

Sheep=OAR3.1 Cattle=UMD3.1 Human=hg38 For HTLV-1 integration sites hg19 was used HPV18=GenBank: AY262282.1

## Acknowledgements

This work was supported by les Amis de l’Institut Bordet, the Fonds de la Recherche Scientifique (FRS), Télévie, the International Brachet Stiftung (IBS), the Région wallonne WALInnov project CAUSEL (convention n° 1710030), WALGEMED, and a Télévie Grant to V.H. M.A. holds a Post-doctoral Researcher fellowship of the FRS. K.D. is a Scientific Research Worker of Télévie. L.V. is supported by the Research Foundation Flanders (FWO 1.8.020.09.N.00). B.C. received a strategic basic research fund of the Research Foundation – Flanders (grant number FWO 1S28920N). Dr. Carine Van Lint kindly provided DNA from the U1 HIV-1 cell line. Computational resources were provided by GIGA and the Consortium des Équipements de Calcul Intensif (CÉCI). We thank Wouter Coppieters, Latifa Karim, Manon Deckers and the GIGA Genomics Platform for sequencing services. We thank Dr. Lionel Habran, Sonia Pisvin, Hélène Piron, Renée Gathy (Laboratory of molecular biology and immunohistology of the University Hospital of Liége) and Dr. Stéphanie Gofflot (BUL, Biobank of the University of Liège) for their assistance with HPV sample selection.

## Author contributions

K.D. conceived and designed the study, K.D and M.A. optimized the method, generated and analyzed data. V.H. developed the bioinformatics pipeline and analyzed data. F.A. contributed to data generation. A.M. and O.H. provided HTLV-1 patient materials, P.D., B.E. and V. B. took care of HPV diagnostic samples, L.V., B.C. and L.L. provided, processed and analyzed HIV-1 patient and U1/Jurkat dilutions samples, P.G., N.A. and F.V. collected and provided animal samples. K.D. wrote the first draft, M.A., V.H., A.V., P.G., M.G, C.C., L.V. contributed to the final manuscript. A.V. and M.G. supervised the study.

