## Supplementary figures and tables for "Pooled CRISPR Inverse PCR sequencing (PCIP-seq): simultaneous sequencing of retroviral insertion points and the integrated provirus with long reads"

**Supplementary Note**

**Rationale behind the use of CRISPR-cas9 to cleave circular DNA**

It is established practice to linearize plasmids (generally via cutting with a restriction enzyme) prior to their use as template in PCR. It is believed that this avoids supercoiling and thereby increases PCR efficiency[1]. Following the same logic, we speculated that linearizing our circularized DNA could also increase PCR efficiency. The gel below shows an experiment carried out using 8ug of DNA from a BLV infected sheep with a proviral load of 82.6%. The DNA was circularized and linear DNA was eliminated (to prevent PCR amplification/recombination involving the remaining linear fragments) using plasmid safe DNase (see methods for a complete description). One quarter of the resultant DNA was subject to CRISPR-cas9 cleavage using the Pool A guides, the second quarter was cleaved using the Pool B guides, the remaining half was kept aside. The linearized DNA was cleaned and used as template in 2x 50ul PCR reactions using the appropriate primer pairs for Pool A (PA) or Pool B (PB). For the uncut DNA half was used as template for 2x 50ul PCR reactions using the PA primers and the other half was used for 2x 50ul PCR reactions using the PB primers. Following 25 PCR cycles, 10ul of each reaction were loaded on a 1% agarose gel. As can be seen in the gel below, the band intensity for the CRISPR-cas9 cut samples is higher. It should be noted that in lane 3 the PCR smear is shifted down, we generally discard these types of products as the fraction of host-virus fragments is low. (A=unshared genomic DNA, B=genomic DNA sheared to 8kb)

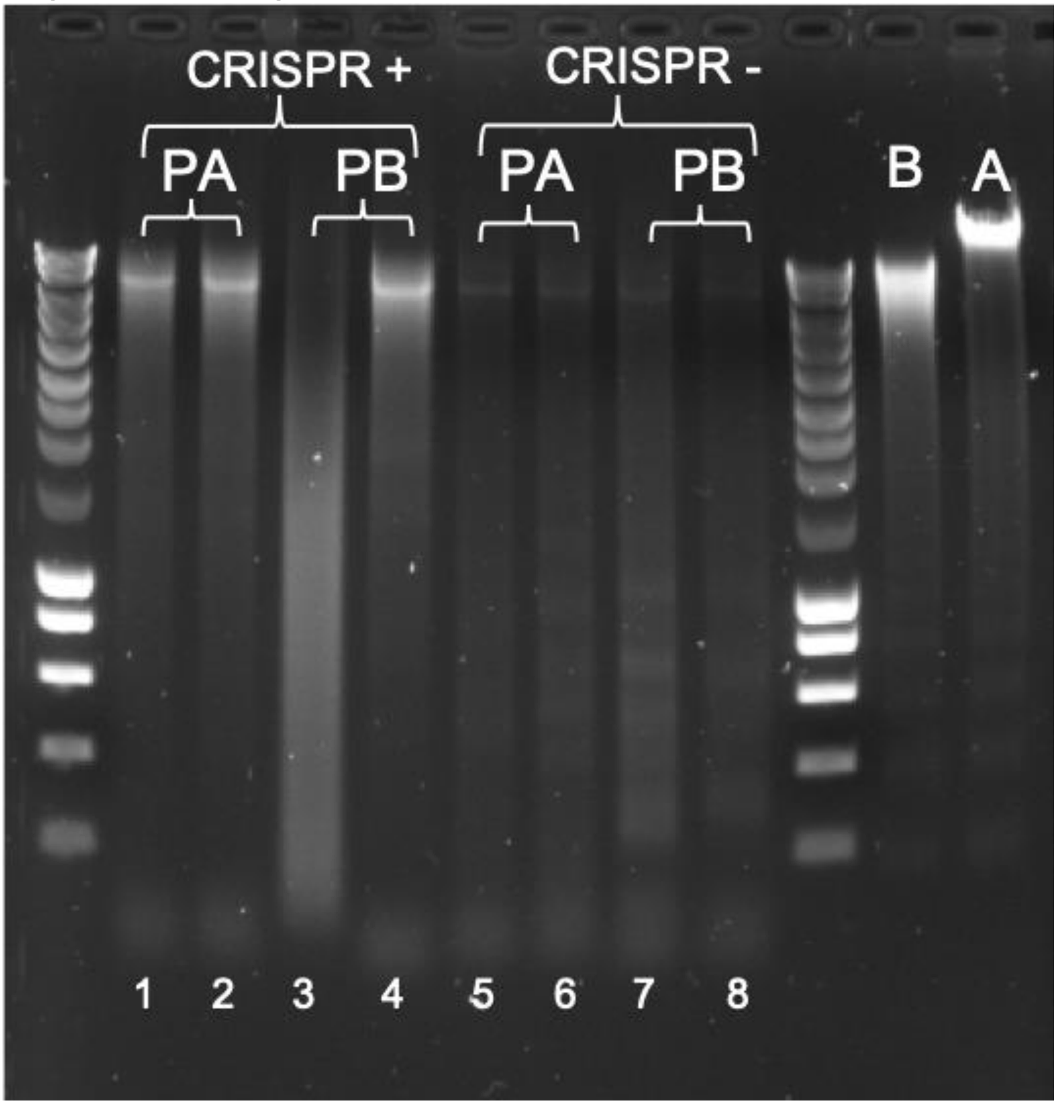


Following clean up and elution in ~40 ul of H2O we took an equal volume (3ul) of each library and indexed them via PCR, in a 50 ul reaction volume and using 8 cycles. Again, following clean up, an equal volume of library was pooled and a nanopore library (LSK-109) was prepared and sequenced on a r9.4 flow cell. Base calling and demultiplexing was carried out as described in the methods. The results are outlined in the table below. In addition the coverage of the resultant reads is shown.

| Lib | Treatment | DNA concentration PCR 1 (ng/ul) | DNA concentration PCR 2 (ng/ul) | Raw reads | Chimeric reads % | Pure Host / Pure Viral reads (%) | Mean Length | N50 | Median Length | Insertion sites PCIP | Largest clone PCIP (%) | Insertion sites Illumina | Largest clone Illumina (%) |
|---|---|---|---|---|---|---|---|---|---|---|---|---|---|
| 1 | PA-Cut BC31 | 22.52 | 69.48 | 113,485 | 55.6 | 0.25 / 44.2 | 2880.6 | 3855.0 | 2217.0 | 2122 | 25.8 | 1700 | 30.849 |
| 2 | PA-Cut BC32 | 26.18 | 72.06 | 137,109 | 54.1 | 0.47 / 45.4 | 2770.6 | 3710.0 | 2141.0 | 2216 | 24.7 | “ | “ |
| 3 | PB-Cut BC33 | 71.85 | 63.7 | 6,844 | 1.01 | 98.5 / 0.51 | 263.8 | 277.0 | 195.5 | 2 | 50 | “ | “ |
| 4 | PB-Cut BC34 | 34.17 | 86.65 | 126,655 | 49.4 | 0.19 / 50.4 | 2616.2 | 3395.0 | 2010.0 | 2281 | 24.5 | “ | “ |
| 5 | PA-UnCut BC35 | 13.4 | 33.32 | 42,795 | 22.5 | 0.19 / 77.3 | 1759.8 | 2670.0 | 1227.0 | 660 | 30.9 | “ | “ |
| 6 | PA-UnCut BC36 | 17.26 | 42.53 | 66,602 | 19.7 | 0.19 /80.2 | 1549.1 | 2381.0 | 1056.0 | 713 | 30.4 | “ | “ |
| 7 | PB-UnCut BC37 | 22.27 | 48.24 | 114,967 | 10.4 | 0.16 / 89.4 | 917.9 | 1579.0 | 497.0 | 690 | 29.5 | “ | “ |
| 8 | PB-UnCut BC38 | 14.71 | 35.92 | 64,789 | 18.1 | 0.19 / 81.7 | 1461.4 | 2111.0 | 992.0 | 736 | 30.4 | “ | “ |

The table shows that libraries prepared with the CRISPR cut generally produced more raw reads and a much larger fraction of them is composed of the desired chimeric reads containing proviral and host DNA. The CRISPR cut libraries also identified a large number of integration sites. The comparison with an Illumina based library prepared from the same timepoint, using ~4ug of template, shows that PCIP can identify more integration sites. This experiment also shows that only libraries with a size distribution that mirrors that observed in the sheared DNA should be sequenced, libraries with a preponderance of shorter fragments mainly represent nonspecific amplification.


1. Chen, J., Kadlubar, F. F. & Chen, J. Z. DNA supercoiling suppresses real-time PCR: a new approach to the quantification of mitochondrial DNA damage and repair. *Nucleic Acids Res* **35,** 1377–1388 (2007).

Coverage of the pure viral reads as well as the chimeric reads on the BLV proviral genome
(BC refers to the barcode used for each library)

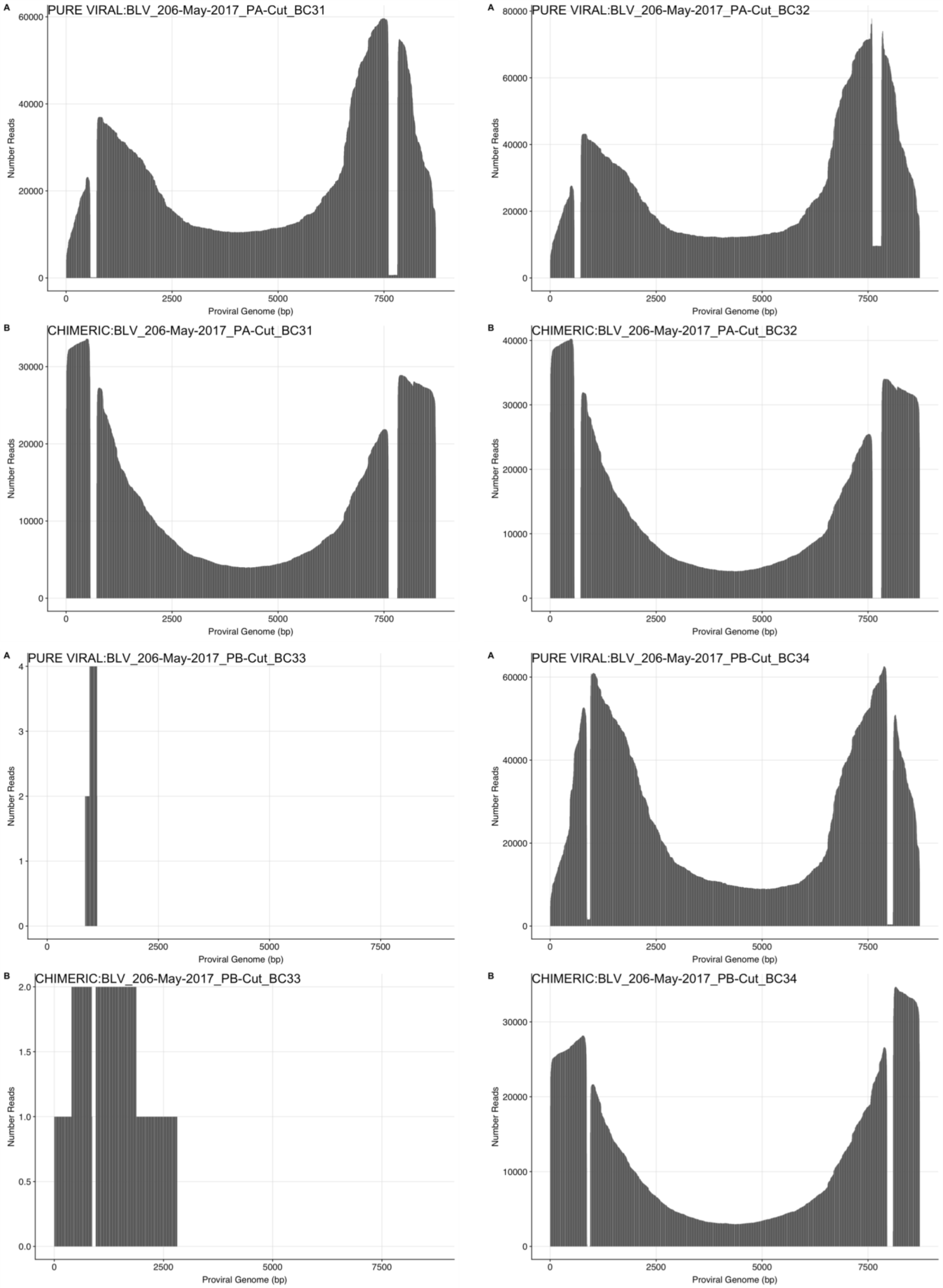

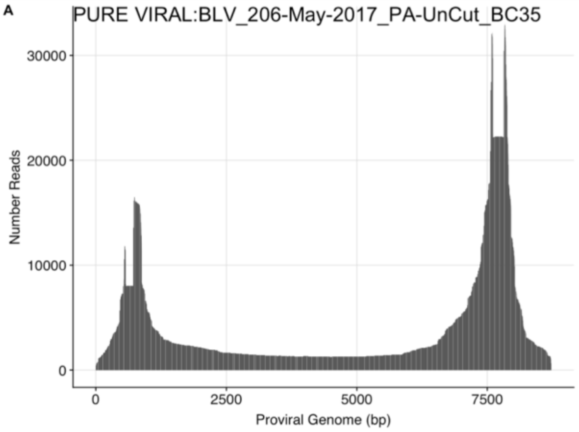
A
PURE VIRAL:BLV_206-May-2017_PA-UnCut_BC35
Number Reads
0
10000
20000
30000
Proviral Genome (bp)
0
2500
5000
7500

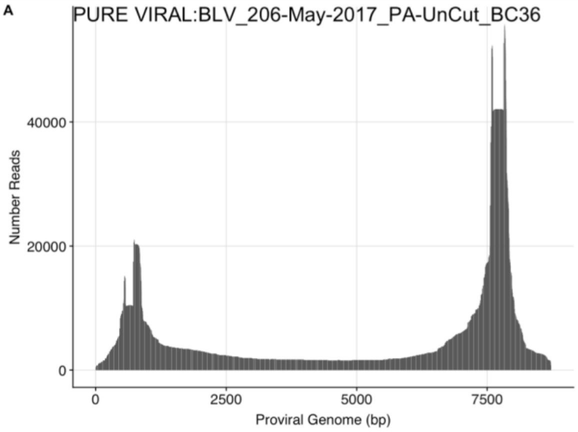
A
PURE VIRAL:BLV_206-May-2017_PA-UnCut_BC36
Number Reads
0
20000
40000
Proviral Genome (bp)
0
2500
5000
7500

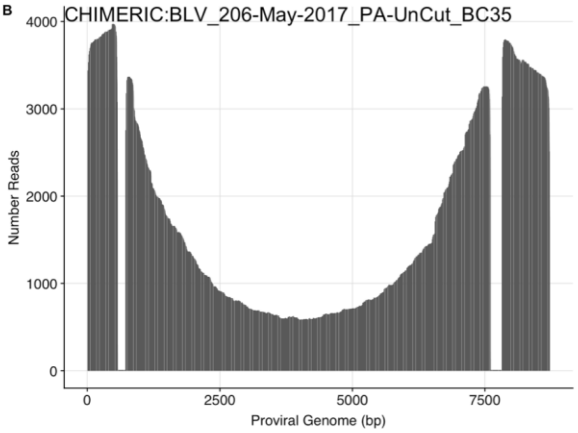
B
CHIMERIC:BLV_206-May-2017_PA-UnCut_BC35
Number Reads
0
1000
2000
3000
4000
Proviral Genome (bp)
0
2500
5000
7500

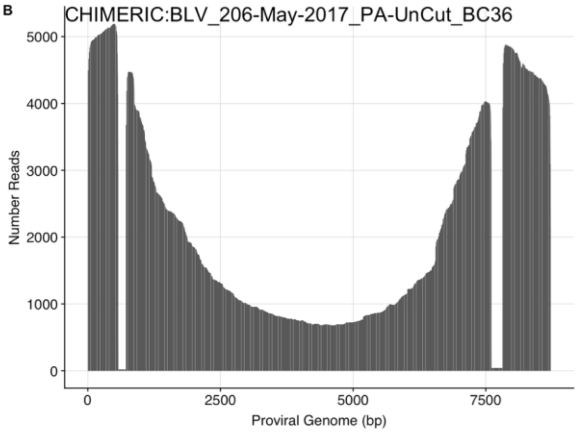
B
CHIMERIC:BLV_206-May-2017_PA-UnCut_BC36
Number Reads
0
1000
2000
3000
4000
5000
Proviral Genome (bp)
0
2500
5000
7500

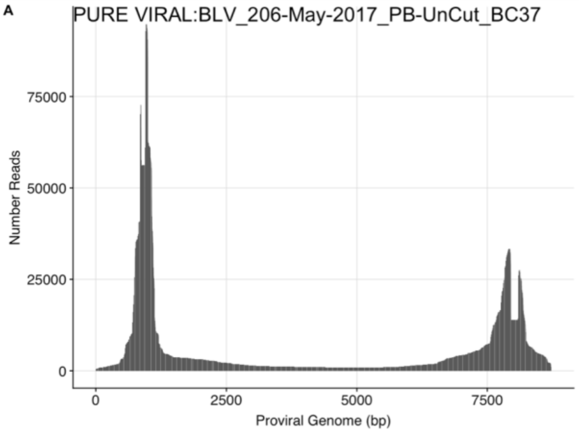
A
PURE VIRAL:BLV_206-May-2017_PB-UnCut_BC37
Number Reads
0
25000
50000
75000
Proviral Genome (bp)
0
2500
5000
7500

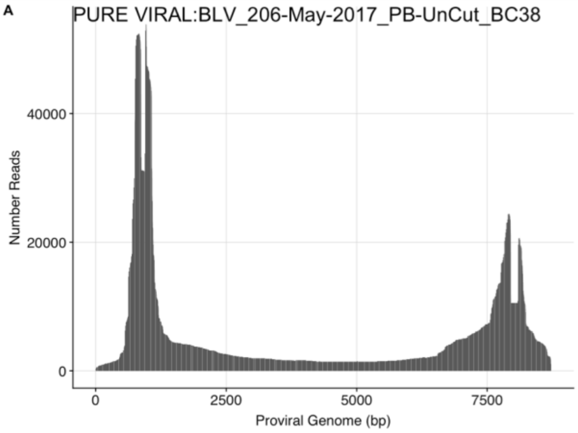
A
PURE VIRAL:BLV_206-May-2017_PB-UnCut_BC38
Number Reads
0
20000
40000
Proviral Genome (bp)
0
2500
5000
7500

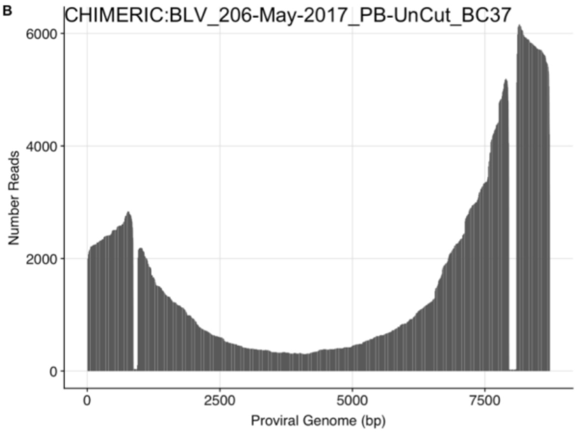
B
CHIMERIC:BLV_206-May-2017_PB-UnCut_BC37
Number Reads
0
2000
4000
6000
Proviral Genome (bp)
0
2500
5000
7500

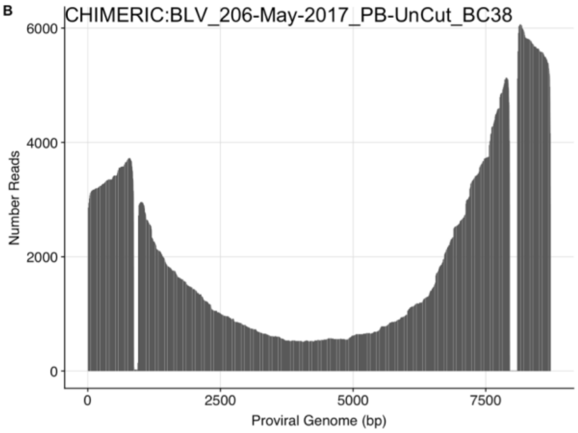
B
CHIMERIC:BLV_206-May-2017_PB-UnCut_BC38
Number Reads
0
2000
4000
6000
Proviral Genome (bp)
0
2500
5000
7500
