## Supplementary note for "Pooled CRISPR Inverse PCR sequencing (PCIP-seq): simultaneous sequencing of retroviral insertion points and the integrated provirus with long reads"

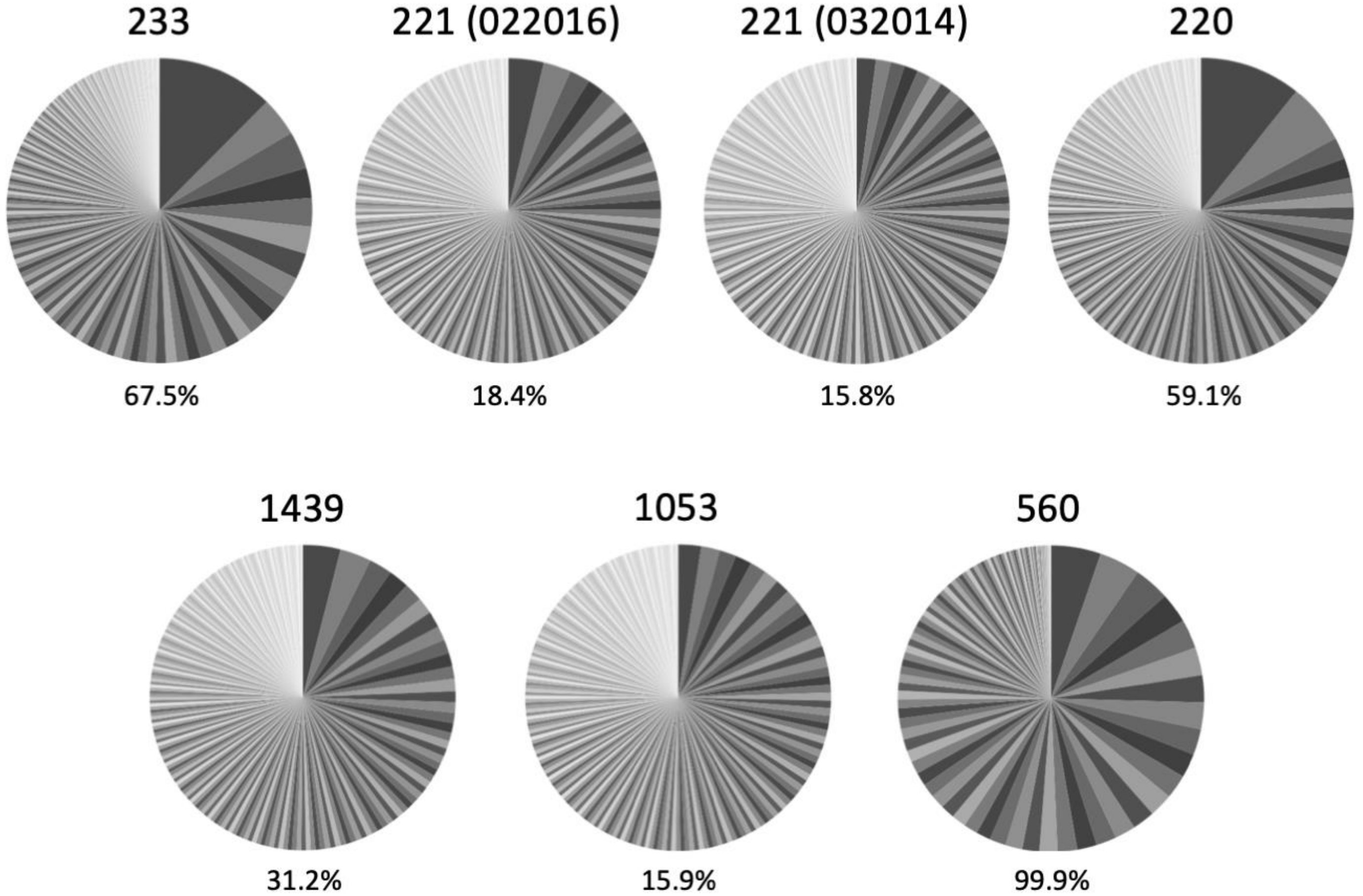

**Supplementary Figure 1** Pie charts showing the relative abundance of the 200 largest clones in the four sheep (top) and three cattle (bottom) infected with BLV, each slice of the pie represents a single insertion site, the % below indicated what fraction of the overall reads these 200 clones represent.

### Ovine 221 (022016) & 221 (032014)
### BLV SNPs validated via clone specific PCR

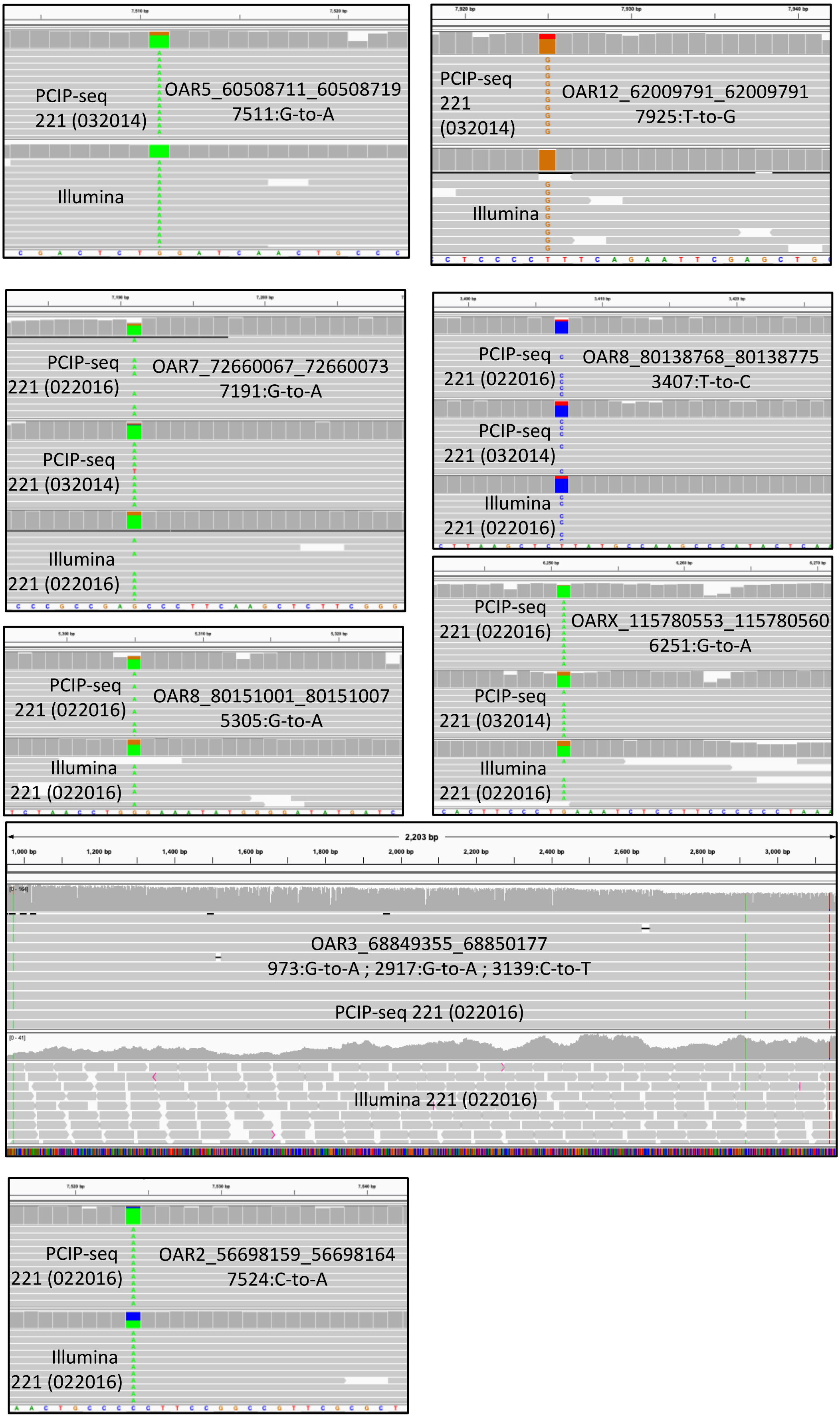

**Supplementary Figure 2** SNPs identified by PCIP-seq in BLV validated by clone specific PCR.

### Bovine 1439 BLV SNPs validated via clone specific PCR

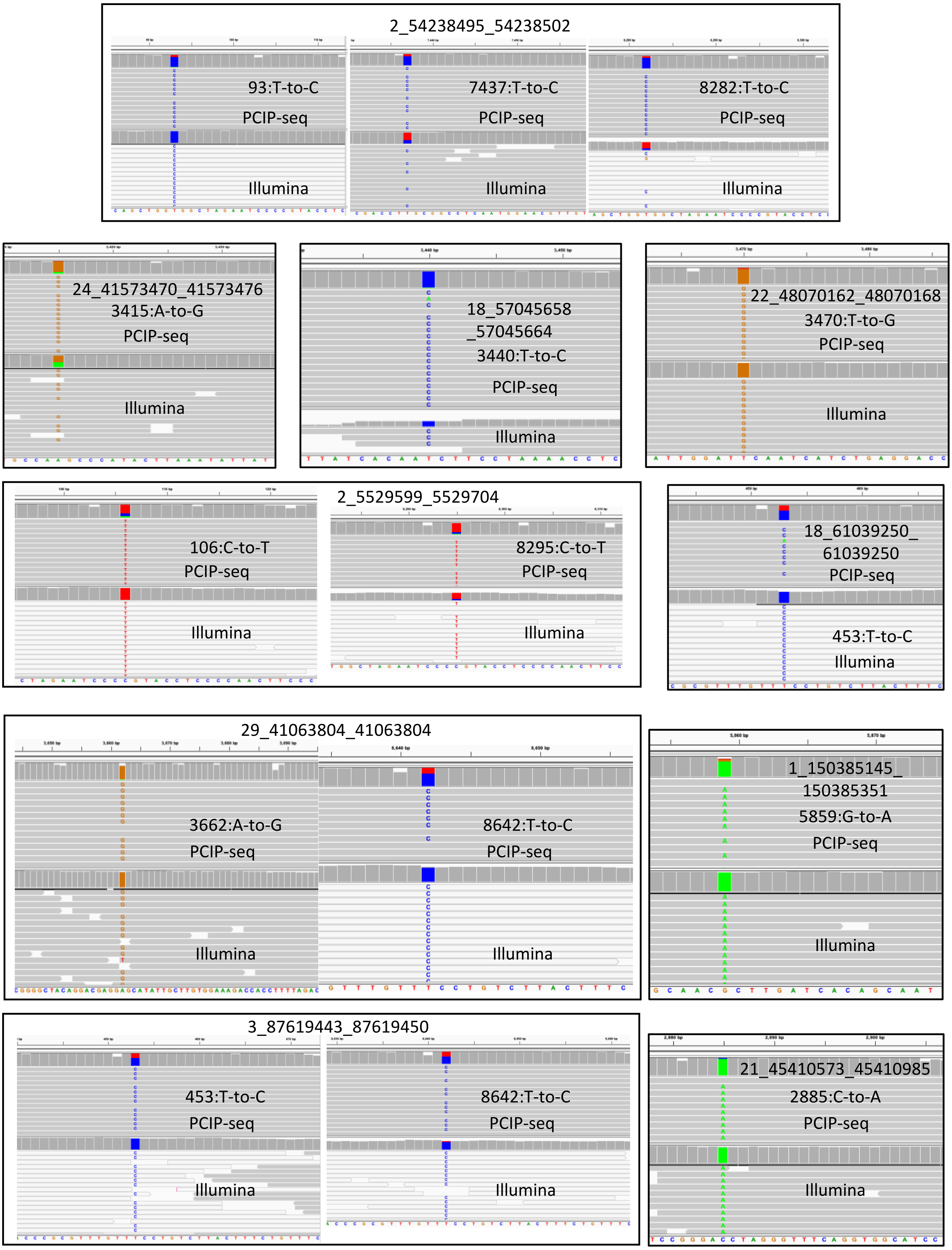

**Supplementary Figure 2 continued**

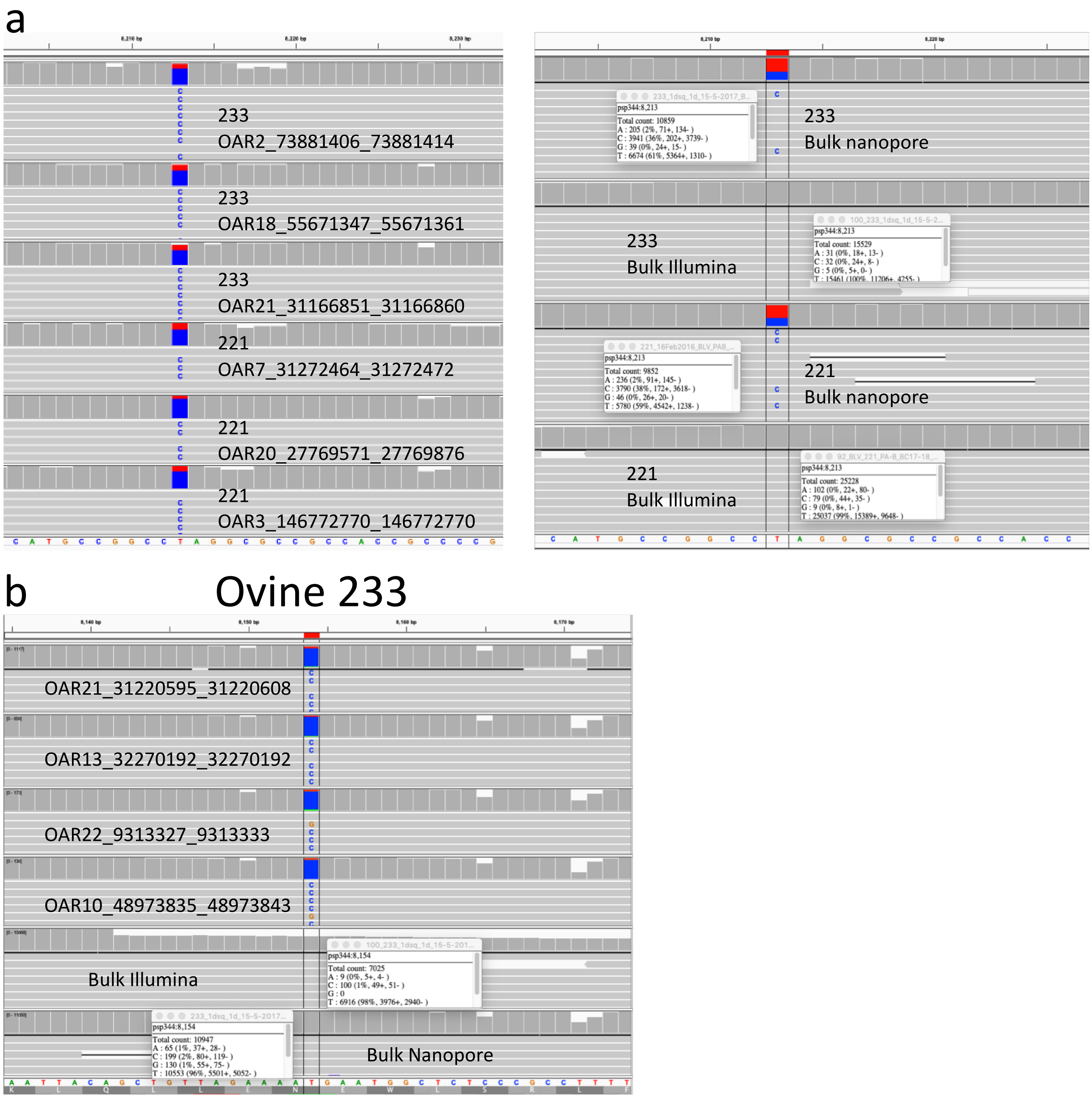

**Supplementary Figure 3** Distinguishing between real SNPs and technical artifacts **(a)** We observed a number of BLV proviruses in all the samples that had an apparent SNP at position 8213. Shown are three examples from sheep 233 and 211. When we looked at this position in reads mapped to the provirus without first sorting based on insertion site (referred to as bulk) we saw a C called 36 and 38% of the time respectively in the Nanopore data. In the bulk Illumina data, generated from the same sample, we saw the C is called 0% of the time indicating a technical artifact. As a consequence, SNPs from this position were excluded. **(b)** In animal 233 we found 16 proviruses (provirus inclusion was based on the less stringent criteria of >10 reads covering the position, not filtered for PCR duplicates) carrying a T-to-C transition within the Tax ORF at position 8154, this variant does not change the amino acid. Shown are screen captures for 4 of the proviruses carrying the SNP, Illumina and Nanopore bulk sequencing from the same sample show C is called at a 2% frequency in Nanopore, while with Illumina C is called at a 1% frequency. This indicates that the SNPs observed in these proviruses are not a technical artifact.

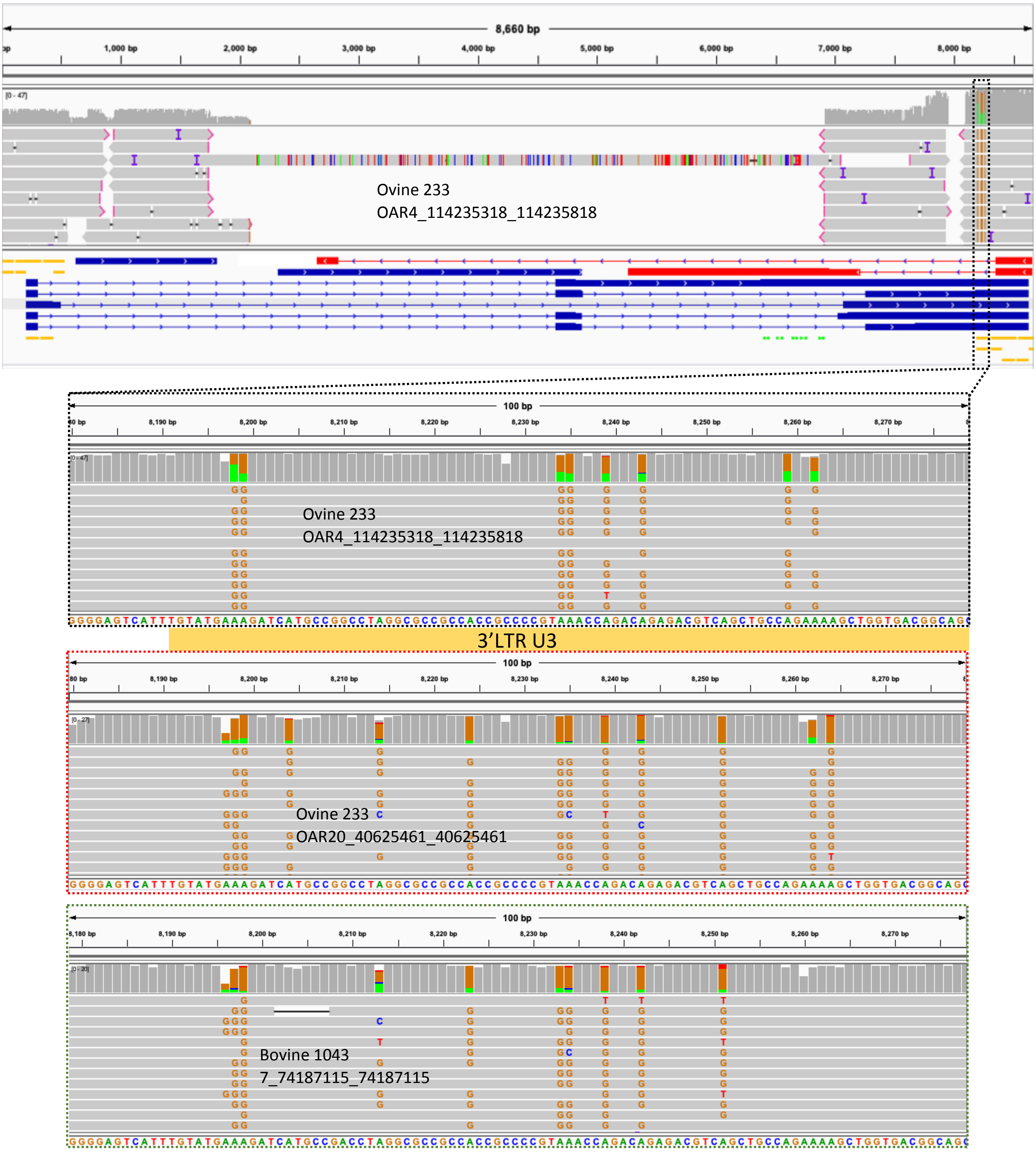

**Supplementary Figure 4** Hypermutation of a ~70bp region in U3 of the 3’LTR of BLV proviruses.

### Ovine 221 (022016) & 221 (032014)
### BLV SVs validated by clone specific PCR

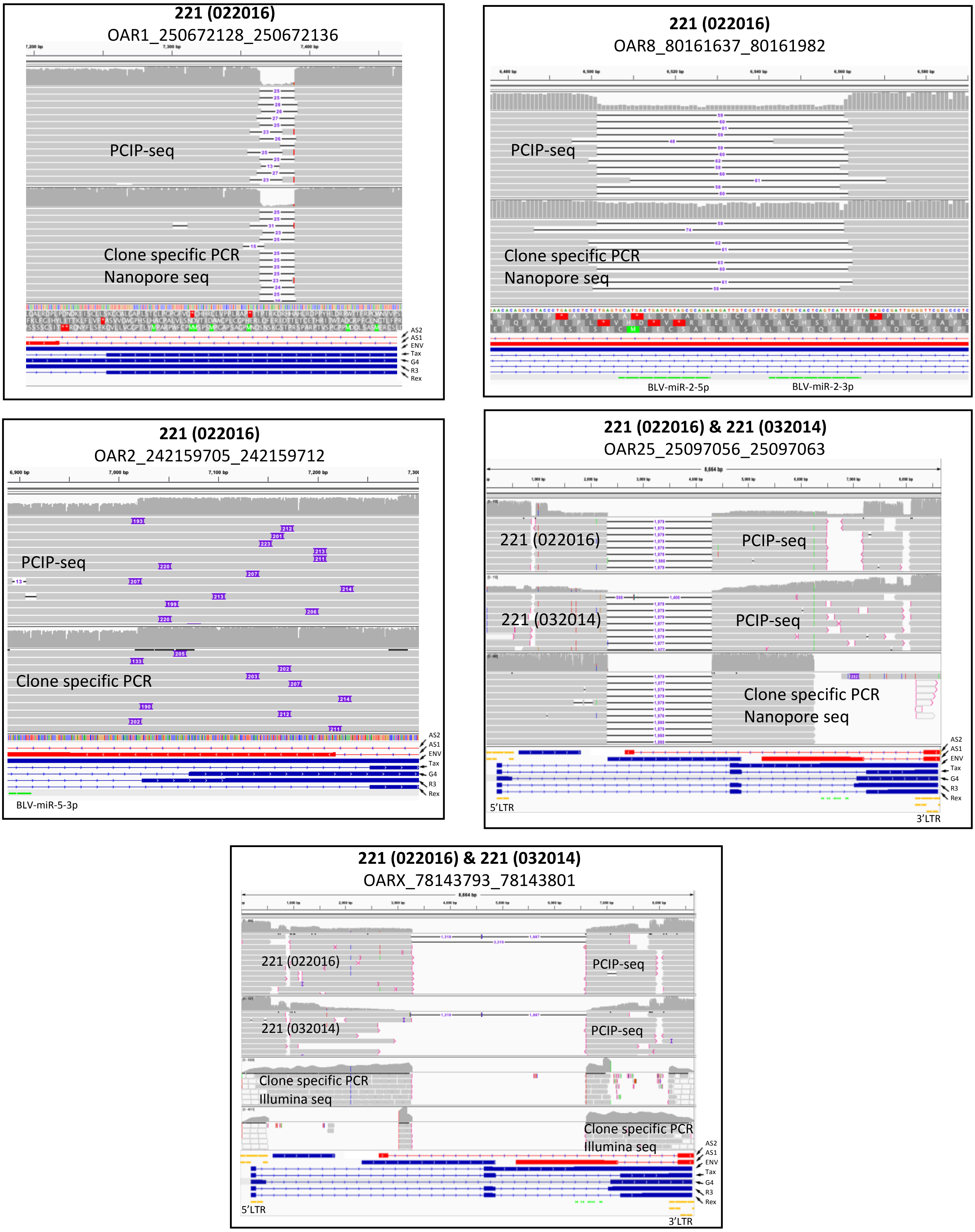

**Supplementary Figure 5** Clone specific PCR to validate BLV structural variants.

### Ovine 233
### BLV SVs validated by clone specific PCR

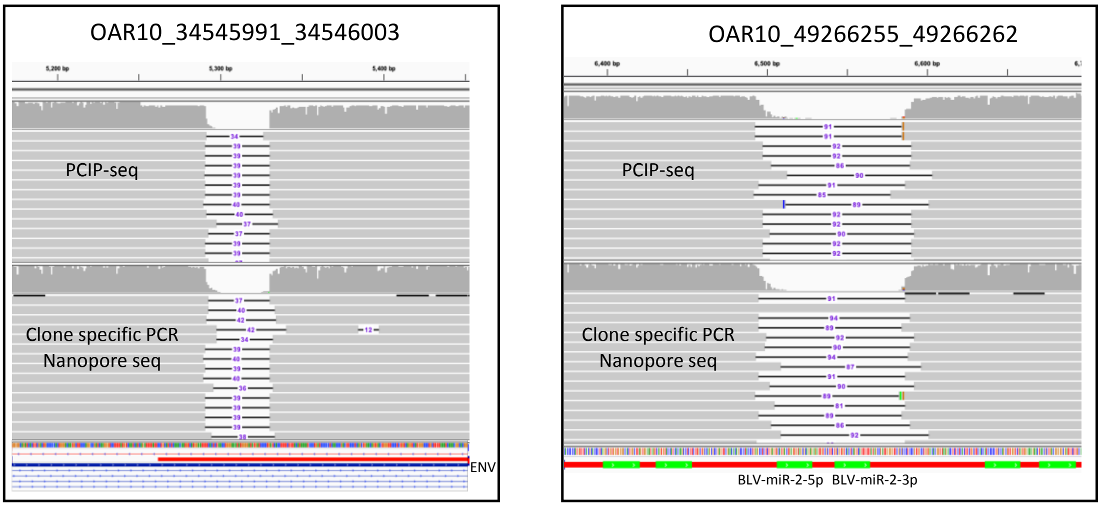

### Bovine 1439
### BLV SVs validated by clone specific PCR

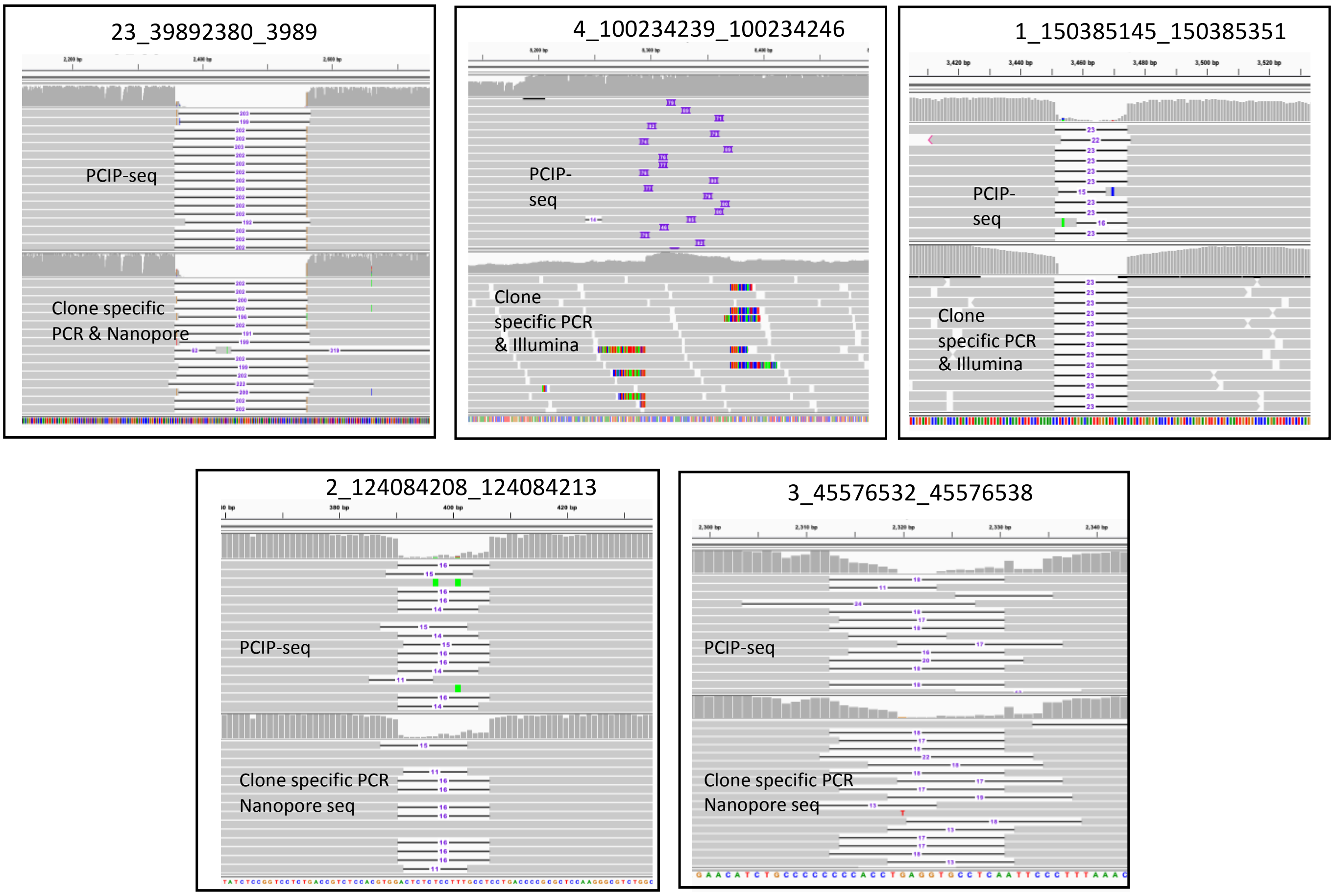

**Supplementary Figure 5 continued**

### Bovine 1439
### BLV 5’ deletions validated by clone specific PCR

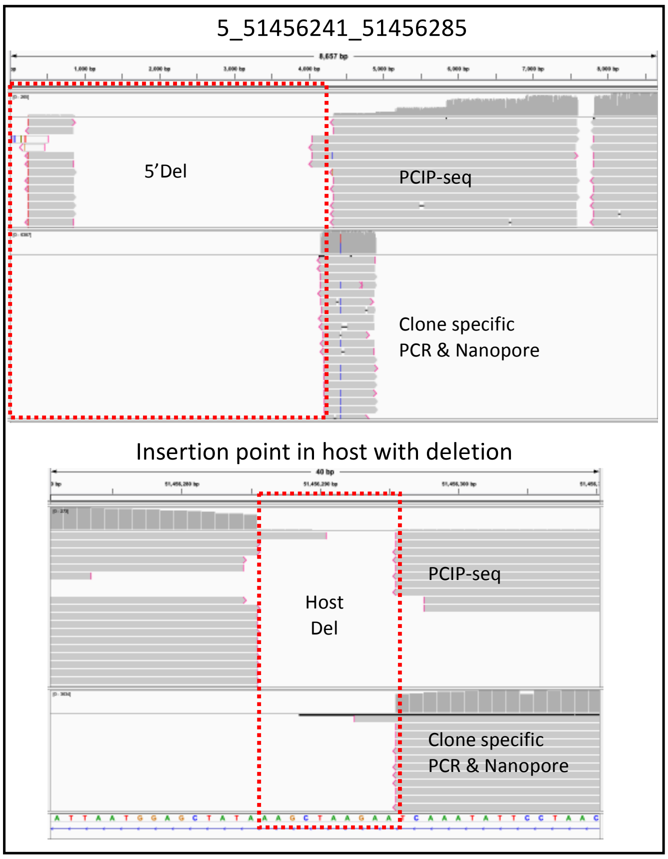

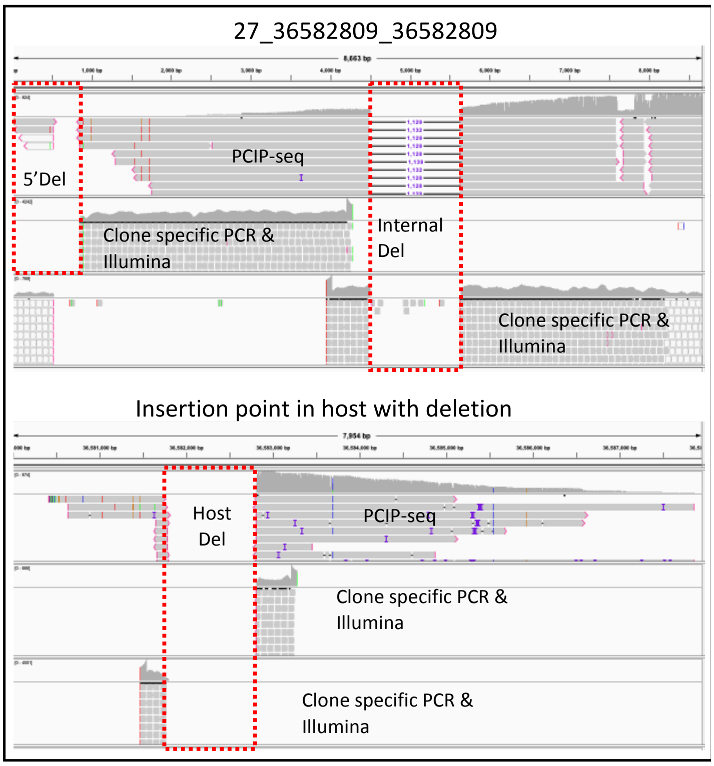

**Supplementary Figure 5 continued**

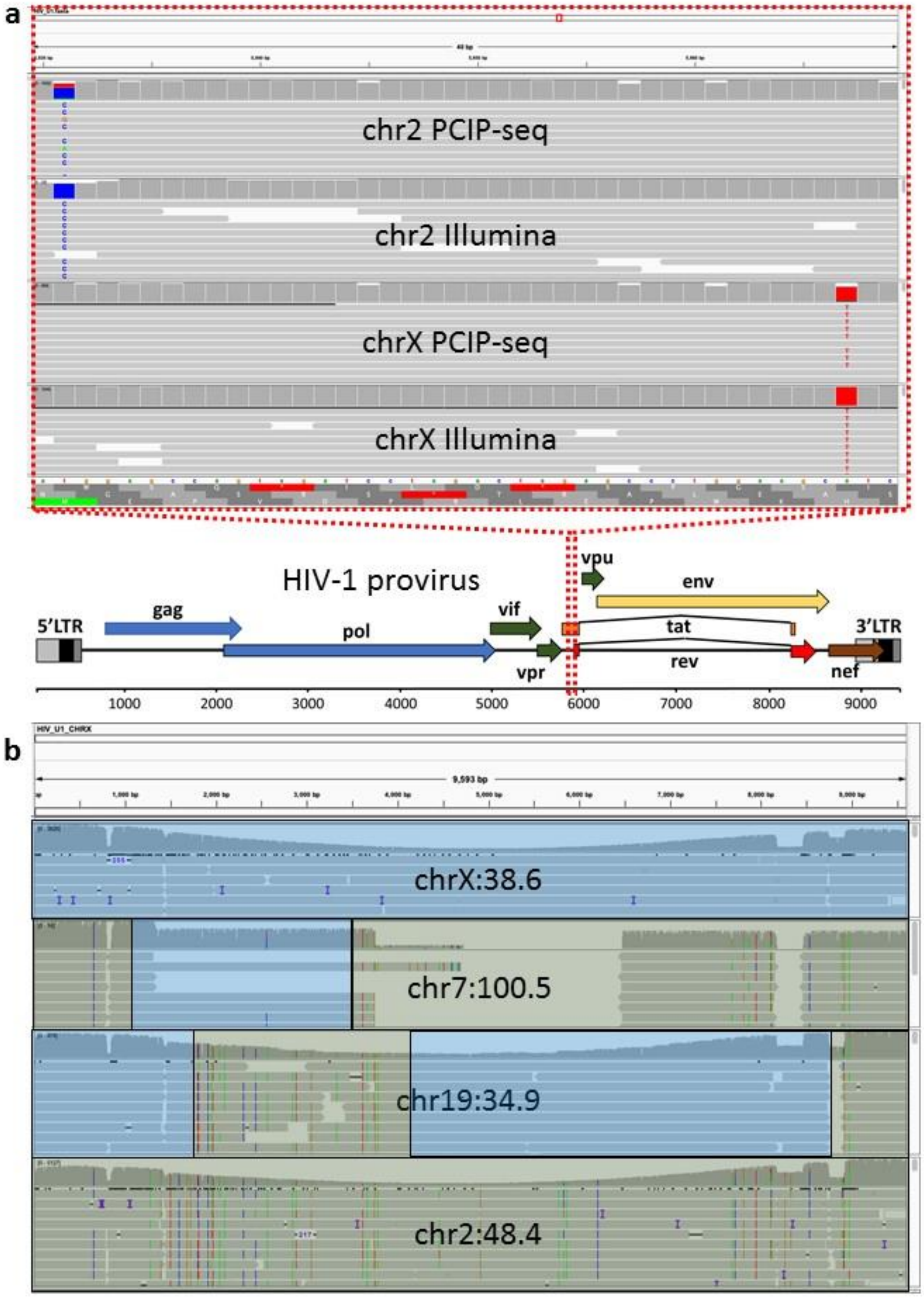

**Supplementary Figure 6** SNPs and recombination observed in the HIV-1 cell line U1 (**a**) Screen shot from IGV, representative PCIP-seq reads and clone specific PCR products sequenced on Illumina. This region corresponds to the first 13 amino acids of the Tat protein. In the chr2 provirus a T-to-C changes ATG to ACG and the first methionine to a threonine. In the chrX provirus an A-to-T changes CAT to CTT replacing a histidine at position 13 with a leucine. (**b**) Two proviruses, chr7:100.5 & chr19:34.9 identified as the products of recombination between major chrX and chr2 proviruses. IGV screen shot shows proviral reads from all four proviruses mapped to a full length proviral genome (the sequence of the chrX provirus was used as the reference). The colored vertical lines indicate SNPs and identify sequences derived from the chr2 provirus, highlighted in green. Sequences originating from the chrX provirus is highlighted in blue. Sequences of the provirus from chr19 and chr7 that match either chrX or chr2 provirus are highlighted in the appropriate color.

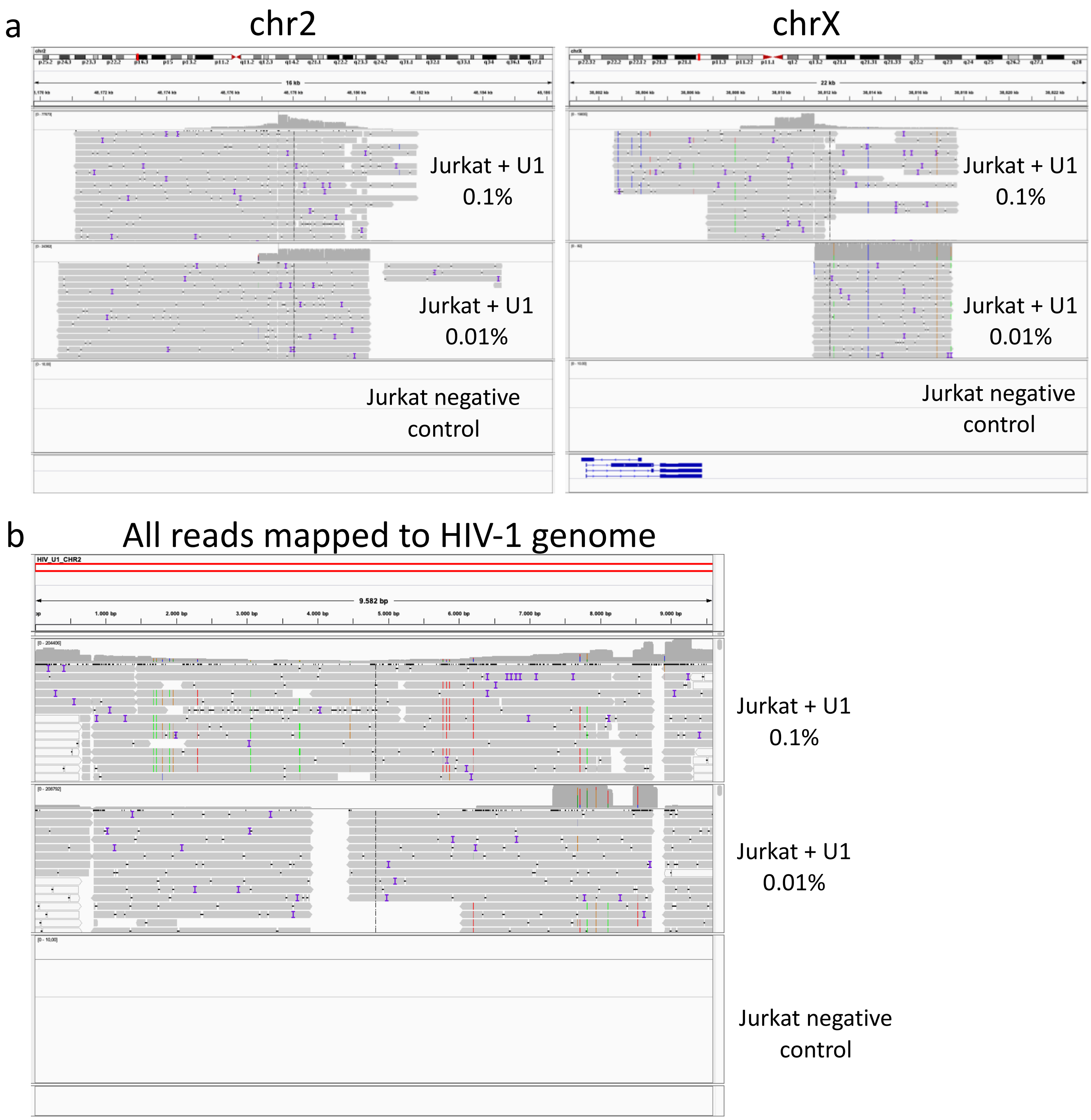

**Supplementary Figure 7** Three PCIP-seq libraries were prepared in parallel using 5 µg of template DNA, all used the same guides and primers. Following sequencing and demultiplexing the Jurkat negative control produced 12,137 reads, Jurkat + U1 0.01% produced 234,421 reads and Jurkat + U1 0.1% 252,913 reads. **(a)** The resultant reads were mapped to the human genome, the major integration sites observed in U1 on chr2 and chrX are shown. **(b)** The reads were also mapped the HIV-1 genome. No reads of pure HIV-1 or chimeric HIV-1/host reads were observed in the Jurkat negative control. In Jurkat + U1 0.01% samples 12.6% of the reads were chimeric HIV-1/host, in Jurkat + U1 0.1% this rose to 43.2%.

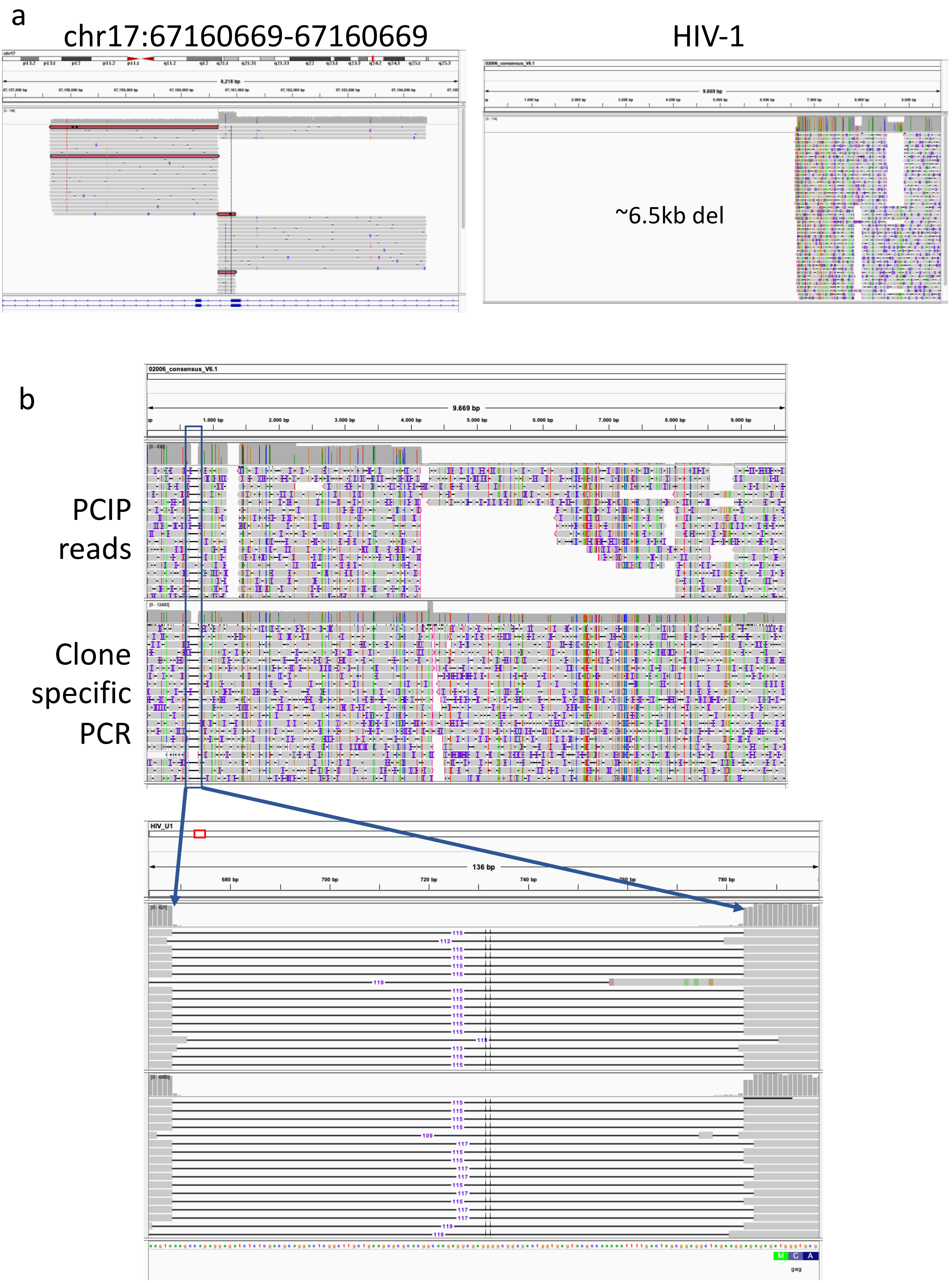

**Supplementary Figure 8** Deletions identified in the HIV-1 proviruses **(a)** Screen shot from IGV shows a provirus integration site in the host genome (patient 02006) as well as the associated provirus sequence. Two reads have been highlighted (out of many) that are found both upstream and downstream of the provirus integration site. With a full-length provirus this would not be possible, however, with a provirus carrying a large deletion including the 5’ or 3’ regions targeted by the guides a single read can encompass the truncated provirus as well as host DNA from both upstream and downstream of the integration. The provirus associated with this integration site has a 5’ deletion that removes ~6.5kb. **(b)** Provirus with a ~115 bp deletion affecting the region containing the packaging signal (Ψ). This provirus was also amplified via clone specific PCR, the elevated coverage towards the middle is where the two PCR products overlap.

### ERV insertion in the APOB gene

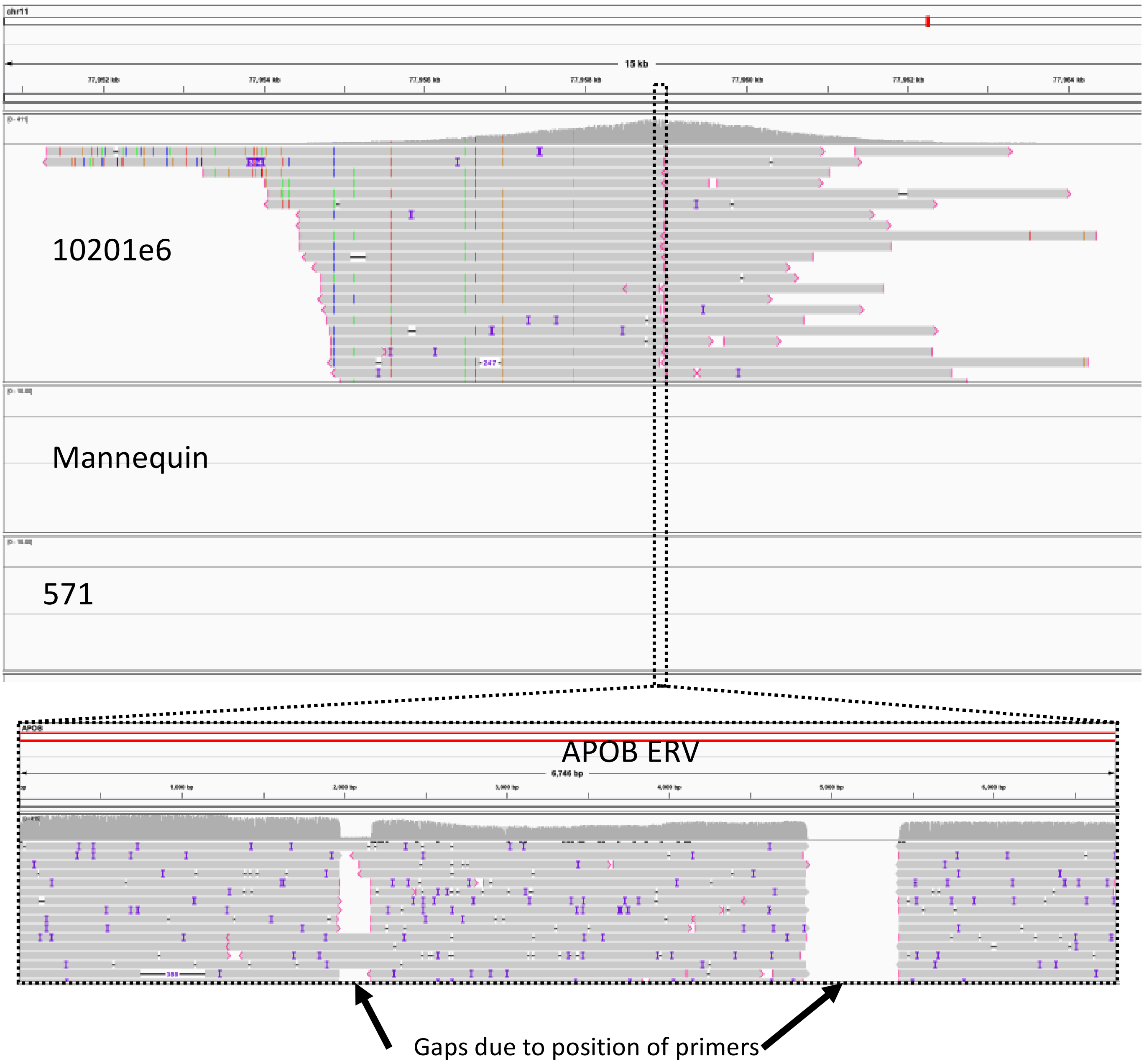

**Supplementary Figure 9** Screen capture from IGV: PCIP-seq identified the insertion site of the ERV responsible for cholesterol deficiency in Holstein cattle. No reads are seen mapping to this position in libraries from the other two cattle (Mannequin & 571). Below is shown the partial sequence of the provirus.

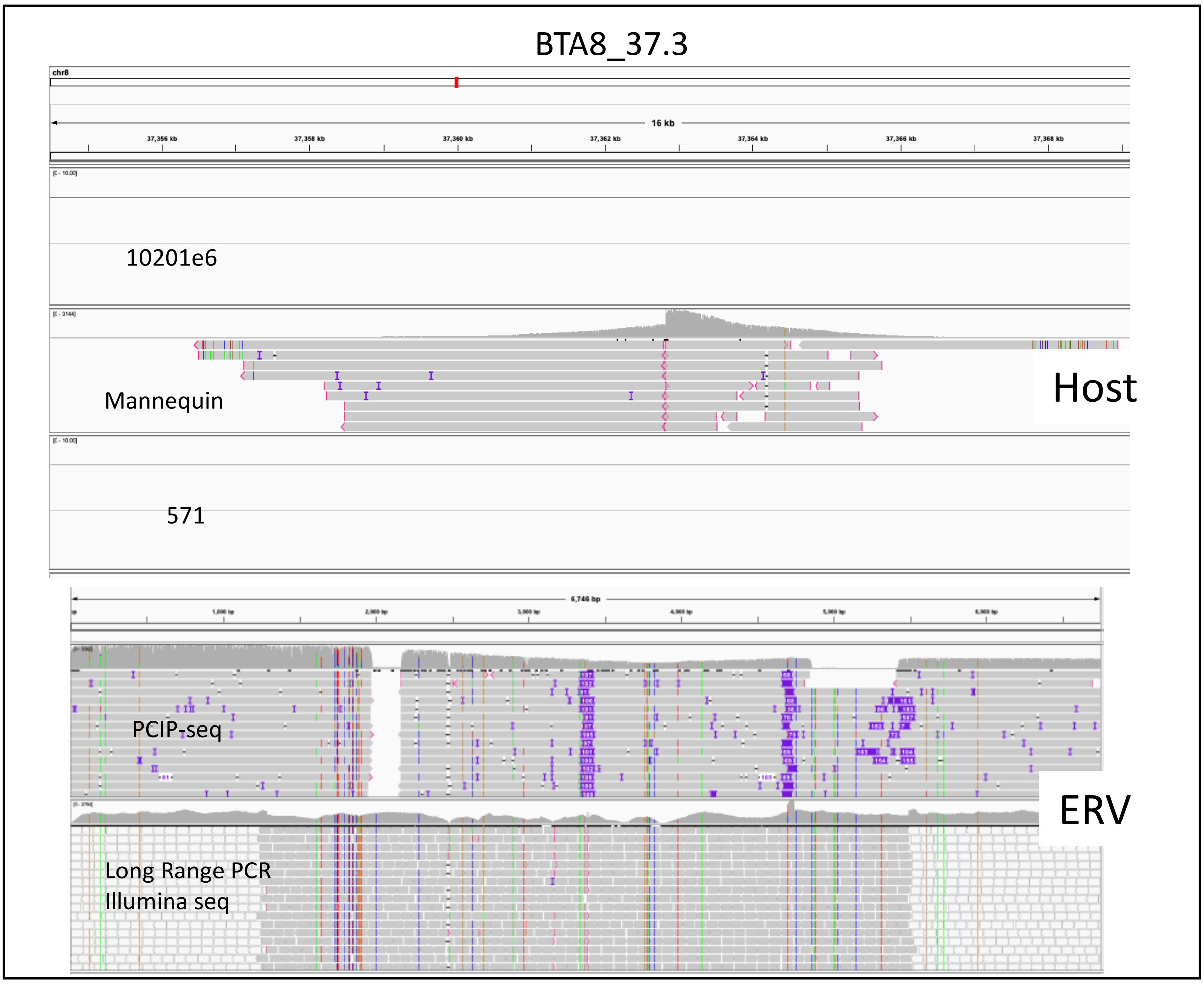

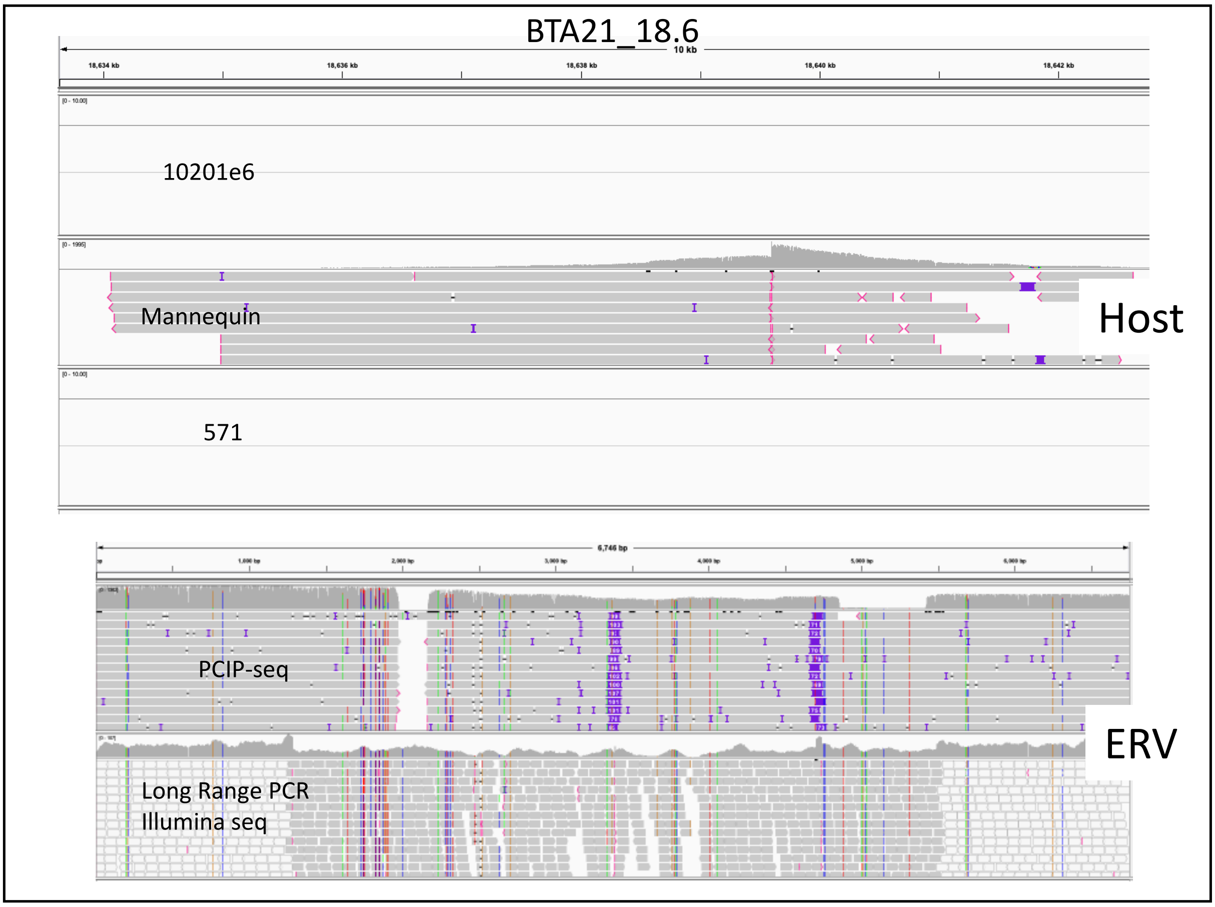

**Supplementary Figure 10** Validated, Bovine endogenous retrovirus (BERVK2) identified via PCIP-seq.

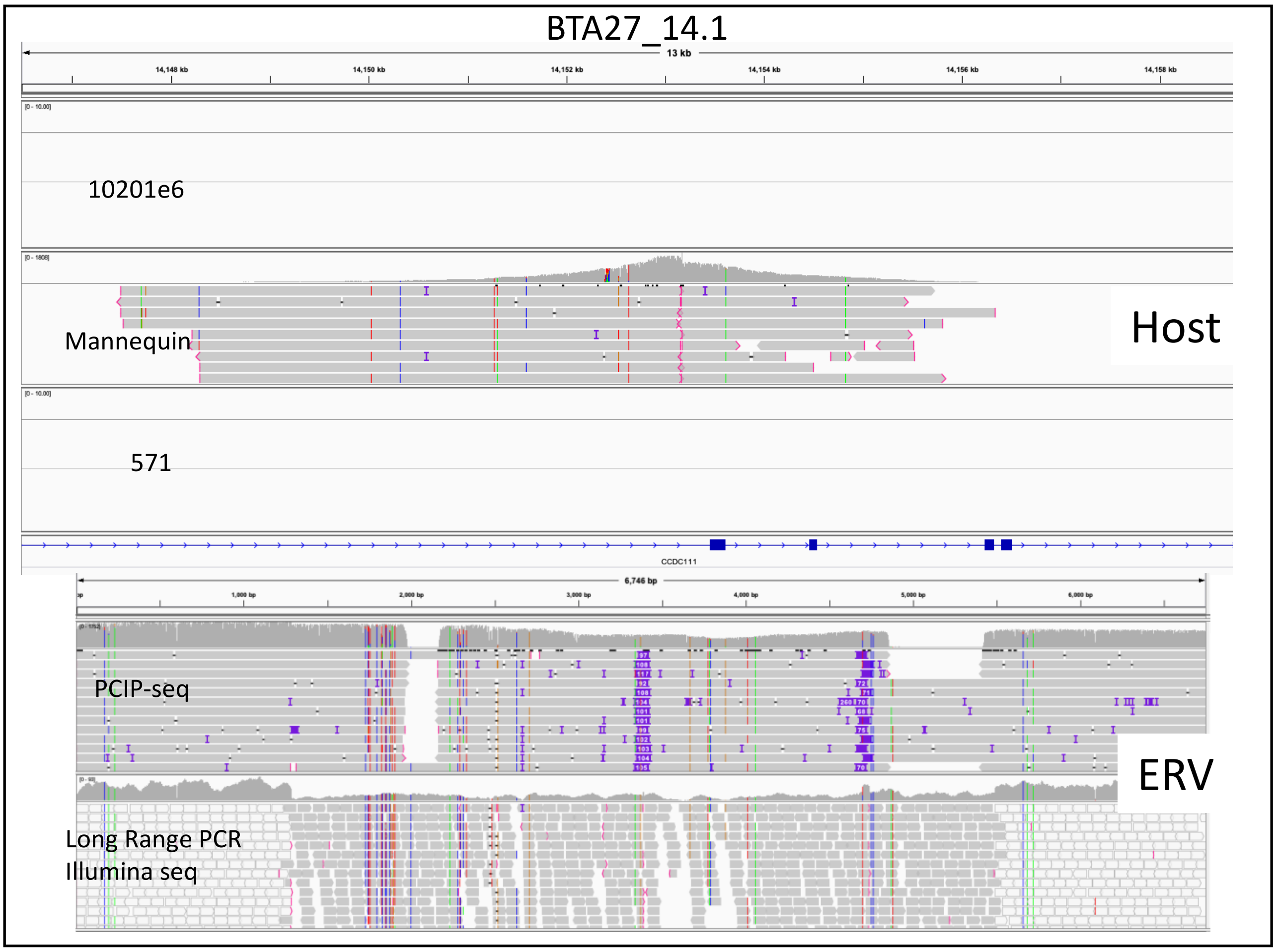

**Supplementary Figure 10 continued**

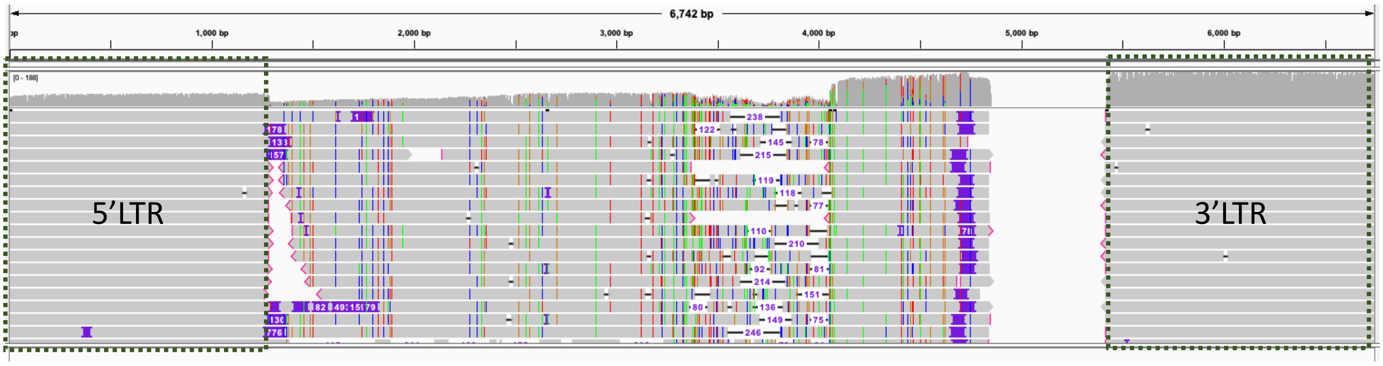

**Supplementary Figure 11** ERV BTA3_115.3 LTRs match APOB (BTA11_77.9) ERV.

a

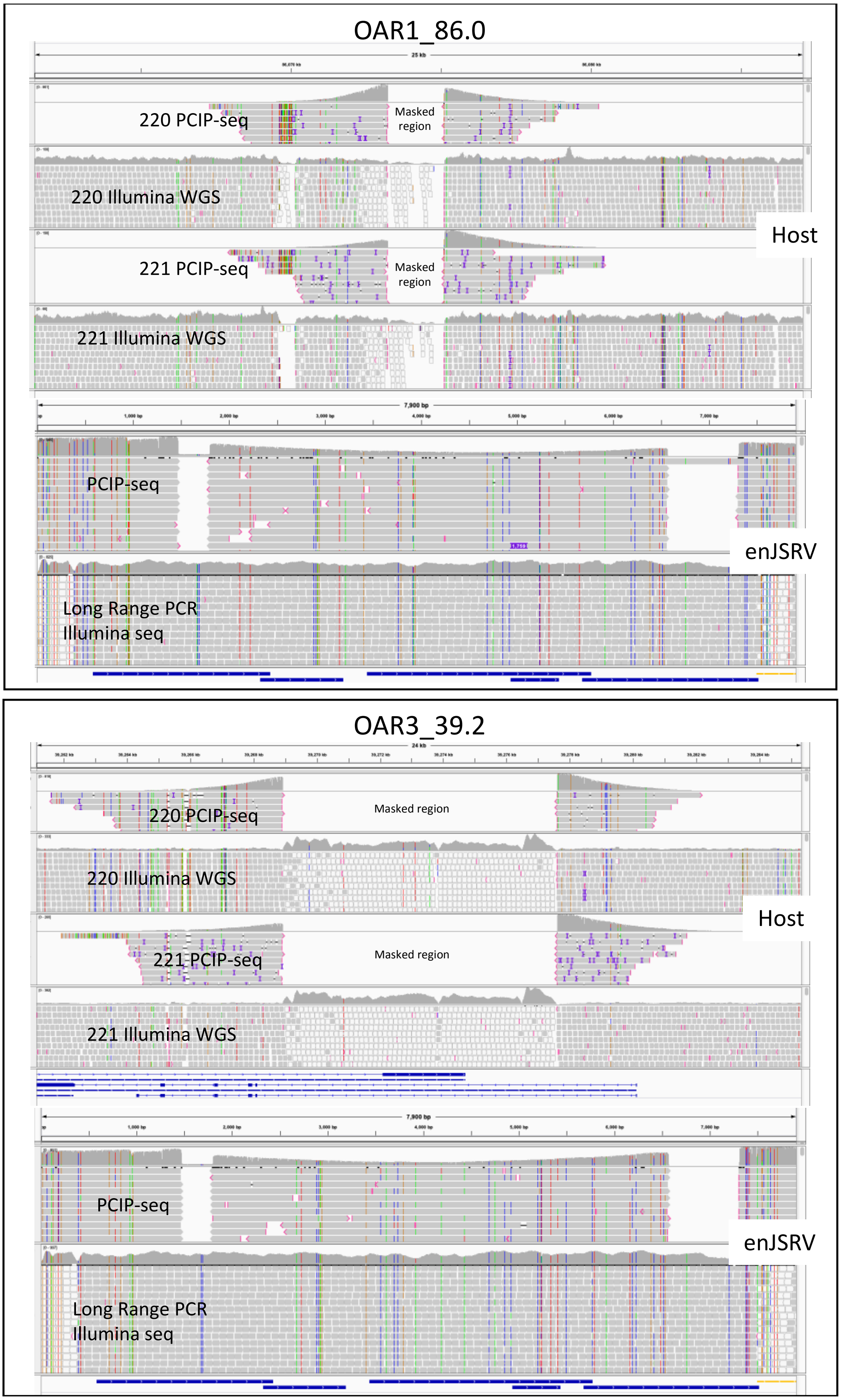

**Supplementary Figure 12** Validated, enJSRV **(a)** The PCIP-seq reads were mapped to the reference genome (OAR3) where sequences matching enJSRV had been masked out, this preventing reads from multiple proviruses mapping to these positions. Hybrid reads in the unique flanking sequence allowed us to determine the sequence of the proviruses present at these locations. **(b)** Evidence of enJSRV insertion was also observed in Illumina whole genome sequencing data from both animals. Colors, flag reads where one end is mapping to another region in the genome, pointing to an insertion at that position.

**b**

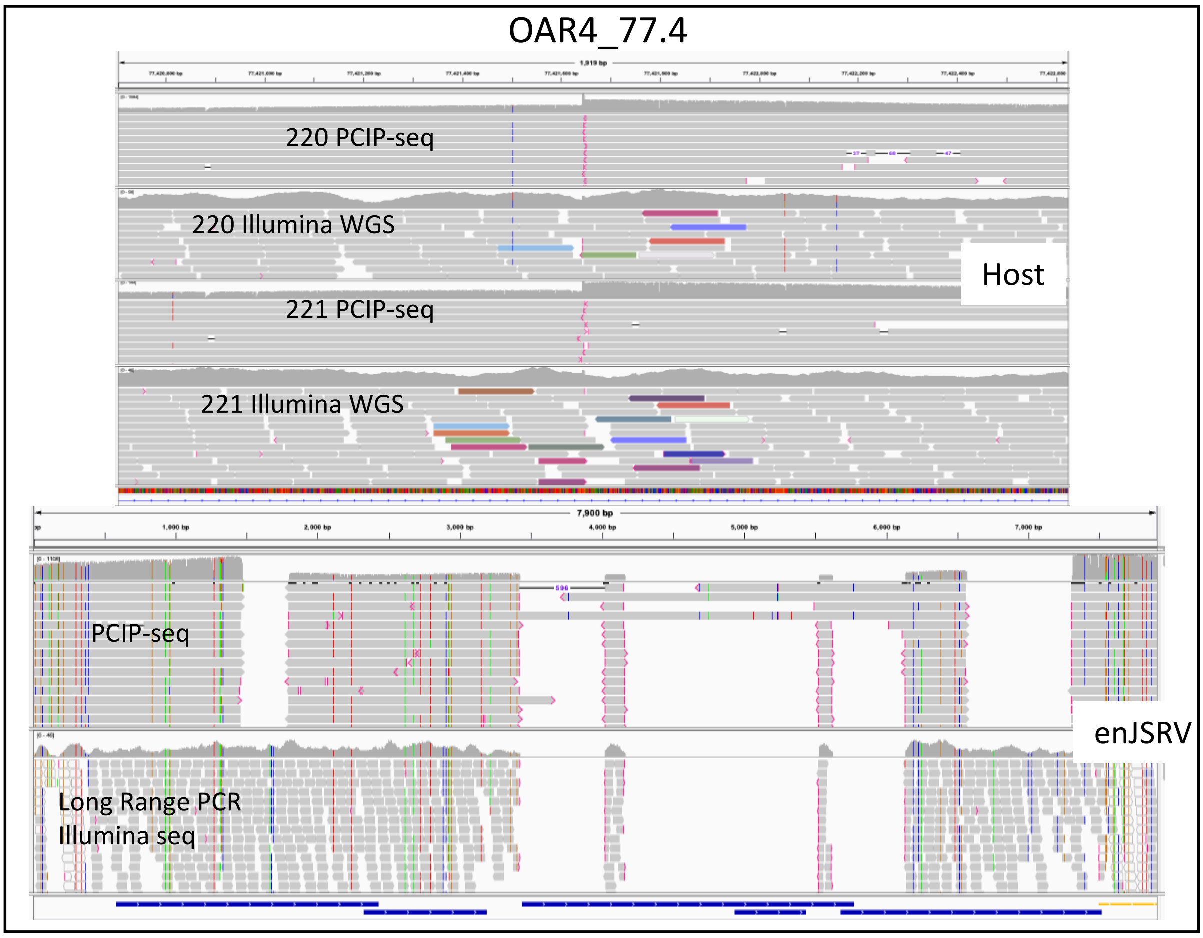

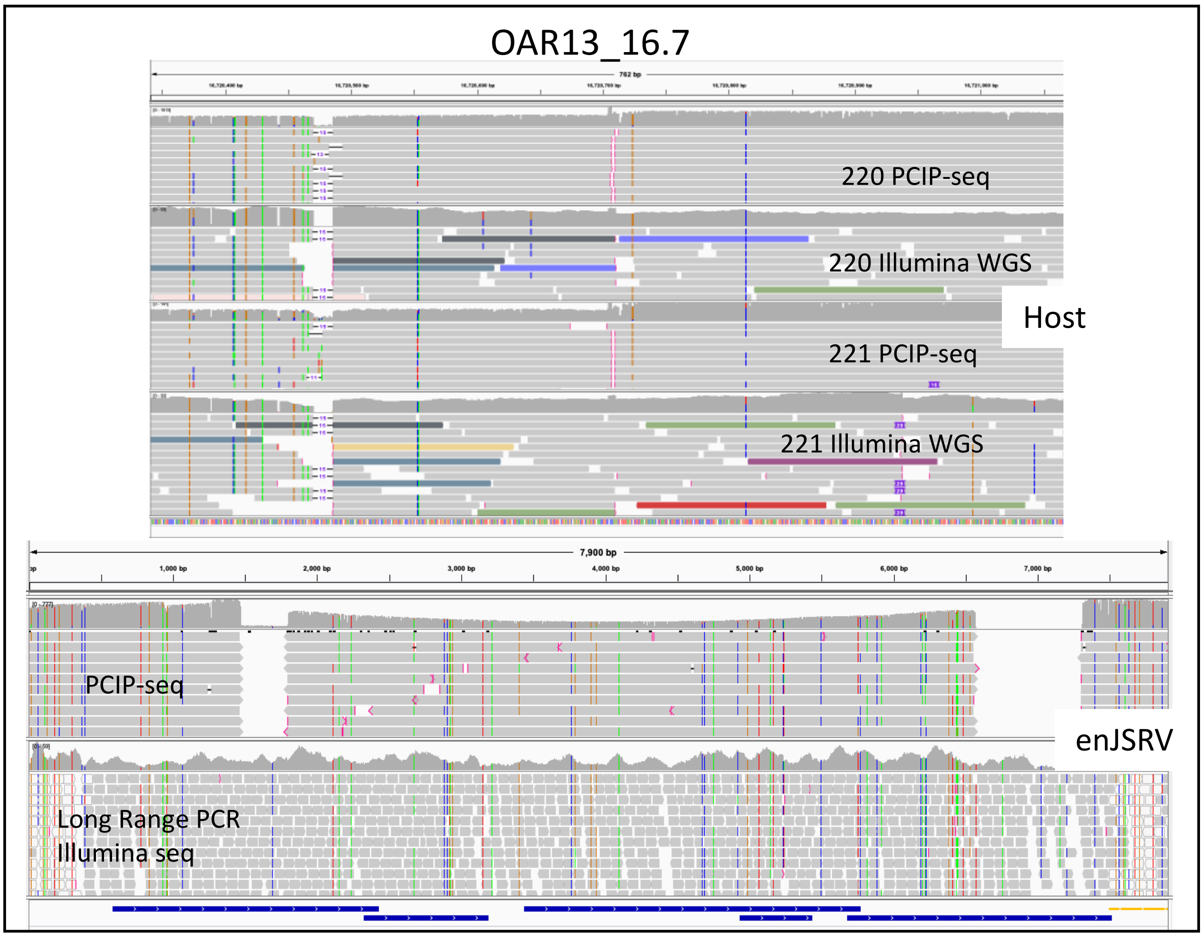

**Supplementary Figure 12 continued**

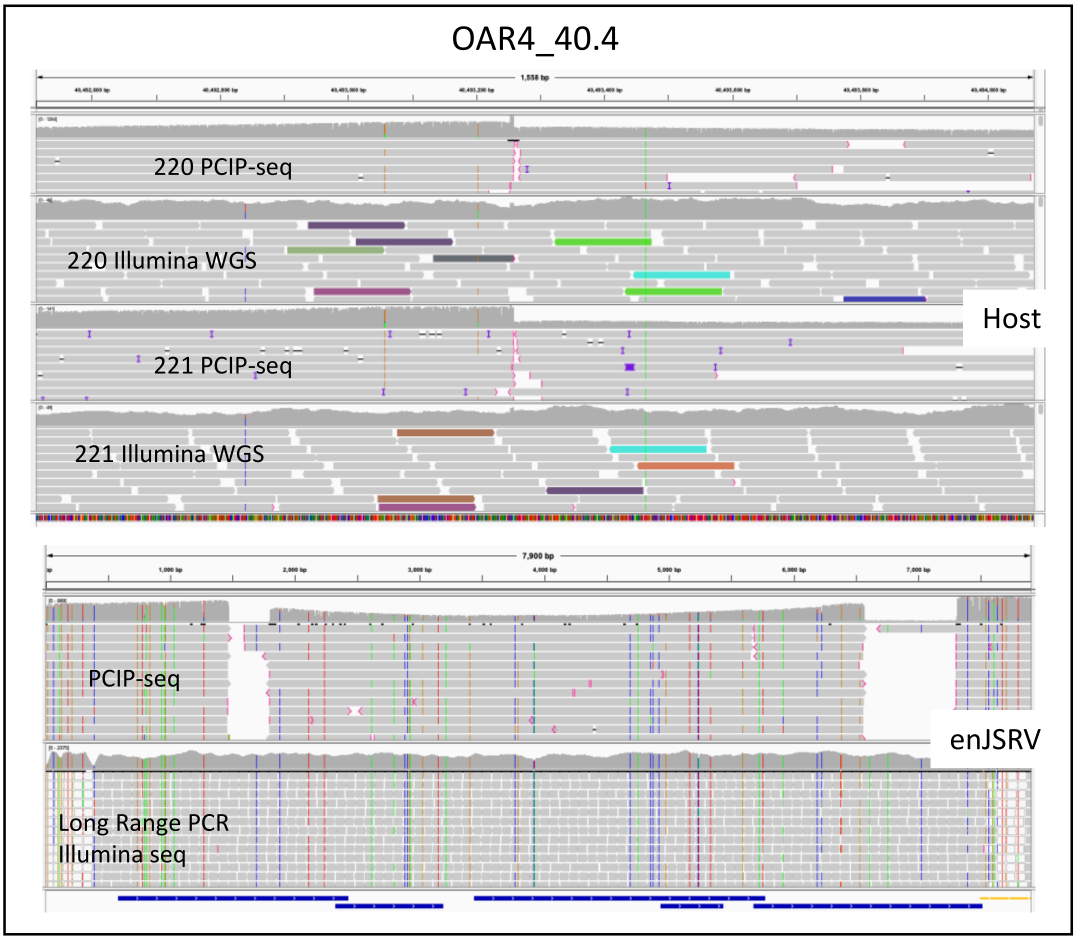

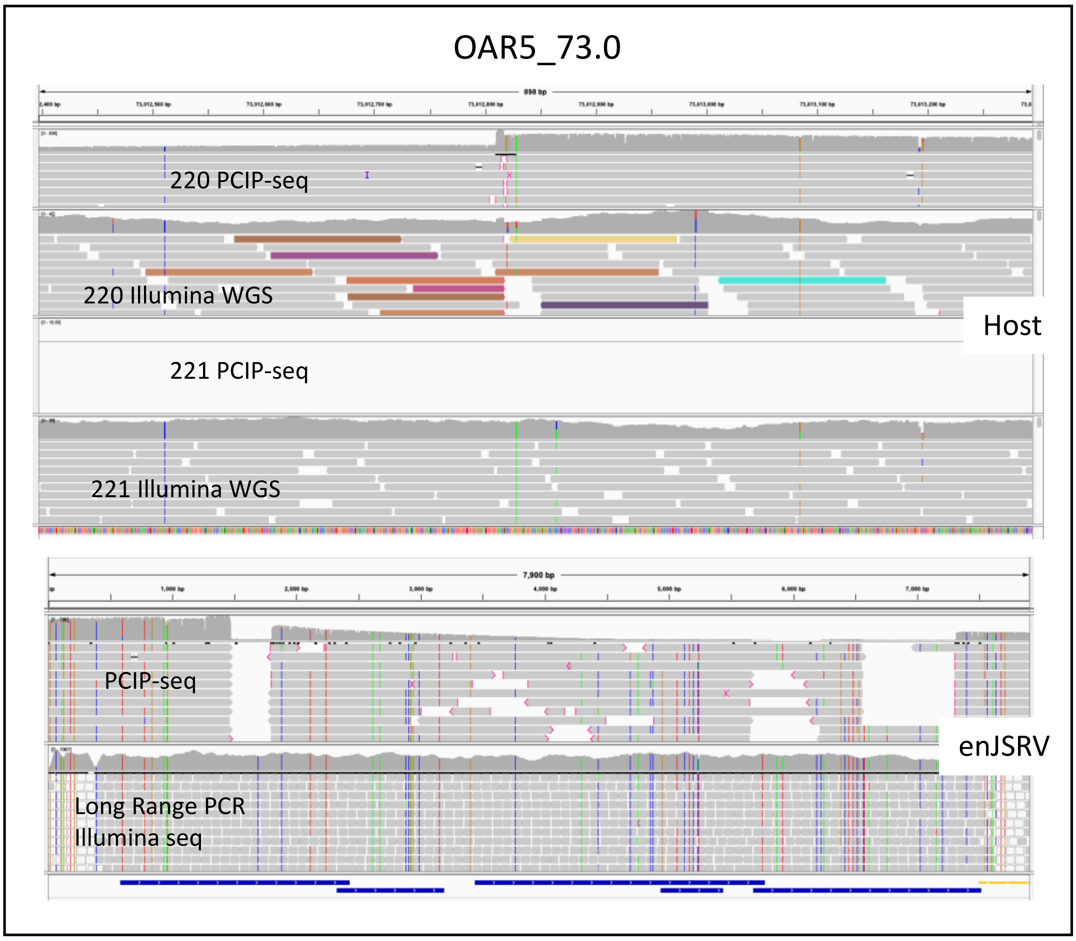

**Supplementary Figure 12 continued**

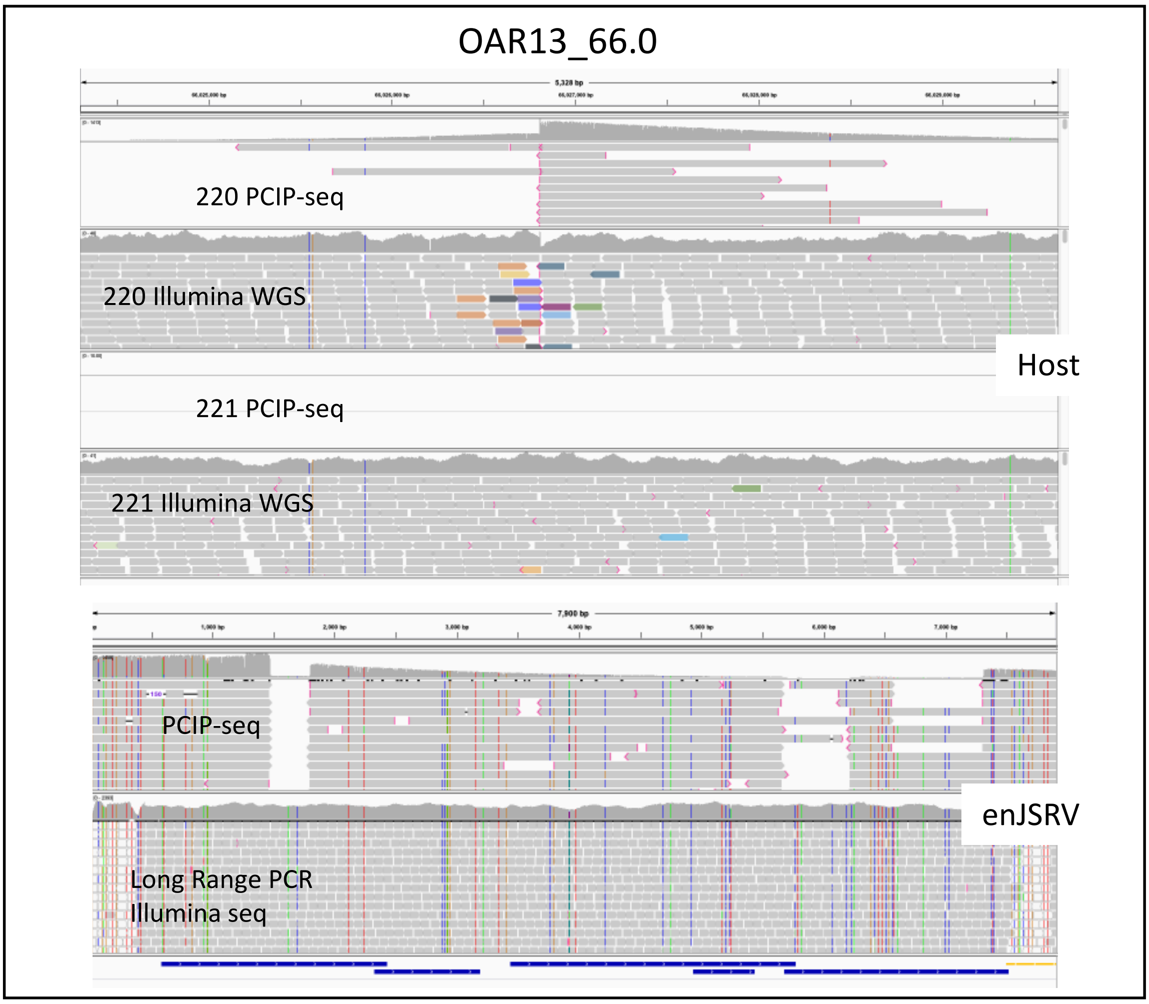

**Supplementary Figure 12 continued**

| Sample | Insertion sites ILLUMINA | Insertion sites PCIP-seq | U-IS ILL. in PCIP (%) | Pearson Correlation | Insertion sites ILLUMINA (>3) | Insertion sites PCIP-seq (>3) | U-IS ILL. in PCIP (%) (>3) | Raw PCIP-seq reads | Raw Illumina reads |
|---|---|---|---|---|---|---|---|---|---|
| 233 | 1110 | 5311 | 81.2 | 0.949810181 | 448 | 2302 | 85.9 | 524698 | 173196 |
| 221 (022016) | 1122 | 8023 | 40.4 | 0.511939213 | 74 | 3546 | 50 | 180276 | 9579 |
| 221 (032014) | 4473 | 5374 | 44.4 | 0.526457101 | 1555 | 1524 | 34.9 | 32266 | 391478 |
| 220 | 915 | 1352 | 36.1 | 0.894732877 | 401 | 664 | 47.6 | 44876 | 299554 |
| 1439 | 5784 | 5773 | 47.7 | 0.894732877 | 1449 | 3053 | 63.9 | 181055 | 216525 |
| 560 | 379 | 172 | 15.8 | 0.616804459 | 81 | 77 | 33.3 | 6802 | 192170 |
| 1053 | 8496 | 17903 | 62.0 | 0.811169919 | 2196 | 7777 | 68.5 | 367454 | 219461 |

**Supplementary Table 1** Comparing PCIP-seq to ligation mediated PCR and Illumina sequencing. For the Illumina libraries the template DNA used was 4 μg. For the PCIP-seq it varied between libraries (233=7μg, 221(022016)=4μg, 221(032014)=4μg, 220=2μg, 1439=3μg, 560=1μg, 1053=6μg). >3 signifies insertion sites supported by more than 3 reads after PCR duplicate removal. ILLUMINA = Ligation mediated PCR with Illumina sequencing. U-IS ILL. in PCIP = Unique insertion sites (%) identified in ILLUMINA and also found in PCIP-seq. Correlation Abundance Overlapping IS. Pearson's correlation Abundance = correlation of abundances from proviruses detected in both Illumina and PCIP-seq.

| Sample name | Species | PVL | # Insertion sites | # Proviruses examined for SNPs | # Variants detected (AF > 0.6) | # Proviruses with variant (AF > 0.6) | # Positions within proviruses with variant (AF > 0.6) |
|---|---|---|---|---|---|---|---|
| 233 | OAR | 78.3 | 5311 | 789 | 233 | 168 | 136 |
| 221 (022016) | OAR | 63.0 | 8023 | 408 | 93 | 79 | 86 |
| 221 (032014) | OAR | 16.0 | 5374 | 70 | 6 | 6 | 6 |
| 220 | OAR | 3.8 | 1352 | 130 | 50 | 42 | 36 |
| 1439 | BosT | 45.0 | 5773 | 587 | 311 | 211 | 137 |
| 1053 | BosT | 23.5 | 17903 | 1243 | 241 | 182 | 169 |

**Supplementary Table 2** Numbers of SNPs identified in each sample.

**1053**

| Provirus | Type | Region in BLV | Approx size | Clone specific PCR |
|---|---|---|---|---|
| 1_120275095_120275095 | DEL | 230-252 | 22 | no |
| 1_147862114_147862122 | DEL | 2241-2275 | 34 | no |
| 2_106933456_106933462 | DEL | 7674-7708 | 34 | no |
| 3_6970332_6970339 | DEL | 5109-6728 | 1619 | no |
| 3_90671155_90671163 | DEL | 2608-2919 | 311 | no |
| 4_114867583_114867589 | DEL | 4574-4637 | 63 | no |
| 5_25818093_25818100 | DEL | 4482-4526 | 44 | no |
| 6_95273607_95273614 | DEL | 4487-5537 | 1050 | no |
| 6_112133285_112133291 | DEL | 5217-5368 | 151 | no |
| 10_101509344_101509352 | DEL | 7324-7425 | 101 | no |
| 12_36183673_36183673 | DEL | 1808-1835 | 27 | no |
| 13_35328779_35328785 | DEL | 3679-4603 | 924 | no |
| 15_24605050_24605054 | DEL | 8136-8162 | 26 | no |
| 16_28380797_28380803 | DEL | 2984-3895 | 911 | no |
| 17_64277037_64277043 | DEL | 5418-5636 | 218 | no |
| 20_7882911_7882911 | DEL | 8111-8137 | 26 | no |
| 20_7882911_7882911 | DEL | 8230-8340 | 110 | no |
| 21_53434814_53434824 | DEL | 6854-7130 | 276 | no |
| 21_53434814_53434824 | DEL | 7202-7246 | 44 | no |
| 22_40343810_40343823 | DEL | 4629-4838 | 209 | no |
| 22_48239823_48239830 | DEL | 2271-2799 | 528 | no |
| 23_41760533_41760533 | DEL | 8100-8201 | 101 | no |
| 24_22643966_22643974 | DEL | 6857-7165 | 308 | no |
| 25_33749737_33749744 | DEL | 4225-4264 | 39 | no |
| 28_28470239_28470248 | DEL | 4496-5191 | 695 | no |
| 29_25146501_25146508 | DEL | 3901-5251 | 1350 | no |
| X_33071616_33071616 | DEL | 3322-3969 | 647 | no |
| X_61600607_61600612 | DEL | 6193-6783 | 590 | no |

**1439**

| Provirus | Type | Region in BLV | Approx size | Clone specific PCR |
|---|---|---|---|---|
| 10_65013091_65013093 | DEL | 2164-3192 | 1028 | no |
| 1_150385145_150385351 | DEL | 3451-3474 | 23 | yes |
| 2_121703720_121703726 | DEL | 5350-5399 | 49 | no |
| 23_39892380_39892560 | DEL | 2364-2560 | 196 | yes |
| 2_4188067_4188067 | DEL | 2176-2570 | 394 | no |
| 24_3748146_3748155 | DEL | 5419-5497 | 78 | no |
| 27_36582809_36582809 | DEL | 4522-5636 | 1114 | yes |
| 27_36582809_36582809 | DEL | 1-852 | 852 | yes |
| 4_100234239_100234246 | INS | 8296-8370 | 75 | yes |
| 5_51456241_51456285 | DEL | 1-4152 | 4152 | yes |
| 2_124084208_124084213 | DEL | 391-406 | 15 | yes |
| 3_45576532_45576538 | DEL | 2316-2336 | 20 | yes |
| 5_95348339_95348346 | DEL | 8167-8200 | 33 | no |
| 8_112613917_112613964 | DEL | 4225-6244 | 2019 | no |
| 5_6307451_6307451 | INS | 3251-3590 | 338 | no |

**221 (022016 & 032014)**

| Provirus | Type | Region in BLV | Approx size | Clone specific PCR |
|---|---|---|---|---|
| OAR3_128671913_128671921 | DEL | 4591-4620 | 30 | no |
| OAR18_26694984_26694991 | DEL | 5287-5508 | 222 | no |
| OAR25_25097056_25097063 | DEL | 2325-4303 | 1979 | yes |
| OARX_110727773_110727797 | DEL | 2858-2970 | 113 | no |
| OARX_78143793_78143801 | DEL | 3284-6602 | 3298 | yes |

**221 (032014)**

| Provirus | Type | Region in BLV | Approx size | Clone specific PCR |
|---|---|---|---|---|
| OAR14_25755878_25755884 | DEL | 5846-6486 | 640 | no |

**221 (022016)**

| Provirus | Type | Region in BLV | Approx size | Clone specific PCR |
|---|---|---|---|---|
| OAR1_25125478_25125485 | DEL | 6237-6255 | 19 | no |
| OAR1_250672128_250672136 | DEL | 7365-7389 | 25 | yes |
| OAR2_73878244_73878251 | DEL | 237-264 | 28 | no |
| OAR3_149619110_149619110 | DEL | 7610-7726 | 117 | no |
| OAR3_211678275_211678275 | DEL | 6228-6285 | 58 | no |
| OAR8_80161637_80161982 | DEL | 6502-6561 | 60 | yes |
| OAR13_10090846_10090865 | DEL | 6484-6561 | 78 | no |
| OAR16_10037623_10037623 | DEL | 1287-1396 | 110 | no |
| OAR21_31148897_31148902 | DEL | 7292-7544 | 253 | no |
| OAR24_28280610_28280610 | DEL | 6807-6828 | 22 | no |
| OAR2_242159705_242159712 | INS | 7017-7232 | 215 | yes |

**233**

| Provirus | Type | Region in BLV | Approx size | Clone specific PCR |
|---|---|---|---|---|
| OAR10_34545991_34546003 | DEL | 5298-5330 | 32 | yes |
| OAR10_49266255_49266262 | DEL | 6512-6586 | 74 | yes |
| OAR14_42146250_42146256 | DEL | 1658-1724 | 66 | no |
| OAR16_3998022_3998027 | DEL | 4479-4706 | 227 | no |
| OAR19_37466567_37466573 | DEL | 278-428 | 150 | no |
| OAR23_14140808_14140814 | DEL | 3270-5878 | 2608 | no |
| OAR3_184106381_184106391 | DEL | 5799-5874 | 75 | no |
| OAR7_72584331_72584331 | DEL | 4574-5453 | 879 | no |
| OAR7_72649090_72649098 | DEL | 539-629 | 90 | no |

**Supplementary Table 3** BLV structural variants identified via PCIP-seq.

| | 02006 | 06042 |
|---|---|---|
| Clinical characteristics | | |
| Age (years) | 52 | 47 |
| Gender | Male | Male |
| Time of diagnosis (year) | 2002 | 2006 |
| Viral load zenith (log10 HIV-1 c/ml) | 5.45 | 5.65 |
| CD4 count, nadir (cells/mm^3) | 44 | 118 |
| Start of ART (year) | 2002 | 2009 |
| Total ART duration (years) | 15 | 8 |
| Time of sampling (year) | 2017 | 2017 |
| CD4 count at sampling (cells/mm^3) | 293 | 659 |
| Viral load at sampling (HIV-1 c/ml) | <50 | <50 |
| CD4/CD8 ratio at sampling | 0.33 | 0.56 |
| Virological markers | | |
| Total HIV-1 DNA (c/10^6 CD4 T-cells) | 4644 | 5648 |

**Supplementary Table 4** Clinical information for the HIV-1 patients

| Patient | ID | strand | Approximate coverage at integration site | Clonal | Overlapping Gene | geneID | DELETIONS | HYPERMUT | VALIDATION | Notes |
|---|---|---|---|---|---|---|---|---|---|---|
| 02006 | chr1:1016083-1016083 | + | 48 | no | NA | NA | NA | NA | NA | |
| 02006 | chr1:19627829-19627829 | - | 768 | no | NBL1;MICOS10;MICOS10-NBL1 | ENSG00000158747;ENSG00000173436;ENSG00000270136 | LTR3 or LTR5 | NA | NA | |
| 02006 | chr1:26381880-26381880 | - | 108 | no | NA | NA | NA | NA | NA | |
| 02006 | chr1:26701728-26701728 | - | 115 | no | ARID1A | ENSG00000117713 | NA | NA | NA | |
| 02006 | chr1:27472247-27473034 | + | 2626 | no | WASF2 | ENSG00000158195 | LTR3 or LTR5 | NA | NA | |
| 02006 | chr1:32134188-32134188 | + | 15 | no | KPNA6 | ENSG00000025800 | NA | NA | NA | |
| 02006 | chr1:39004534-39004534 | + | 32 | no | AKIRIN1 | ENSG00000174574 | LTR3 | NA | NA | |
| 02006 | chr1:45853189-45853189 | + | 1770 | no | MAST2 | ENSG00000086015 | NA | NA | NA | |
| 02006 | chr1:52464788-52464788 | - | 7 | no | TUT4 | ENSG00000134744 | NA | NA | NA | |
| 02006 | chr1:70274742-70274743 | - | 43 | yes | ANKRD13C | ENSG00000118454 | LTR5 | NA | NA | |
| 02006 | chr1:87642624-87642624 | - | 65 | no | PKN2-AS1 | ENSG00000237505 | NA | NA | NA | |
| 02006 | chr1:89843535-89843535 | - | 159 | no | AC093423.3;LRRC8D | ENSG00000271949;ENSG00000171492 | LTR3 or LTR5 | NA | NA | |
| 02006 | chr1:120921873-120921874 | - | 884 | no | LINC00623 | ENSG00000226067 | NA | NA | NA | |
| 02006 | *chr1:143178830-143328870 | + | 2300 | yes | NA | NA | NA | NA | NA | **Full length intact provirus, in segmental duplication, flanking sequence maps to multiple positions** |
| 02006 | chr1:150113876-150113876 | + | 1 | no | VPS45 | ENSG00000136631 | NA | NA | NA | |
| 02006 | chr1:150381707-150381707 | + | 32 | yes | RPRD2 | ENSG00000163125 | INTERNAL | NA | NA | |
| 02006 | chr1:150862263-150862263 | + | 10 | no | ARNT | ENSG00000143437 | INTERNAL and LTR3 | NA | NA | |
| 02006 | chr1:186446839-186446839 | - | 18 | no | AL096803.2;PDC | ENSG00000229739;ENSG00000116703 | LTR3 | NA | NA | |
| 02006 | chr1:211963111-211963111 | - | 1575 | no | INTS7 | ENSG00000143493 | NA | NA | NA | |

| | | | | | | | | | |
|---|---|---|---|---|---|---|---|---|---|
| 02006 | chr1:225406245-225406245 | + | 672 | yes | LBR | ENSG00000143815 | NA | NA | NA |
| 02006 | chr2:9904677-9904678 | - | 3116 | no | TAF1B | ENSG00000115750 | LTR3 or LTR5 | NA | NA |
| 02006 | chr2:24631492-24631492 | + | 2635 | yes | NCOA1 | ENSG00000084676 | LTR3 | NA | NA |
| 02006 | chr2:42298830-42298830 | - | 6 | no | EML4 | ENSG00000143924 | LTR3 | NA | NA |
| 02006 | chr2:64718907-64718907 | + | 117 | no | SERTAD2 | ENSG00000179833 | NA | NA | NA |
| 02006 | chr2:89825679-89842825 | - | 19 | no | NA | NA | NA | NA | NA |
| 02006 | chr2:113929712-113929712 | - | 5 | no | ACTR3 | ENSG00000115091 | NA | NA | NA |
| 02006 | chr2:169986844-169986844 | + | 5 | no | UBR3 | ENSG00000144357 | NA | NA | NA |
| 02006 | chr2:174838060-174838060 | - | 234 | yes | CHN1 | ENSG00000128656 | LTR3 | NA | NA |
| 02006 | chr2:230450109-230450109 | - | 4 | no | SP100 | ENSG00000067066 | LTR5 | NA | NA |
| 02006 | chr2:230516833-230516834 | - | 2852 | no | SP100 | ENSG00000067066 | LTR3 or LTR5 | NA | NA |
| 02006 | chr2:232143921-232143921 | - | 1639 | no | DIS3L2 | ENSG00000144535 | NA | NA | NA |
| 02006 | chr2:237425257-237425257 | + | 103 | no | AC112721.1 | ENSG00000222022 | NA | NA | NA |
| 02006 | chr2:241325537-241325537 | - | 447 | no | SEPTIN2 | ENSG00000168385 | NA | NA | NA |
| 02006 | chr3:31537708-31537708 | - | 19 | no | STT3B | ENSG00000163527 | NA | NA | NA |
| 02006 | chr3:32444127-32444127 | + | 194 | no | CMTM7 | ENSG00000153551 | NA | NA | NA |
| 02006 | chr3:37369720-37369720 | - | 83 | no | NA | NA | NA | NA | NA |
| 02006 | chr3:45986908-45986908 | - | 377 | yes | FYCO1 | ENSG00000163820 | LTR3 | NA | NA |
| 02006 | chr3:47030997-47030997 | + | 85 | yes | SETD2 | ENSG00000181555 | NA | NA | NA |
| 02006 | chr3:56729144-56729144 | - | 44 | no | ARHGEF3 | ENSG00000163947 | NA | NA | NA |
| 02006 | chr3:106444741-106444741 | + | 176 | no | NA | NA | NA | NA | NA |
| 02006 | chr3:128197389-128197389 | - | 15 | no | EEFSEC | ENSG00000132394 | NA | TRUE | NA |
| 02006 | chr3:169788136-169788136 | + | 8 | no | MYNN | ENSG00000085274 | NA | NA | NA |
| 02006 | chr3:197755892-197755892 | - | 231 | no | FYTTD1 | ENSG00000122068 | NA | NA | NA |
| 02006 | chr4:160123-160123 | - | 310 | no | ZNF718 | ENSG00000250312 | NA | NA | NA |

| | | | | | | | | | | |
|---|---|---|---|---|---|---|---|---|---|---|
| 02006 | chr4:36218813-36218813 | - | 565 | no | ARAP2 | ENSG00000047365 | NA | NA | NA | |
| 02006 | chr4:49123990-49123990 | + | 15 | no | NA | NA | NA | NA | NA | Insertion point in satellite repeat |
| 02006 | chr4:54018653-54018653 | - | 26 | no | CHIC2;AC058822.1 | ENSG00000109220;ENSG00000282278 | NA | NA | NA | |
| 02006 | chr4:68576272-68576272 | - | 13 | no | NA | NA | LTR3 or LTR5 | NA | NA | |
| 02006 | chr4:76124574-76124581 | - | 2137 | yes | NUP54 | ENSG00000138750 | NA | NA | NA | |
| 02006 | chr4:125287928-125287928 | + | 382 | no | NA | NA | NA | NA | NA | |
| 02006 | chr5:35733674-35733674 | + | 1244 | no | SPEF2;AC137810.1 | ENSG00000152582;ENSG00000248969 | LTR3 or LTR5 | NA | NA | |
| 02006 | chr5:65963223-65963223 | - | 6 | no | ERBIN | ENSG00000112851 | LTR3 | NA | NA | |
| 02006 | chr5:98853970-98853970 | + | 1463 | no | NA | NA | NA | NA | NA | |
| 02006 | chr5:126413588-126413588 | + | 157 | no | AC093535.1;GRAMD2B | ENSG00000250602;ENSG00000155324 | NA | NA | NA | |
| 02006 | chr5:154798549-154798549 | + | 275 | yes | LARP1 | ENSG00000155506 | NA | TRUE | NA | |
| 02006 | chr6:30682000-30682001 | + | 1649 | no | PPP1R18 | ENSG00000146112 | NA | NA | NA | |
| 02006 | chr6:31284389-31284389 | - | 86 | no | HLA-B | ENSG00000234745 | NA | NA | NA | |
| 02006 | chr6:31466668-31466668 | - | 364 | no | HCP5 | ENSG00000206337 | NA | NA | NA | |
| 02006 | chr6:144749768-144749768 | - | 219 | no | UTRN | ENSG00000152818 | NA | NA | NA | |
| 02006 | chr6:149331149-149331150 | - | 791 | no | TAB2 | ENSG00000055208 | NA | NA | NA | |
| 02006 | chr6:159744428-159744428 | - | 5 | no | SOD2;WTAP;SOD2 | ENSG00000285441;ENSG00000146457;ENSG00000112096 | NA | NA | NA | |
| 02006 | chr7:44613211-44613211 | + | 70 | no | OGDH | ENSG00000105953 | NA | NA | NA | |
| 02006 | chr7:112863561-112863561 | - | 9 | no | BMT2;AC002463.1 | ENSG00000164603;ENSG00000223646 | NA | NA | NA | |
| 02006 | chr7:140846040-140846040 | - | 143 | yes | BRAF | ENSG00000157764 | LTR3 | NA | NA | |
| 02006 | chr8:26322239-26322239 | - | 354 | no | PPP2R2A | ENSG00000221914 | NA | NA | NA | |
| 02006 | chr8:143940947-143940947 | + | 3137 | no | PLEC | ENSG00000178209 | NA | NA | NA | |
| 02006 | chr9:20356269-20356269 | + | 345 | no | MLLT3 | ENSG00000171843 | NA | NA | NA | |

| | | | | | | | | | | |
|---|---|---|---|---|---|---|---|---|---|---|
| 02006 | chr9:20608766-20608771 | + | 1106 | yes | MLLT3 | ENSG00000171843 | NA | NA | LTR3 | |
| 02006 | chr9:97682561-97682561 | - | 1643 | no | XPA | ENSG00000136936 | LTR3 or LTR5 | NA | NA | |
| 02006 | chr9:97684177-97684177 | - | 1635 | no | XPA | ENSG00000136936 | NA | NA | NA | |
| 02006 | *chr10:41845251-42110306 | + | 2629 | yes | NA | NA | NA | NA | NA | **Full length intact provirus, in segmental duplication, flanking sequence maps to multiple positions (aka 02006_chr10_sc)** |
| 02006 | chr10:72182676-72182677 | + | 133 | no | ASCC1 | ENSG00000138303 | NA | NA | NA | |
| 02006 | chr10:102286149-102286149 | - | 1288 | no | GBF1 | ENSG00000107862 | LTR3 or LTR5 | NA | NA | |
| 02006 | chr10:119577391-119577407 | + | 8284 | yes | TIAL1 | ENSG00000151923 | LTR5 | NA | LTR5_LTR3 | |
| 02006 | chr10:119729370-119729370 | + | 2 | no | INPP5F | ENSG00000198825 | NA | NA | NA | |
| 02006 | chr11:34052095-34052095 | - | 45 | no | CAPRIN1;AC090469.1 | ENSG00000135387;ENSG00000286626 | LTR3 or LTR5 | NA | NA | |
| 02006 | chr11:65197842-65197842 | - | 8 | no | CAPN1 | ENSG00000014216 | LTR3 or LTR5 | NA | NA | |
| 02006 | chr11:66520477-66520477 | + | 10 | no | BBS1;AP002748.4 | ENSG00000174483;ENSG00000256349 | LTR3 or LTR5 | NA | NA | |
| 02006 | chr11:70227470-70227470 | - | 15 | no | NA | NA | NA | NA | NA | |
| 02006 | chr11:73917174-73917175 | + | 563 | no | PAAF1 | ENSG00000175575 | LTR3 or LTR5 | NA | NA | |
| 02006 | chr11:75330782-75330782 | + | 12 | no | ARRB1 | ENSG00000137486 | NA | NA | NA | |
| 02006 | chr11:86269413-86269419 | - | 3490 | yes | EED | ENSG00000074266 | NA | NA | NA | |
| 02006 | chr11:102384372-102384372 | - | 582 | no | NA | NA | LTR3 or LTR5 | NA | NA | |
| 02006 | chr11:107699310-107699310 | - | 579 | no | NA | NA | NA | NA | NA | |
| 02006 | chr11:119279088-119279144 | - | 11114 | yes | CBL | ENSG00000110395 | NA | NA | LTR3 | |
| 02006 | chr11:128226471-128226471 | - | 419 | yes | LINC02098 | ENSG00000272575 | LTR3 | NA | LTR5_LTR3 | |
| 02006 | chr12:56251024-56251024 | - | 872 | no | ANKRD52 | ENSG00000139645 | NA | NA | NA | |
| 02006 | chr12:111596412-111596412 | + | 3 | no | ATXN2 | ENSG00000204842 | NA | NA | NA | |
| 02006 | chr12:121828834-121828834 | - | 1 | no | SETD1B | ENSG00000139718 | NA | NA | NA | |

| | | | | | | | | | | |
|---|---|---|---|---|---|---|---|---|---|---|
| 02006 | chr13:16283554-17732159* | - | 465 | yes | NA | NA | NA | NA | NA | ***Full length intact provirus, in satellite repeat, matches centromere of chr13, chr14, chr21, chr22 (aka 02006_cen)** |
| 02006 | chr13:28217655-28217656 | + | 1297 | no | PAN3 | ENSG00000152520 | NA | NA | NA | |
| 02006 | chr13:41009612-41009613 | + | 264 | no | ELF1 | ENSG00000120690 | NA | NA | NA | |
| 02006 | chr13:98748841-98748841 | + | 365 | no | SLC15A1 | ENSG00000088386 | LTR3 or LTR5 | NA | NA | |
| 02006 | chr13:99234322-99234331 | + | 2338 | no | UBAC2 | ENSG00000134882 | NA | NA | NA | |
| 02006 | chr13:100302946-100302946 | + | 121 | no | PCCA | ENSG00000175198 | LTR3 | NA | NA | |
| 02006 | chr14:65416742-65416742 | - | 424 | no | FUT8 | ENSG00000033170 | NA | NA | NA | |
| 02006 | chr14:76188763-76188764 | + | 526 | no | GPATCH2L | ENSG00000089916 | NA | NA | NA | |
| 02006 | chr14:106140348-106140348 | - | 7 | no | NA | NA | LTR3 | NA | NA | |
| 02006 | chr15:43821570-43821571 | - | 2678 | yes | MFAP1 | ENSG00000140259 | LTR3 or LTR5 | NA | NA | |
| 02006 | chr15:49000157-49000157 | + | 232 | no | SECISBP2L | ENSG00000138593 | NA | NA | NA | |
| 02006 | chr15:76648943-76648943 | + | 1475 | yes | SCAPER | ENSG00000140386 | LTR3 | NA | NA | |
| 02006 | chr16:1750631-1750631 | + | 1166 | yes | MAPK8IP3 | ENSG00000138834 | LTR3 | NA | NA | |
| 02006 | chr16:4491521-4491521 | - | 226 | no | NMRAL1;HMOX2 | ENSG00000153406;ENSG00000103415 | NA | NA | NA | |
| 02006 | chr16:29362672-29362679 | - | 1033 | yes | AC025279.1;SNX29P2 | ENSG00000198106;ENSG00000271699 | 115bp internal del | NA | LTR5_LTR3 | **Full length provirus with 115bp del** |
| 02006 | chr16:29433119-29433119 | + | 357 | no | SMG1P6 | ENSG00000254634 | NA | NA | NA | Insertion point in segmental duplication, also maps to: chr16:29534883-29534884 & chr16:30274869-30274869 |
| 02006 | chr16:29527594-29527594 | + | 11 | no | SMG1P2 | ENSG00000205534 | NA | NA | NA | |
| 02006 | chr16:30156074-30156074 | - | 62 | no | NA | NA | NA | NA | NA | |
| 02006 | chr16:30267583-30267583 | + | 11 | no | SMG1P5 | ENSG00000183604 | NA | NA | NA | |
| 02006 | chr16:31733056-31733056 | - | 226 | no | ZNF720 | ENSG00000197302 | NA | NA | NA | |
| 02006 | chr16:47597471-47597471 | + | 11 | no | PHKB | ENSG00000102893 | LTR3 | NA | NA | |
| 02006 | chr16:57218299-57218299 | - | 2 | no | RSPRY1 | ENSG00000159579 | NA | NA | NA | |

| | | | | | | | | | |
|---|---|---|---|---|---|---|---|---|---|
| 02006 | chr16:66882446-66882446 | + | 118 | no | PDP2 | ENSG00000172840 | NA | NA | NA |
| 02006 | chr16:67037627-67037627 | + | 263 | no | CBFB | ENSG00000067955 | LTR3 or LTR5 | NA | NA |
| 02006 | chr16:68090945-68090946 | - | 2113 | no | NFATC3;AC130462.1 | ENSG00000072736;ENSG00000261864 | NA | NA | NA |
| 02006 | chr16:78079160-78079161 | - | 1052 | no | NA | NA | LTR3 | NA | NA |
| 02006 | chr16:81328023-81328023 | + | 79 | no | GAN | ENSG00000261609 | NA | NA | NA |
| 02006 | chr16:85885591-85885591 | + | 116 | no | NA | NA | NA | NA | NA |
| 02006 | chr17:4289609-4289609 | + | 64 | no | UBE2G1 | ENSG00000132388 | NA | NA | NA |
| 02006 | chr17:5346123-5346123 | + | 4 | no | RABEP1 | ENSG00000029725 | NA | NA | NA |
| 02006 | chr17:30406481-30406481 | - | 11 | no | CPD | ENSG00000108582 | NA | NA | NA |
| 02006 | chr17:30509437-30509437 | - | 444 | no | GOSR1 | ENSG00000108587 | NA | NA | NA |
| 02006 | chr17:37453838-37453839 | + | 517 | no | TADA2A | ENSG00000276234 | NA | NA | NA |
| 02006 | chr17:40240347-40240348 | - | 1555 | no | WIPF2 | ENSG00000171475 | NA | NA | NA |
| 02006 | chr17:42244762-42244762 | - | 233 | yes | STAT5B | ENSG00000173757 | LTR3 | NA | NA |
| 02006 | chr17:42252594-42252594 | - | 477 | no | STAT5B | ENSG00000173757 | NA | NA | NA |
| 02006 | chr17:42254307-42254307 | - | 62 | no | STAT5B | ENSG00000173757 | NA | NA | NA |
| 02006 | chr17:42270841-42270841 | - | 59 | no | STAT5B | ENSG00000173757 | NA | NA | NA |
| 02006 | chr17:47182967-47182967 | + | 908 | no | CDC27 | ENSG00000004897 | LTR3 | NA | NA |
| 02006 | chr17:64515517-64515517 | - | 483 | yes | CEP95 | ENSG00000258890 | LTR3 or LTR5 | NA | NA |
| 02006 | chr17:67160669-67160669 | + | 93 | yes | HELZ | ENSG00000198265 | LTR5 | NA | NA |
| 02006 | chr17:80677360-80677360 | - | 5 | no | RPTOR | ENSG00000141564 | NA | NA | NA |
| 02006 | chr18:58746310-58746310 | + | 108 | no | AC104365.1;MALT1 | ENSG00000267476;ENSG00000172175 | NA | NA | NA |
| 02006 | chr19:2106957-2106957 | + | 124 | no | AP3D1 | ENSG00000065000 | NA | NA | NA |
| 02006 | chr19:2789049-2789049 | - | 42 | no | AC006538.1 | ENSG00000172009 | NA | NA | NA |
| 02006 | chr19:8447846-8447847 | - | 1506 | no | HNRNPM | ENSG00000099783 | LTR5 | NA | NA |
| 02006 | chr19:8468181-8468181 | - | 23 | no | HNRNPM | ENSG00000099783 | LTR3 or LTR5 | NA | NA |

| | | | | | | | | | |
|---|---|---|---|---|---|---|---|---|---|
| 02006 | chr19:9146101-9146101 | - | 1085 | no | ZNF317 | ENSG00000130803 | NA | NA | NA |
| 02006 | chr19:10132512-10132512 | + | 215 | no | NA | NA | LTR3 | NA | NA |
| 02006 | chr19:13967475-13967475 | + | 45 | no | RFX1 | ENSG00000132005 | NA | NA | NA |
| 02006 | chr19:15392131-15392131 | + | 76 | yes | AKAP8L | ENSG00000011243 | NA | NA | NA |
| 02006 | chr19:19012411-19012411 | + | 50 | no | SUGP2 | ENSG00000064607 | NA | NA | NA |
| 02006 | chr19:35224061-35224061 | + | 167 | no | NA | NA | NA | NA | NA |
| 02006 | chr19:43570080-43570080 | - | 25 | no | XRCC1;L34079.1 | ENSG00000073050;ENSG00000268361 | LTR3 | NA | NA |
| 02006 | chr19:43988330-43988330 | - | 22 | no | ZNF155 | ENSG00000204920 | NA | NA | NA |
| 02006 | chr19:45044823-45044824 | - | 184 | no | CLASRP | ENSG00000104859 | NA | NA | NA |
| 02006 | chr19:46931657-46931657 | + | 754 | no | ARHGAP35 | ENSG00000160007 | NA | NA | NA |
| 02006 | chr20:2115584-2115584 | - | 1465 | no | STK35 | ENSG00000125834 | LTR3 or LTR5 | NA | NA |
| 02006 | chr20:36921829-36921830 | + | 3341 | yes | SAMHD1 | ENSG00000101347 | NA | NA | NA |
| 02006 | chr21:39196370-39196370 | + | 1548 | no | BRWD1 | ENSG00000185658 | NA | NA | NA |
| 02006 | chr21:46656339-46656339 | + | 442 | no | PRMT2 | ENSG00000160310 | NA | TRUE | NA |
| 02006 | chr22:31231065-31231066 | + | 394 | no | LIMK2 | ENSG00000182541 | LTR3 or LTR5 | NA | NA |
| 02006 | chr22:36347896-36347896 | + | 16 | no | MYH9 | ENSG00000100345 | NA | NA | NA |
| 02006 | chr22:37144529-37144529 | - | 233 | no | IL2RB | ENSG00000100385 | NA | NA | NA |
| 02006 | chr22:40543701-40543701 | - | 847 | no | MRTFA | ENSG00000196588 | NA | NA | NA |
| 02006 | chr22:40585152-40585152 | - | 33 | no | MRTFA | ENSG00000196588 | LTR3 | NA | NA |
| 02006 | chr22:49862648-49862648 | - | 3820 | yes | ZBED4 | ENSG00000100426 | LTR3 | NA | NA |
| 02006 | chrX:37845020-37845020 | + | 16 | no | AF241726.2;DYNLT3 | ENSG00000250349;ENSG00000165169 | LTR3 or LTR5 | NA | NA |
| 02006 | chrX:136682979-136682979 | + | 1942 | no | ARHGEF6 | ENSG00000129675 | LTR3 or LTR5 | NA | NA |
| 02006 | chrX:153728151-153728151 | + | 13 | no | ABCD1 | ENSG00000101986 | NA | NA | NA |
| 06042 | chr1:25255059-25255059 | - | 262 | no | RSRP1 | ENSG00000117616 | LTR3 or LTR5 | NA | NA |
| 06042 | chr1:155900488-155900489 | + | 1028 | no | RIT1 | ENSG00000143622 | LTR3 or LTR5 | NA | NA |

| | | | | | | | | | |
|---|---|---|---|---|---|---|---|---|---|
| 06042 | chr1:156312305-156312305 | - | 472 | no | CCT3 | ENSG00000163468 | NA | TRUE | NA |
| 06042 | chr2:61290670-61290671 | - | 399 | no | USP34 | ENSG00000115464 | NA | NA | NA |
| 06042 | chr2:109140922-109140922 | - | 176 | no | SH3RF3 | ENSG00000172985 | NA | NA | NA |
| 06042 | chr3:11821911-11821911 | + | 537 | no | TAMM41 | ENSG00000144559 | NA | NA | NA |
| 06042 | chr3:46012891-46012891 | - | 13 | no | NA | NA | NA | NA | NA |
| 06042 | chr3:46016949-46016949 | - | 12 | no | NA | NA | NA | NA | NA |
| 06042 | chr3:48869429-48869430 | + | 2368 | no | SLC25A20 | ENSG00000178537 | NA | NA | NA |
| 06042 | chr3:49064077-49064077 | - | 154 | yes | QRICH1 | ENSG00000198218 | NA | NA | NA |
| 06042 | chr3:51442102-51442102 | + | 1040 | no | DCAF1 | ENSG00000145041 | NA | TRUE | NA |
| 06042 | chr3:74911977-74911977 | + | 177 | no | NA | NA | NA | NA | NA |
| 06042 | chr5:55727289-55727289 | - | 6 | no | SLC38A9 | ENSG00000177058 | NA | NA | NA |
| 06042 | chr5:112902151-112902155 | + | 1509 | no | REEP5 | ENSG00000129625 | NA | NA | NA |
| 06042 | chr5:177645378-177645378 | + | 768 | no | AC139795.1 | ENSG00000246596 | NA | NA | NA |
| 06042 | chr6:26395114-26395114 | + | 426 | no | NA | NA | INTERNAL | NA | NA |
| 06042 | chr6:31467275-31467275 | + | 30 | no | HCP5 | ENSG00000206337 | NA | NA | NA |
| 06042 | chr6:54898539-54898539 | - | 5 | no | FAM83B | ENSG00000168143 | NA | NA | NA |
| 06042 | chr7:2952869-2952869 | - | 213 | no | CARD11 | ENSG00000198286 | NA | NA | NA |
| 06042 | chr7:56432642-56432643 | + | 452 | yes? | AC092447.7 | ENSG00000237268 | LTR3 or LTR5 | NA | NA |
| 06042 | chr7:100373718-100373718 | + | 945 | no | PILRA | ENSG00000085514 | NA | NA | NA |
| 06042 | chr7:144643923-144643924 | - | 1883 | yes | TPK1 | ENSG00000196511 | INTERNAL | NA | NA |
| 06042 | chr8:59999204-59999204 | - | 175 | no | NA | NA | LTR3 or LTR5 | NA | NA |
| 06042 | chr8:120945552-120945552 | + | 2078 | no | AC068413.1 | ENSG00000253619 | LTR3 or LTR5 | TRUE | NA |
| 06042 | chr9:127959223-127959223 | - | 73 | no | FAM102A | ENSG00000167106 | NA | NA | NA |
| 06042 | chr9:128590915-128590915 | + | 2 | no | SPTAN1 | ENSG00000197694 | NA | NA | NA |
| 06042 | chr9:128904247-128904247 | - | 971 | yes | LRRC8A | ENSG00000136802 | LTR3 | NA | NA |

| | | | | | | | | | | |
|---|---|---|---|---|---|---|---|---|---|---|
| 06042 | chr10:73367972-73367972 | + | 1161 | no | NA | NA | NA | NA | NA | |
| 06042 | chr10:74192227-74192227 | - | 2 | no | ADK | ENSG00000156110 | NA | NA | NA | |
| 06042 | chr10:89376486-89376488 | + | 1360 | no | LIPA | ENSG00000107798 | NA | NA | NA | |
| 06042 | chr11:4063466-4063485 | + | 2453 | no | STIM1 | ENSG00000167323 | NA | NA | NA | |
| 06042 | chr11:10861921-10861921 | + | 187 | no | ZBED5-AS1 | ENSG00000247271 | NA | NA | NA | |
| 06042 | chr11:34089807-34089807 | - | 1563 | no | CAPRIN1 | ENSG00000135387 | NA | NA | NA | |
| 06042 | chr11:62781235-62781236 | + | 3478 | no | TAF6L;TMEM223 | ENSG00000162227;ENSG00000168569 | NA | NA | NA | |
| 06042 | chr11:67159377-67159377 | + | 5 | no | KDM2A | ENSG00000173120 | NA | NA | NA | |
| 06042 | chr11:68307652-68307652 | - | 9 | no | NA | NA | NA | NA | NA | |
| 06042 | chr11:95681111-95681111 | - | 2492 | no | AP000820.2 | ENSG00000285842 | NA | NA | NA | |
| 06042 | chr12:54267066-54267066 | - | 74 | yes | CBX5;SCAT2 | ENSG00000094916;ENSG00000257596 | LTR3 or LTR5 | NA | NA | |
| 06042 | chr12:57354431-57354431 | + | 222 | no | R3HDM2 | ENSG00000179912 | NA | NA | NA | |
| 06042 | chr12:92914274-92914274 | - | 2 | no | EEA1 | ENSG00000102189 | NA | NA | NA | |
| 06042 | chr12:120056029-120056029 | - | 4 | no | BICDL1 | ENSG00000135127 | NA | NA | NA | |
| 06042 | chr14:17262496-17262496 | + | 98 | no | NA | NA | NA | NA | NA | Insertion point in centromeric repeat |
| 06042 | chr15:78893249-78893254 | - | 645 | no | MORF4L1 | ENSG00000185787 | NA | NA | NA | |
| 06042 | chr16:1691122-1691122 | - | 898 | no | JPT2 | ENSG00000206053 | NA | NA | NA | |
| 06042 | chr16:3540342-3540342 | + | 31 | no | NLRC3 | ENSG00000167984 | NA | NA | NA | |
| 06042 | chr16:14149364-14149364 | + | 19 | no | MRTFB | ENSG00000186260 | NA | NA | NA | |
| 06042 | chr16:14211240-14211240 | + | 1519 | no | MRTFB | ENSG00000186260 | LTR3 or LTR5 | NA | NA | |
| 06042 | chr16:21484467-21484468 | + | 4449 | no | SMG1P3 | ENSG00000180747 | NA | NA | NA | Insertion point in segmental duplication, also maps to: chr16:21915957-21915957 |
| 06042 | chr16:24995263-24995263 | - | 111 | no | ARHGAP17 | ENSG00000140750 | NA | NA | NA | |
| 06042 | chr16:50203656-50203657 | + | 2550 | no | TENT4B | ENSG00000121274 | NA | NA | NA | |
| 06042 | chr16:53662878-53662878 | - | 42 | no | RPGRIP1L | ENSG00000103494 | NA | NA | NA | |

| | | | | | | | | | |
|---|---|---|---|---|---|---|---|---|---|
| 06042 | chr16:88505647-88505647 | + | 46 | no | ZFPM1 | ENSG00000179588 | NA | NA | NA |
| 06042 | chr17:18029959-18029959 | + | 9 | no | ATPAF2 | ENSG00000171953 | NA | NA | NA |
| 06042 | chr17:28051014-28051014 | + | 8 | no | NLK | ENSG00000087095 | NA | NA | NA |
| 06042 | chr17:42253536-42253536 | - | 481 | no | STAT5B | ENSG00000173757 | NA | NA | NA |
| 06042 | chr17:42263154-42263154 | - | 113 | no | STAT5B | ENSG00000173757 | NA | NA | NA |
| 06042 | chr17:42265627-42265627 | - | 517 | no | STAT5B | ENSG00000173757 | LTR3 or LTR5 | NA | NA |
| 06042 | chr17:42267277-42267277 | - | 824 | no | STAT5B | ENSG00000173757 | NA | NA | NA |
| 06042 | chr17:49802734-49802734 | - | 573 | no | KAT7 | ENSG00000136504 | NA | NA | NA |
| 06042 | chr17:50753966-50753966 | - | 1165 | no | LUC7L3 | ENSG00000108848 | NA | NA | NA |
| 06042 | chr17:56846150-56846150 | + | 352 | no | TRIM25;DGKE | ENSG00000121060;ENSG00000153933 | NA | NA | NA |
| 06042 | chr17:76784687-76784687 | - | 2 | no | NA | NA | NA | NA | NA |
| 06042 | chr17:77413224-77413224 | + | 44 | no | SEPTIN9 | ENSG00000184640 | NA | NA | NA |
| 06042 | chr19:14098796-14098796 | - | 8 | no | PRKACA | ENSG00000072062 | NA | NA | NA |
| 06042 | chr19:15321839-15321840 | + | 311 | no | BRD4 | ENSG00000141867 | NA | TRUE | NA |
| 06042 | chr19:37329721-37329726 | - | 226 | no | ZNF875 | ENSG00000181666 | NA | NA | NA |
| 06042 | chr19:40297189-40297189 | - | 187 | no | NA | NA | NA | NA | NA |
| 06042 | chr19:40726270-40726271 | + | 6 | no | ITPKC | ENSG00000086544 | NA | NA | NA |
| 06042 | chr19:40858971-40858972 | + | 1255 | no | AC008537.1 | ENSG00000268797 | NA | NA | NA |
| 06042 | chr20:44540248-44540248 | + | 73 | no | PKIG | ENSG00000168734 | LTR3 or LTR5 | TRUE | NA |
| 06042 | chr22:30151237-30151238 | - | 811 | no | HORMAD2 | ENSG00000176635 | NA | NA | NA |
| 06042 | chr22:31255388-31255388 | + | 41 | no | LIMK2 | ENSG00000182541 | NA | NA | NA |
| 06042 | chrX:124043305-124043305 | + | 480 | no | STAG2 | ENSG00000101972 | NA | NA | NA |
| 06042 | chrX:124354469-124354469 | + | 18 | no | SH2D1A;STAG2 | ENSG00000183918;ENSG00000101972 | NA | NA | NA |

**Supplementary Table 5** HIV integration sites identified in patients 02006 and 06042

Approximate coverage at integration site was obtained via igvtools count command.
Integration sites were classified as clonal or not by manual inspection of the integration sites in IGV. If the reads showed evidence of more than one shear site they were classified as clonal.
DELETIONS referred to as “LTR3 or LTR5” are proviruses where it is not possible to definitively distinguish between the shear site and the actual deletion breakpoint in the provirus.

| # | Approximate location in genome (BTA6) | Provirus name | 10201e6 | Mannequin | 571 | Provirus |
|---|---|---|---|---|---|---|
| 1 | chr1:108,822,892-108,832,262 | BTA1_108.8 | no | no | YES | Full |
| 2 | chr1:140,473,236-140,486,732 | BTA1_140.4 | YES | no | YES | Full |
| 3 | chr2:7,341,443-7,349,776 | BTA2_7.3 | no | no | YES | Full |
| 4 | chr2:68,574,688-68,583,604 | BTA2_68.5 | YES | no | no | Partial |
| 5 | chr2:108,763,340-108,771,071 | BTA2_108.7 | no | YES | no | Full |
| 6 | chr2:136,856,893-136,860,100 | BTA2_136.8 | YES | no | no | Full |
| 7 | chr3:11,025,879-11,032,187 | BTA3_11.0 | no | YES | no | Full |
| 8 | chr3:21,243,379-21,247,173 | BTA3_21.24 | no | YES | no | Full |
| 9 | chr3:21,262,507-21,266,148 | BTA3_21.26 | no | YES | no | Full |
| 10 | chr3:115,305,677-115,313,191 | BTA3_115.3 | YES | no | no | **Full*** |
| 11 | chr4:23,529,679-23,538,398 | BTA4_23.5 | YES | no | no | Partial |
| 12 | chr4:106,804,424-106,812,368 | BTA4_106.8 | no | no | YES | Full |
| 13 | chr5:76,505,040-76,518,833 | BTA5_76.5 | YES | YES | YES | Full |
| 14 | chr6:19,795,982-19,804,772 | BTA6_19.7 | YES | YES | YES | Full |
| 15 | chr6:33,664,998-33,674,349 | BTA6_33.6 | YES | no | no | Full |
| 16 | chr6:93,979,584-93,984,028 | BTA6_93.9 | YES | YES | YES | Partial |
| 17 | chr7:18,507,208-18,514,234 | BTA7_18.5 | no | YES | no | Partial |
| 18 | chr7:62,318,935-62,329,558 | BTA7_62.3 | YES | no | no | Full |
| 19 | chr7:109,501,965-109,512,061 | BTA7_109.5 | YES | no | YES | Full |
| 20 | chr8:16,410,224-16,424,259 | BTA8_16.4 | YES | no | YES | Full |
| 21 | chr8:37,357,029-37,369,016 | BTA8_37.3 | no | YES | no | Full |
| 22 | chr8:67,963,331-67,972,754 | BTA8_67.9 | no | YES | no | Full |
| 23 | chr8:81,237,785-81,244,766 | BTA8_81.2 | YES | YES | no | Full |
| 24 | chr9:15,412,806-15,418,477 | BTA9_15.4 | YES | no | no | Partial |
| 25 | chr9:83,082,008-83,092,749 | BTA9_83.0 | YES | no | no | Full |
| 26 | chr9:84,257,434-84,262,548 | BTA9_84.2 | YES | no | no | Full |
| 27 | chr9:101,949,614-101,957,434 | BTA9_101.9 | YES | YES | no | Full |
| 28 | chr10:71,920,524-71,928,975 | BTA10_71.9 | YES | no | no | Full |
| 29 | chr10:87,425,735-87,443,841 | BTA10_87.4 | YES | YES | YES | Partial |
| 30 | chr11:50,592,847-50,606,524 | BTA11_50.5 | YES | no | YES | Full |
| 31 | chr11:61,788,705-61,792,024 | BTA11_61.7 | no | YES | no | Full |
| 32 | chr11:77,955,413-77,963,724 | BTA11_77.9 | YES | no | no | **Full#** |
| 33 | chr12:72,978,039-72,985,406 | BTA12_72.9 | YES | YES | no | Full |
| 34 | chr12:74,723,248-74,731,915 | BTA12_74.7 | YES | YES | no | Partial |
| 35 | chr15:9,435,764-9,439,369 | BTA15_9.4 | YES | YES | YES | Full |
| 36 | chr16:10,720,162-10,727,571 | BTA16_10.7 | YES | no | no | Full |
| 37 | chr16:13,308,596-13,315,659 | BTA16_13.3 | YES | no | no | Partial |
| 38 | chr16:28,504,653-28,536,456 | BTA16_28.5 | YES | no | YES | Full |
| 39 | chr18:27,619,893-27,626,348 | BTA18_27.6 | YES | no | YES | Partial |
| 40 | chr18:27,715,161-27,722,285 | BTA18_27.7 | no | no | YES | Full |
| 41 | chr18:50,368,602-50,378,304 | BTA18_50.3 | YES | YES | YES | Full |
| 42 | chr18:60,211,168-60,220,590 | BTA18_60.2 | YES | YES | YES | Partial |
| 43 | chr18:61,691,367-61,697,347 | BTA18_61.6 | YES | no | YES | Full |
| 44 | chr19:5,180,841-5,189,334 | BTA19_5.1 | YES | no | no | Partial |
| 45 | chr19:22,014,748-22,025,138 | BTA19_22.0 | YES | no | no | Full |
| 46 | chr19:51,039,969-51,101,363 | BTA19_51.0 | no | YES | YES | Partial |
| 47 | chr20:15,283,426-15,290,599 | BTA20_15.2 | YES | no | no | Full |
| 48 | chr20:55,126,259-55,134,120 | BTA20_55.1 | no | YES | no | Full |
| 49 | chr21:1,241,740-1,256,399 | BTA21_1.2 | YES | YES | YES | Partial |
| 50 | chr21:2,303,211-2,307,834 | BTA21_2.3 | no | YES | no | Full |
| 51 | chr21:4,133,180-4,142,631 | BTA21_4.1 | no | no | no | Full |
| 52 | chr21:18,634,068-18,645,042 | BTA21_18.6 | no | YES | no | Full |
| 53 | chr22:160,456-166,792 | BTA22_160.4 | no | no | YES | Full |
| 54 | chr23:41,312,657-41,328,100 | BTA23_41.3 | YES | no | no | Full |
| 55 | chr23:52,329,640-52,337,577 | BTA23_52.3 | YES | no | no | Full |
| 56 | chr24:12,819,683-12,824,449 | BTA24_12.6 | YES | YES | no | Partial |
| 57 | chr24:53,067,680-53,078,844 | BTA24_53.0 | no | no | YES | Full |
| 58 | chr25:20,428,960-20,444,963 | BTA25_20.4 | no | no | no | Full |
| 59 | chr26:50,606,858-50,616,960 | BTA26_50.6 | YES | no | no | Full |
| 60 | chr27:14,146,146-14,156,627 | BTA27_14.1 | no | YES | no | Full |
| 61 | chr28:17,575,320-17,582,731 | BTA28_17.5 | YES | no | no | Full |
| 62 | chr29:39,631,808-39,639,476 | BTA29_39.6 | YES | no | no | Full |
| 63 | chrX:27,723,875-27,732,458 | BTAX_27.7 | no | YES | no | Full |
| 64 | chrX:30,183,463-30,187,122 | BTAX_30.1 | YES | no | no | Partial |
| 65 | chrX:36,260,818-36,264,888 | BTAX_36.2 | YES | no | no | Partial |
| 66 | chrX:43,949,278-43,960,449 | BTAX_43.9 | no | no | YES | Full |
| 67 | chrX:47,314,044-47,327,526 | BTAX_47.3 | no | no | YES | Full |

**Supplementary Table 6** Endogenous retroviruses (BERVK2) identified in cattle via PCIP-seq.

*****LTR matches APOB ERV (BTA11_77.9)
#ERV inserted into APOB
Full = Full length ERV.
Partial = ERV with large deletion**.**

| | Approximate location in genome (OAR3) | ERV name | **220** | **221** | provirus |
|---|---|---|---|---|---|
| 1 | chr1:57,132,178-57,139,903 | OAR1_57.13 | no | YES | Full |
| 2 | chr1:86,065,652-86,091,348 | OAR1_86.0 | YES | YES | Full |
| 3 | chr1:129,489,883-129,502,056 | OAR1_129.4 | no | YES | Full |
| 4 | chr1:220,250,002-220,258,800 | OAR1_220.2 | YES | YES | Full |
| 5 | chr1:240,077,458-240,092,905 | OAR1_240.0 | YES | YES | Partial |
| 6 | chr1:253,739,233-253,756,582 | OAR1_253.7 | YES | YES | Partial |
| 7 | chr2:196,585,537-196,593,010 | OAR2_196.5 | YES | no | Full |
| 8 | chr3:39,261,134-39,285,428 | OAR3_39.2 | YES | YES | Full |
| 9 | chr3:39653898-39656987 | OAR3_39.6 | YES | YES | Partial |
| 10 | chr3:151,767,643-151,783,037 | OAR3_151.7 | YES | YES | Partial |
| 11 | chr3:182,538,937-182,555,692 | OAR3_182.5 | YES | no | Full |
| 12 | chr4:40,485,410-40,504,790 | OAR4_40.4 | YES | YES | Full |
| 13 | chr4:77,416,611-77,428,510 | OAR4_77.4 | YES | YES | Partial |
| 14 | chr5:7,744,521-7,756,178 | OAR5_7.74 | YES | YES | Partial |
| 15 | chr5:64,916,815-64,926,920 | OAR5_64.9 | YES | no | Partial |
| 16 | chr5:73,009,027-73,018,771 | OAR5_73.0 | YES | no | Full |
| 17 | chr6:5,400,881-5,410,594 | OAR6_5.4 | no | YES | Full |
| 18 | chr6:6,789,991-6,858,767 | OAR6_6.7 | YES | YES | Partial |
| 19 | chr6:26,968,086-26,977,558 | OAR6_26.9 | no | YES | Full |
| 20 | chr8:2,974,531-2,988,179 | OAR8_2.9 | YES | YES | Partial |
| 21 | chr8:49,483,598-49,499,241 | OAR8_49.4 | YES | YES | Partial |
| 22 | chr9:48,096,442-48,105,912 | OAR9_48.0 | no | YES | Full |
| 23 | chr9:89,743,769-89,752,495 | OAR9_89.7 | no | YES | Partial |
| 24 | chr10:70,892,072-70,919,960 | OAR10_70.8 | YES | no | Partial |
| 25 | chr11:32,085,050-32,095,786 | OAR11_32.0 | YES | YES | Full |
| 26 | chr13:5,676,353-5,686,765 | OAR13_5.6 | no | YES | Full |
| 27 | chr13:16,714,529-16,726,069 | OAR13_16.7 | YES | YES | Full |
| 28 | chr13:37,514,438-37,529,955 | OAR13_37.5 | YES | YES | Full |
| 29 | chr13:66022872-66031772 | OAR13_66.0 | YES | no | Full |
| 30 | chr14:13,811,039-13,844,103 | OAR14_13.8 | YES | YES | Partial |
| 31 | chr14:15,011,370-15,043,076 | OAR14_15.0 | YES | YES | Partial |
| 32 | chr14:56,232,971-56,236,157 | OAR14_56.2 | YES | YES | Full |
| 33 | chr14:57,491,683-57,503,056 | OAR14_57.4 | no | YES | Partial |
| 34 | chr14:57,605,121-57,623,737 | OAR14_57.6 | YES | YES | Partial |
| 35 | chr15:10,864,017-10,870,430 | OAR15_10.8 | no | YES | Full |
| 36 | chr17:48,876,178-48,887,208 | OAR17_48.8 | no | YES | Full |
| 37 | chr18:1,738,143-1,751,356 | OAR18_1.7 | no | YES | Partial |
| 38 | chr18:67,778,281-67,799,930 | OAR18_67.7 | YES | YES | Full |
| 39 | chr19:52,665,989-52,689,785 | OAR19_52.6 | YES | YES | Partial |
| 40 | chr20:433,819-443,901 | OAR20_0.4 | YES | no | Full |
| 41 | chr20:1,237,366-1,250,699 | OAR20_1.2 | no | YES | Partial |
| 42 | chr20:27,598,593-27,615,677 | OAR20_27.5 | no | YES | Full |
| 43 | chr21:6,694,384-6,709,701 | OAR21_6.6 | YES | no | Partial |
| 44 | chr22:46,781,990-46,790,196 | OAR22_46.7 | no | YES | Full |
| 45 | chr26:8,253,764-8,265,010 | OAR26_8.2 | no | YES | Full |
| 46 | chrX:3,690,949-3,701,009 | OARX_3.6 | YES | no | Full |
| 47 | chrX:62,939,566-62,949,333 | OARX_62.9 | YES | YES | Partial |
| 48 | chrX:78,127,416-78,132,398 | OARX_78.1 | YES | no | Partial |

**Supplementary Table 7** Endogenous retroviruses (enJSRV) identified in sheep via PCIP-seq.

Full = Full length ERV.
Partial = ERV with large deletion.

| Patient | ID | Estimated read count | Overlapping Gene | geneID | Notes |
|---|---|---|---|---|---|
| HPV18_PX | chr1:201993711-201993711 | 1 | RNPEP | ENSG00000176393 | |
| HPV18_PX | chr1:54070808-54070808 | 1 | TCEANC2 | ENSG00000116205 | |
| HPV18_PX | chr1:74339164-74339164 | 2 | FPGT-TNNI3K | ENSG00000259030 | |
| HPV18_PX | chr11:72988358-72988358 | 6 | FCHSD2 | ENSG00000137478 | |
| HPV18_PX | chr12:124528897-124528897 | 5 | NCOR2 | ENSG00000196498 | |
| HPV18_PX | chr12:62430096-62430096 | 3 | NA | NA | |
| HPV18_PX | chr12:88750111-88750111 | 2 | NA | NA | |
| HPV18_PX | chr13:32401471-32401471 | 1 | N4BP2L1 | ENSG00000139597 | |
| HPV18_PX | chr13:59883976-59883976 | 1 | DIAPH3 | ENSG00000139734 | |
| HPV18_PX | chr13:70017637-70017637 | 1 | KLHL1 | ENSG00000150361 | |
| HPV18_PX | chr13:96145444-96145444 | 1 | HS6ST3 | ENSG00000185352 | |
| HPV18_PX | chr16:35696743-35696743 | 4 | NA | NA | |
| HPV18_PX | chr16:46391666-46391666 | 15 | NA | NA | |
| HPV18_PX | chr16:60839237-60839237 | 3 | NA | NA | |
| HPV18_PX | chr17:50736162-50736162 | 1 | LUC7L3 | ENSG00000108848 | |
| HPV18_PX | chr17:71945217-71945217 | 1 | NA | NA | |
| HPV18_PX | chr18:33256597-33256597 | 2 | CCDC178 | ENSG00000166960 | |
| HPV18_PX | chr2:175176252-175176252 | 1 | NA | NA | |
| HPV18_PX | chr2:184979785-184979785 | 1 | NA | NA | |
| HPV18_PX | chr2:222973976-222973976 | 1 | NA | NA | |
| HPV18_PX | chr20:26724089-27697774 | 1 | NA | NA | Virus in satellite repeat |
| HPV18_PX | chr20:59882951-59882951 | 4 | SYCP2 | ENSG00000196074 | |
| HPV18_PX | chr21:31443081-31443081 | 5 | TIAM1 | ENSG00000156299 | |
| HPV18_PX | chr21:8210410-8210516 | 6 | FP671120.3 | ENSG00000280800 | |
| HPV18_PX | chr21:8225927-8228889 | 9 | FP671120.1 | ENSG00000278996 | |
| HPV18_PX | chr21:8393406-8393551 | 9 | FP236383.2 | ENSG00000280614 | |
| HPV18_PX | chr21:8437761-8437761 | 9 | FP236383.3 | ENSG00000281181 | |
| HPV18_PX | chr21:8453856-8454775 | 19 | NA | NA | |
| HPV18_PX | chr3:141177260-141177260 | 1 | NA | NA | |
| HPV18_PX | chr3:183646815-183646815 | 5 | KLHL24 | ENSG00000114796 | |
| HPV18_PX | chr3:52477576-52477615 | 67 | NISCH | ENSG00000010322 | |
| HPV18_PX | chr3:52491989-52492028 | 67 | NISCH | ENSG00000010322 | |
| HPV18_PX | chr3:52564151-52564190 | 75 | SMIM4 | ENSG00000168273 | |
| HPV18_PX | chr4:113196089-113196089 | 3 | ANK2 | ENSG00000145362 | |
| HPV18_PX | chr4:118149173-118149173 | 2 | NDST3 | ENSG00000164100 | |
| HPV18_PX | chr4:125160196-125160196 | 2 | NA | NA | |
| HPV18_PX | chr4:8361851-8361851 | 1 | NA | NA | |
| HPV18_PX | chr5:85159333-85159333 | 2 | NA | NA | |
| HPV18_PX | chr6:12217019-12217019 | 1 | NA | NA | |
| HPV18_PX | chr6:58604926-59721758 | 1 | NA | NA | Virus in satellite repeat |
| HPV18_PX | chr6:60995120-60995120 | 4 | NA | NA | |
| HPV18_PX | chr6:72218404-72218404 | 3 | RIMS1 | ENSG00000079841 | |
| HPV18_PX | chr6:7655460-7655460 | 6 | NA | NA | |

| | | | | |
|---|---|---|---|---|
| HPV18_PX | chr7:55353950-55353950 | 10 | NA | NA |
| HPV18_PX | chr7:63798384-63798384 | 3 | NA | NA |
| HPV18_PX | chr7:7812181-7812181 | 4 | AC007161.3 | ENSG00000283549 |
| HPV18_PX | chr7:98111088-98111088 | 1 | LMTK2 | ENSG00000164715 |
| HPV18_PX | chr8:119801685-119801685 | 13 | TAF2 | ENSG00000064313 |
| HPV18_PX | chr8:2564068-2564068 | 1 | NA | NA |
| HPV18_PX | chr8:93515097-93515097 | 1 | LINC00535 | ENSG00000246662 |
| HPV18_PX | chr8:9886409-9886409 | 2 | NA | NA |
| HPV18_PX | chr9:12503146-12503146 | 1 | NA | NA |
| HPV18_PX | chr9:128458663-128458663 | 1 | ODF2 | ENSG00000136811 |
| HPV18_PX | chrX:19414286-19414286 | 1 | MAP3K15 | ENSG00000180815 |
| HPV18_PX | chrX:41675298-41675299 | 1 | CASK | ENSG00000147044 |

| | | | | |
|---|---|---|---|---|
| HPV18_PY | chr5:37774016-37774016 | 2 | NA | NA |
| HPV18_PY | chr7:64329003-64329003 | 2 | ZNF736 | ENSG00000234444 |
| HPV18_PY | chr4:184039889-184039889 | 2 | NA | NA |
| HPV18_PY | chr18:108534-108534 | 2 | NA | NA |
| HPV18_PY | chr3:59699600-59699600 | 1 | NA | NA |
| HPV18_PY | chr4:90546531-90546531 | 1 | CCSER1 | ENSG00000184305 |
| HPV18_PY | chr5:146985347-146985347 | 1 | PPP2R2B | ENSG00000156475 |
| HPV18_PY | chr6:41200232-41200232 | 1 | TREML2 | ENSG00000112195 |
| HPV18_PY | chr6:113561576-113561576 | 1 | NA | NA |
| HPV18_PY | chr1:107169512-107169512 | 1 | NTNG1 | ENSG00000162631 |
| HPV18_PY | chr1:218361256-218361256 | 1 | TGFB2 | ENSG00000092969 |
| HPV18_PY | chr3:52563123-52563123 | 1 | SMIM4 | ENSG00000168273 |
| HPV18_PY | chr9:15686595-15686595 | 1 | CCDC171 | ENSG00000164989 |
| HPV18_PY | chr9:137787856-137787856 | 1 | AL590627.1 | ENSG00000255585 |
| HPV18_PY | chr10:6703026-6703026 | 1 | AL158210.2 | ENSG00000285743 |
| HPV18_PY | chr10:23788794-23788794 | 1 | KIAA1217 | ENSG00000120549 |
| HPV18_PY | chr10:91570894-91570894 | 1 | NA | NA |
| HPV18_PY | chr11:97096506-97096506 | 1 | NA | NA |
| HPV18_PY | chr19:35339090-35339090 | 1 | CD22 | ENSG00000012124 |

**Supplementary Table 8** HPV integration sites identified in patients HPV18_PX and HPV18_PY

Estimated read count refers to number of reads after PCR duplicates have been removed, see https://github.com/GIGA-AnimalGenomics-BLV/PCIP/blob/master/README.md
